# Lithocholic acid phenocopies rejuvenating and life-extending effects of calorie restriction

**DOI:** 10.1101/2023.12.07.570559

**Authors:** Qi Qu, Yan Chen, Yu Wang, Shating Long, Weiche Wang, Heng-Ye Yang, Mengqi Li, Xiao Tian, Xiaoyan Wei, Yan-Hui Liu, Shengrong Xu, Cixiong Zhang, Mingxia Zhu, Sin Man Lam, Jianfeng Wu, Baoding Zhang, Zhong-Zheng Zheng, Hai-Long Piao, Guanghou Shui, Xianming Deng, Chen-Song Zhang, Sheng-Cai Lin

**Author notes:** These authors contributed equally: Qi Qu, Yan Chen, Yu Wang.

## Abstract

Calorie restriction (CR) is a dietary intervention to promote health and longevity. CR causes various metabolic changes in both the production and circulation of metabolites; however, it remains unclear which of the changed metabolite(s) can account for the physiological benefits of CR. Through metabolomic analysis of metabolites undergoing abundance changes during CR and subsequent functional validation, we found that lithocholic acid (LCA) is the only metabolite that alone can recapitulate the effects of CR, including activation of AMPK and the rejuvenating effects of muscle regeneration, grip strength and running capacity in mice. Interestingly, LCA also activates AMPK and exerts life- and health-extending effects in *Caenorhabditis elegans* and *Drosophila melanogaster*, indicating that these animal models are able to transmit the signalling of LCA once administered. Knockout of AMPK abrogates LCA-induced phenotypes, in nematodes and flies, as well as in mice. Together, we have identified that administration of the CR-upregulated metabolite LCA alone can confer anti-ageing benefits to metazoans, in an AMPK-dependent manner.

---

Calorie restriction (CR) that reduces caloric intake without causing malnutrition has been recognised as a non-pharmacological dietary behaviour for improving health^1-3^. The benefits of CR on lifespan and healthspan have been tested and observed in mice and primates, in addition to yeast, nematodes and flies, indicative of a general connection of reduced food-intake to longevity^4, 5^. During CR, organisms undergo a series of metabolic changes or adaptations^6, 7^, leading to altered abundance of metabolites, which include free fatty acids, cholesterol, vitamins, short-chain organic acids and bile acids, among others^8–10^. Importantly, changes of some serum metabolites have been shown to deter age□associated metabolic changes by controlling the homeostasis of cellular proteins, oxidative damage and inflammation^11–14^. In humans, CR also yields systemic health benefits, which include improved age-related frailty and diseases such as central obesity, insulin resistance, muscle deterioration, dyslipidemia, and cancer, without inducing adverse effects on life quality in both population studies and randomised controlled trials^15, 16^. Consequentially, there have sprung numerous “anti-ageing diet” modalities^17^ such as intermittent fasting^18^, fasting-mimicking diets^19^, and ketogenic diets^20, 21^, although it has been shown that hypercaloric ketogenic diets may have adverse effects on lifespan^22^.

The AMP-activated protein kinase (AMPK), which is highly conserved across eukaryotes^23^ and is activated under CR^24^, has been shown to be a critical mediator of the beneficial effects of CR^25^. AMPK regulates a large number of signalling pathways known to retard ageing, such as the inhibition of the target of rapamycin complex 1 (TORC1)^26–28^, inhibition of forkhead box O proteins (FOXOs)^29–31^ that mimics the reduction of insulin/IGF-1 signalling (rIIS)^32^, elevation of NAD^+^ that activates sirtuins^33, 34^, inhibition of CREB-regulated transcriptional coactivators (CRTCs)^35, 36^, and induction of TFEB^37, 38^. Moreover, AMPK acts on a variety of anti-ageing-related cellular processes, including autophagy^39–41^, proteostasis^42–44^, mitochondrial biogenesis^45–47^, mitohormesis^48^, germline stemness and viability^49^, inflammation^50^, and neurodegeneration^51^. AMPK has thus been a target for identifying caloric restriction mimetics (CRMs)^52^. Known CRMs, including metformin^53^ and resveratrol^54^, can extend lifespan and healthspan through activating AMPK in multiple organisms. However, how the CR-mediated metabolic adaptations in the body signal to activate AMPK to maintain health and extend lifespan remains unknown, particularly concerning which specific metabolite(s) changed during CR is able to or responsible for the modulation of AMPK.

In this study, we reasoned that serum metabolites that undergo changes after CR might harbour the ability to arouse beneficial effects at the cellular and organismal levels. To that end, we subjected mice to CR for 4 months. We found that the serum prepared from the calorie-restricted mice was sufficient to activate AMPK in mouse embryonic fibroblasts (MEFs), HEK293T cells, primary hepatocytes and primary myocytes, as assessed by levels of the phosphorylation of AMPKα (p-AMPKα) and the substrate acetyl coenzyme A carboxylases (p-ACC or p-ACC1/2) (Fig. 1a-d). Moreover, perfusion of the mouse CR serum into ad libitum-fed mice also led to activation of AMPK in both the liver and muscle (Fig 1e, f), indicating that the CR serum suffices to activate AMPK at the organismal level. We also found that heat-treated CR serum retained the ability to activate AMPK in cultured cells, while dialysis abolished it (Fig. 1g, h). Results above suggested that heat-stable metabolites with low molecular weights present in the CR serum can confer the ability to activate AMPK.

**Fig. 1.**
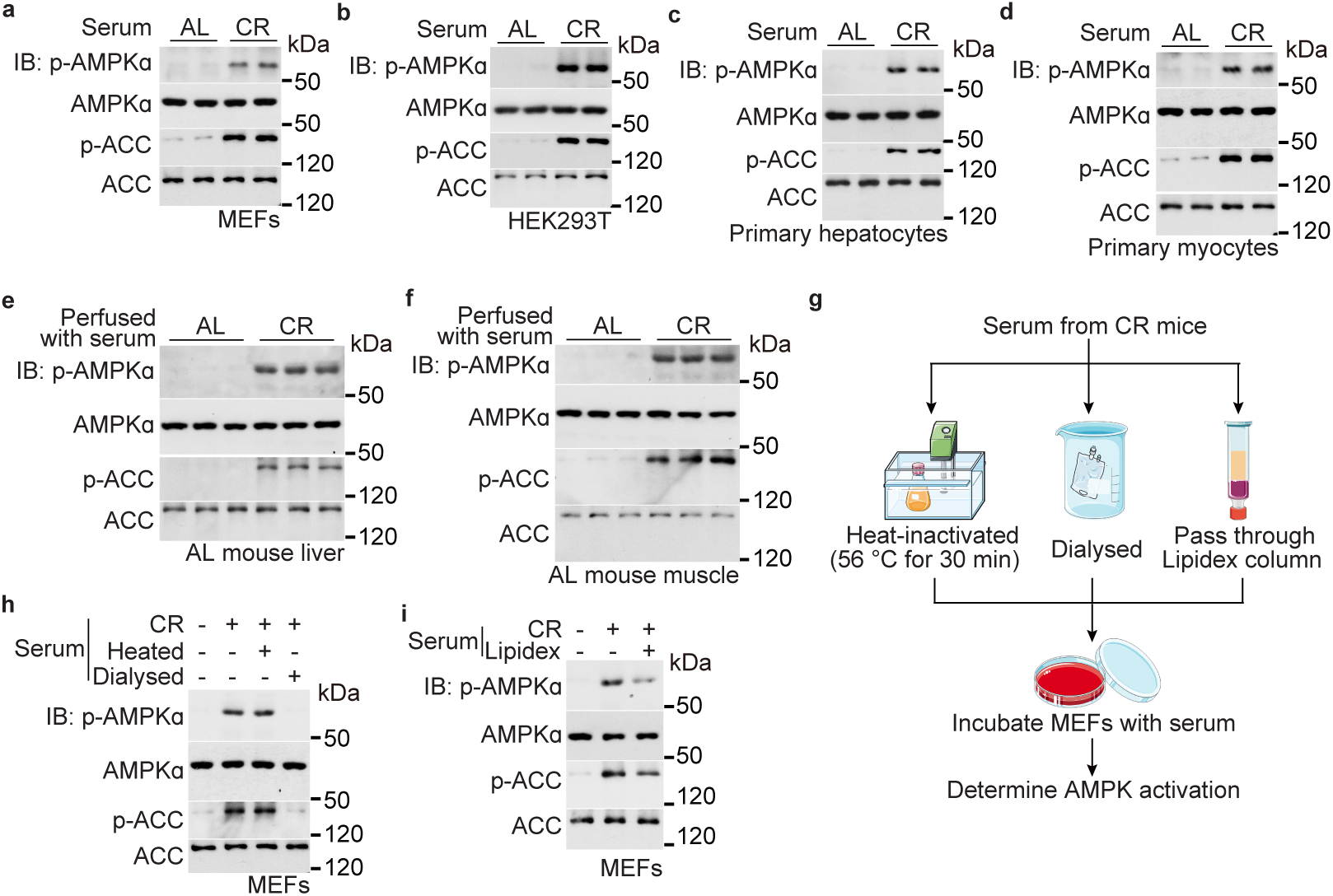
Serum from calorie-restricted mice confers AMPK activation to cells and perfused mice. **a**-**d**, Serum from calorie-restricted (CR) mice activates AMPK in cells cultured in a normal medium. MEFs (**a**), HEK293T cells (**b**), primary hepatocytes (**c**) and primary myocytes (**d**) were cultured respectively in DMEM (**a**, **b**), William’s medium E (**c**) and Ham’s F-10 medium (**d**), except that FBS (**a**, **b**, **d**) and BSA (**c**) supplemented in the medium was replaced with an equal volume of serum from mice calorie-restricted for 4 months (collected at 17:00, right before the next food supply; and same hereafter, unless stated otherwise) or from ad libitum fed (AL) mice as a control, for 4 h. Cells were then lysed, and the activation of AMPK was determined by immunoblotting for the levels of p-AMPKα and p-ACC. **e**, **f**, Serum from CR mice activates AMPK in the muscle and liver of the perfused mouse. Ad libitum-fed mice were jugularly perfused with 100 μl of serum from CR mice, or ad libitum-fed mice as a control, followed by determination for AMPK activation in liver and muscle tissues at 2 h after perfusion, by immunoblotting. **g**-**i**, Heat-stable, polar metabolites with low molecular weights in the CR serum mediate AMPK activation. MEFs were treated with CR serum as in **a**, except that the serum was heat-inactivated (**h**; at 56 °C for 30 min), dialysed (**h**), or passed through Lipidex column (**i**), followed by determination for AMPK activation (**h**, **i**; diagrammed in **g**). Experiments in this figure were performed three times, except **a**, **b** five times.

We next performed a series of metabolomics on the serum samples prepared from mice treated with or without CR. Different mass spectrometry (MS)-based approaches, including HPLC-MS, GC-MS, and CE-MS (all targeted), were applied to resolve small polar metabolites and nonpolar lipids. As summarised in Supplementary Table 1, a total of 1,215 metabolites (with 778 polar metabolites and 437 lipids) were hit, of which 695 metabolites (341 polar metabolites and 354 lipids) were significantly changed (*P* < 0.05) in the CR serum compared to control serum. Most prominent decreases were seen with long-chain fatty acids, phenylalanine and tyrosine, and increases with short-chain fatty acids, bile acids, and acyl-carnitine (Supplementary Table 1). After passing the CR serum through a lipophilic, Lipidex column that absorbs nonpolar compounds/lipids^55^, the pass-through fractions still retained the ability to activate AMPK as tested in MEFs (Fig. 1i), indicating that nonpolar compounds/lipids may not be involved in the AMPK activation. We hereafter focused on the polar metabolites changed after CR (212 increased and 129 decreased), and determined the ability of the individual metabolites to cause activation of AMPK in MEFs. As for the concentrations of the metabolites used in the screening assays, we referred to published studies. For metabolites with only ad libitum fed conditions reported, the concentrations used were adjusted by the observed fold changes after CR (see Extended Data Table 1). For metabolites with no concentrations reported, we set 10 mM as the highest concentration (Extended Data Table 1), given that serum metabolites of highest abundance lie below such a concentration^56^.

Among the 212 increased metabolites, 204 metabolites were tested for AMPK activation in MEFs, with the remaining 8 untested due to either failure of chemical synthesis or Prohibition by the law. As a result, a total of 6 metabolites were identified that could activate AMPK in the initial screening assays (Extended Data Table 1). For the metabolites that were decreased after CR, we tested whether they, 123 subtracted by 6 that were unavailable, could contribute to the activation of AMPK as a consequence of counteracting inhibition. It was found that none of the decreased metabolites showed an inhibitory effect on AMPK, as they at prior-to-CR serum concentrations all failed to attenuate or reverse the CR serum-induced AMPK activation in MEFs (Extended Data Table 1). Among the 6 AMPK-activating metabolites identified in the initial screening assays, LCA is the only metabolite that could activate AMPK at 1 μM in the culture medium, a concentration similar to that detected in the serum after CR, when tested in MEFs, HEK293T cells, primary hepatocytes and primary myocytes (Fig. 2b, c, Extended Data Fig. 1a, b). Of note, LCA was determined to be around 1.1 μM at 4 months of CR and remained constant in mice before and after feeding (Fig. 2a). As for LCA in ad libitum fed mice, low levels (0.3 μM) could be detected in the serum of postprandial mice, and then gradually decreased to undetectable levels at the fasting (postabsorptive) state (Extended Data Fig. 1c). Treatment of MEFs with LCA at 1 μM also led to an accumulation of intracellular LCA at approximately 0.8 μM (around 0.08 nmol/mg protein mass, equivalent to 0.8 μM after normalisation to the average cell volume and cell density reported previously^57, 58^), which is close to those detected in the tissues (i.e., around 0.03 nmol/mg protein mass in muscle, or 0.5 μM calculated according to the density of myocytes reported previously^59, 60^) of CR mice (Fig. 2a, b). Although the serum concentrations of LCA are similar between humans and mice^61, 62^, it is known that the synthesis and inter-conversion of bile acids in mice are different from humans, particularly muricholate that is abundant in mice, but hardly detectable in humans (reviewed in ref. ^63^). It was suggested that muricholate may interfere with the synthesis of other bile acids, potentially acting as a positive feedback regulator to increase LCA during CR^64^. We therefore analysed humanised mice that cannot synthesise muricholates due to the lack of the expression of the Cyp2c gene cluster^65^, and detected 0.8 μM in the serum of Cyp2c-null mice after CR (Fig. 2d), similar to those in wildtype mice, indicating that the increase of LCA by CR is unrelated to muricholates. As AMPK can be activated through several modes, we tested the AMP:ATP or ADP:ATP ratios^66, 67^ in cells treated with LCA, showing that LCA did not change the energy levels (Fig. 2e, f and Extended Data Fig. 1a, b), similar to that seen in the liver and muscle tissues of CR mice (Fig. 2g). We further show that unlike taurocholic acid, LCA did not depend on the cAMP-Epac-MEK pathway to activate AMPK^68^, as the treatment of MEK inhibitor PD98059 (ref. ^69^) failed to prevent the LCA-mediated AMPK activation (Extended Data Fig. 1d). In addition, LCA did not cause bulk Ca^2+^ increase that may lead to the CaMKK2-mediated AMPK activation^70–72^ (Fig. 2h), as assessed by the fluorescence intensities of Fluo-4-AM dye, consistent previous reports^73, 74^. We have further shown that the LCA derivatives, iso-LCA, 3-oxo-LCA, allo-LCA, isoallo-LCA and 3-oxo allo-LCA, did not activate AMPK in MEFs (Fig. 2i), indicating that LCA is a unique metabolite that activates AMPK.

**Fig. 2.**
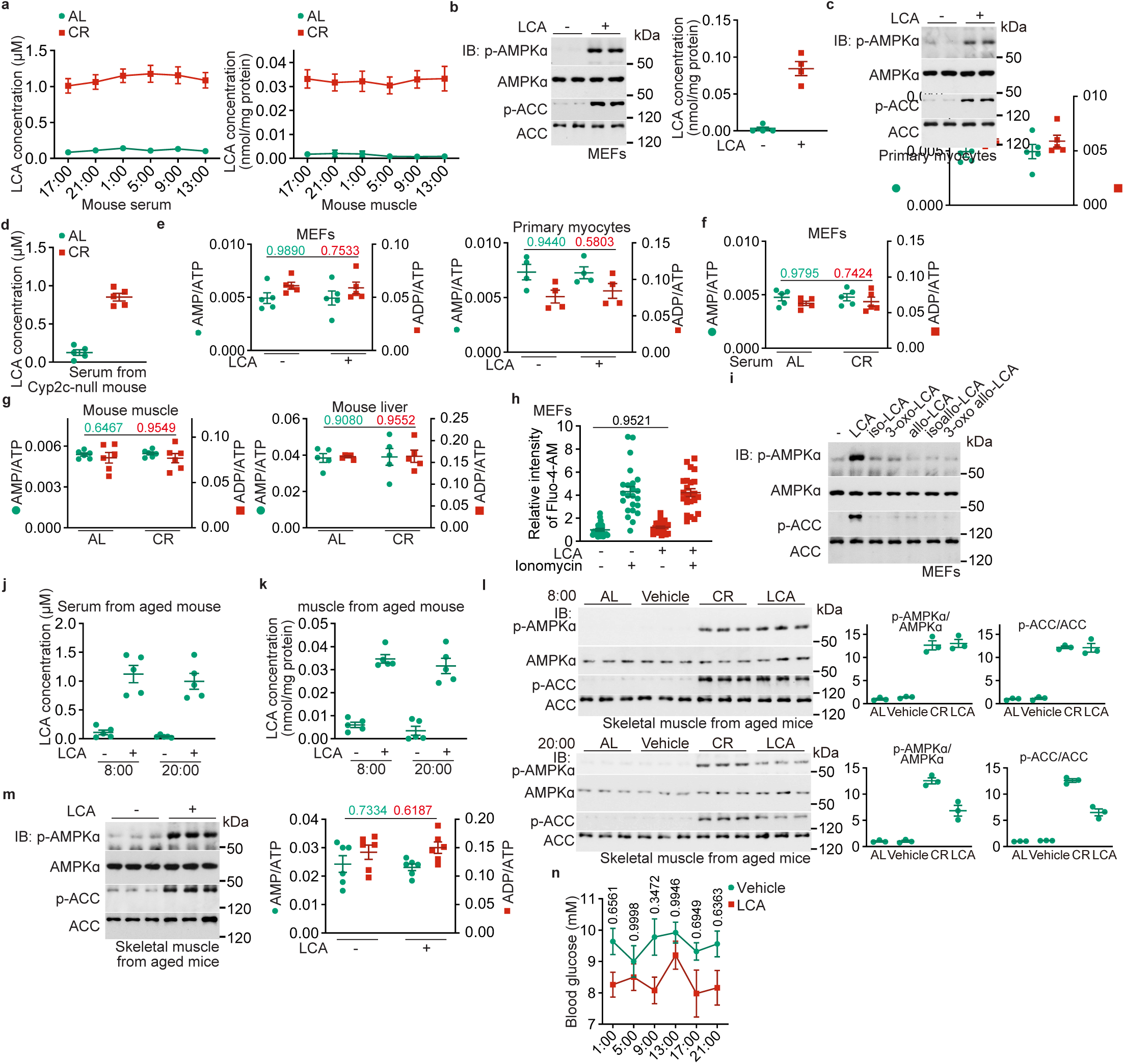
LCA is elevated after calorie restriction and responsible for AMPK activation. **a**, **d**, Metabolomic analysis reveals drastic elevation of LCA in the serum of CR mice. Serum and muscular concentrations of LCA at different times of the day from 4-month-calorie-restricted wildtype (**a**), the Cyp2c-null, or humanised mice (**d**) were determined. Results are shown as mean ± s.e.m.; *n* = 5 mice for each time point (**a**)/condition (**d**). **b**, **c**, LCA represents the AMPK-activating activity of CR serum. MEFs (**b**) or primary myocytes (**c**) were treated with 1 μM LCA, a concentration roughly equivalent to that in the serum of CR mice, for 4 h, followed by determination for the AMPK activity (left panel of **b**, and **c**) and the intracellular concentrations of LCA (right panel of **b**). See also Extended Data Fig. 1a, b for AMPK activation by LCA in HEK293T cells and primary hepatocytes. **e**-**g**, CR or LCA alone activates AMPK in an AMP-independent manner. The AMP:ADP and ADP:ATP ratios from MEFs (**e**, left panel) or primary myocytes (**e**, right panel) treated with 1 μM LCA (**e**) or serum from CR mice (**f**) for 4 h, or muscle and liver tissues from CR mice (**g**, collected at 17:00, the time before the food supply), were determined. Results are shown as mean ± s.e.m.; *n* = 4 (**e**, right panel), 6 (**g**, left panel) or 5 (others) biological replicates, and *P* value by two-sided Student’s *t*-test. **h**, LCA does not elevate intracellular calcium levels. The bulk calcium, as assessed by the intensities of the Fluo-4-AM dye, was determined in MEFs treated with 1 μM LCA for 4 h, or with 1 μM ionomycin for 5 min as a positive control. Results are shown as mean ± s.e.m., normalised to the group without LCA or ionomycin treatment; *n* = 22-23 cells, and *P* value by two-way ANOVA, followed by Tukey. **i**, LCA is a sole derivative of the bile acid, but not the other five, that can activate AMPK. MEFs were treated with 1 μM LCA, or 1 μM iso-LCA, 3-oxo-LCA, allo-LCA, isoallo-LCA or 3-oxoallo-LCA for 4 h, followed by determination of AMPK activities by immunoblotting. **j**, **k**, Administration of mice with LCA through drinking water leads to a similar accumulation of LCA to that by calorie restriction. Aged (1.5-year-old) mice were fed with (2-hydroxypropyl)-β-cyclodextrin-coated LCA at 1 g/l in their drinking water for 1 month, and the concentrations of LCA in the serum (**j**) and muscle tissue (**k**) of mice at two different times of the day (8:00, representing the light cycle, and 20:00, representing the dark cycle), were measured. Results are shown as mean ± s.e.m.; *n* = 5 mice for each condition. **l**, LCA administration activates AMPK in mice. Mice were subjected to CR as in **a**, or treated with LCA as in **j**, followed by determination of AMPK activities in muscle tissues at both light and dark cycles by immunoblotting. Results are shown as mean ± s.e.m., normalised to the AL group without LCA treatment; *n* = 3 mice for each condition. **m**, LCA administration does not elevate AMP in mice. The aged mice were treated as in **j**, followed by determining AMP:ADP and ADP:ATP ratios in muscle. Results are shown as mean ± s.e.m.; *n* = 6 mice for each condition, and *P* value by two-sided Student’s *t*-test. **n**, LCA decreases blood glucose as does CR. Levels of blood glucose at different times of the day in mice treated with LCA (as in **j**), for 1 month, were determined. Results are shown as mean ± s.e.m.; *n* = 5 mice for each treatment/time point, and *P* value by two-way ANOVA, followed by Tukey. Experiments in this figure were performed three times, except **b**, **j**, **k** four times.

We then determined the effects of LCA on the activation of AMPK in mouse tissues. Through testing different administration routes and titrating different doses in different formulations, we found that (2-hydroxypropyl)-β-cyclodextrin-coated LCA at 1 g/l in drinking water, led to an accumulation of approximately 1.1 μM LCA in the serum of aged (1-year-old) mice (Fig. 2j), similar to that of the LCA concentrations in the sera of CR-treated mice (Fig. 2a). Administration of LCA dissolved in drinking water also led to accumulation of muscular LCA levels to approximately 0.04 nmol/mg protein (approximately 0.4 μM), similar to that observed in CR muscle (Fig. 2k). Mice treated with LCA dissolved in drinking water robustly activated AMPK in skeletal muscles to a similar extent to that seen during CR (Fig. 2l, m). Consistent with the ability to activate AMPK^75^, LCA administration decreased blood glucose (Fig. 2n), mirroring the effects of CR on mouse blood glucose^3^. Together, LCA accounts for the ability of CR serum in AMPK activation.

We next determined the effects of LCA on ageing-related phenotypes. We found that in aged mice administered with LCA for a month, various features of muscle performance were improved. There was observed an increased number of oxidative muscle fibres (Fig. 3a; determined by the expression levels of MHCI and MHCIIa, markers for oxidative muscle fibres), a decreased number of glycolytic fibres (Fig. 3a; determined by the expression levels of MHCIIb), and reduced muscle atrophy (Fig. 3b; determined by the mRNA levels of *Trim63* and *Fbxo3*, see ref. ^76, 77^). Interestingly, LCA administration did not cause muscle loss (Fig. 3c; determined by the muscle weight and lean mass), a stark contrast to the decrease of muscle contents seen in calorie-restricted mice and humans^15, 78–80^. We also observed that LCA accelerated muscle regeneration after damage in aged mice (using the cardiotoxin) by promoting the induction of muscle stem cells (determined by the levels of PAX7; ref. ^81^) (Fig. 3d). LCA treatment increased NAD^+^ levels (Fig. 3e), elevated mitochondrial contents (assessed through morphology; the ratio of mitochondrial to nuclear DNA, or mtDNA:nDNA; and the expression of mitochondrial OXPHOS complex; Fig. 3f, g and Extended Data Fig. 2a, b) and the increase of mitochondrial respiratory function (oxygen consumption rates (OCRs); Fig. 3h) in muscle tissues of aged mice. Concurrently, significantly elevated energy expenditure (EE) was observed in these mice (Fig. 3i and Extended Data Fig. 2c, d). Running distance, duration and grip strength were also significantly increased in aged mice treated with LCA (Fig. 3j-l). In line with improved muscle functions, LCA treatment also ameliorated the age-associated glucose intolerance and insulin resistance ((determined by glucose tolerance test (GTT; Fig. 3m), insulin tolerance test (ITT; Fig. 3n), and the hyperinsulinemia euglycemic clamp (Fig. 3o)), without decreasing the rates of glucose production in these mice (Extended Data Fig. 3a-h). Knockout of *AMPK*α (both *AMPK*α*1* and *AMPK*α*2*) in mouse muscle dampened the effects of LCA in improving muscle functions (Fig. 4a-h and Extended Data Fig. 4a, b). We next tested for the possibility of LCA to extend lifespan by utilising *C*. *elegans* and *D*. *melanogaster* as it is known that CR can induce extension of lifespan in these two species^82, 83^. It is also known AMPK is required for lifespan extension in the same two species^29, 30, 84^. However, these organisms are incapable of synthesising LCA de novo. Nevertheless, the requirement of AMPK led us speculate that *C*. *elegans* and *D*. *melanogaster* may possess machineries that can transmit LCA signalling once administered. We therefore treated *C*. *elegans* and *D*. *melanogaster* with LCA as an exogenous stimulus, and determined whether AMPK could be activated in these animals. We found that LCA was as effectively absorbed into nematodes and flies as into mouse muscles (see Methods section for details of culture medium preparation), and activated AMPK in both nematodes and flies (Extended Data Fig. 5a-d). We also observed that LCA at these concentrations did not cause an increase in AMP (Extended Data Fig. 5a-d), suggesting that LCA indeed activates AMPK in nematodes and flies in a similar way to that in mice. Such administered LCA extended the mean lifespan of hermaphroditic nematodes from 22 to 27 days (Fig. 5a), male flies from 47 to 52 days, and female flies from 52 to 56 days (Fig. 5b and Extended Data Fig. 5e; similar to the effects of CR in the flies). LCA also significantly improved healthspan in nematodes and flies, as could be seen with significantly enhanced pharyngeal pumping rates in nematodes (Fig. 5c), improved oxidative stress resistance in both nematodes and flies (Fig. 5d-f and Extended Data Fig. 5f, g), better tolerance of cold, heat and starvation (food-deprivation) in flies (Fig. 5g-i and Extended Data Fig. 5h-j), higher NAD^+^ levels (Fig. 5j and Extended Data Fig. 5k), mtDNA:nDNA ratios (Fig. 5k and Extended Data Fig. 5l), mitochondrial genes (Fig. 5l and Extended Data Fig. 5m) in nematodes and flies, and higher OCRs in nematodes (Fig. 5m). We also found that knockout of AMPK in nematodes (by knocking out *aak*-*2*, the nematode orthologue for *AMPK*α*2*) and flies (knocking down *AMPK*α) abrogated all the anti-ageing effects of LCA (Fig. 5a-m), indicating that AMPK is required in the roles of LCA. Therefore, we have demonstrated that LCA accounts for the anti-ageing effects of CR in nematodes, flies and mice.

**Fig. 3.**
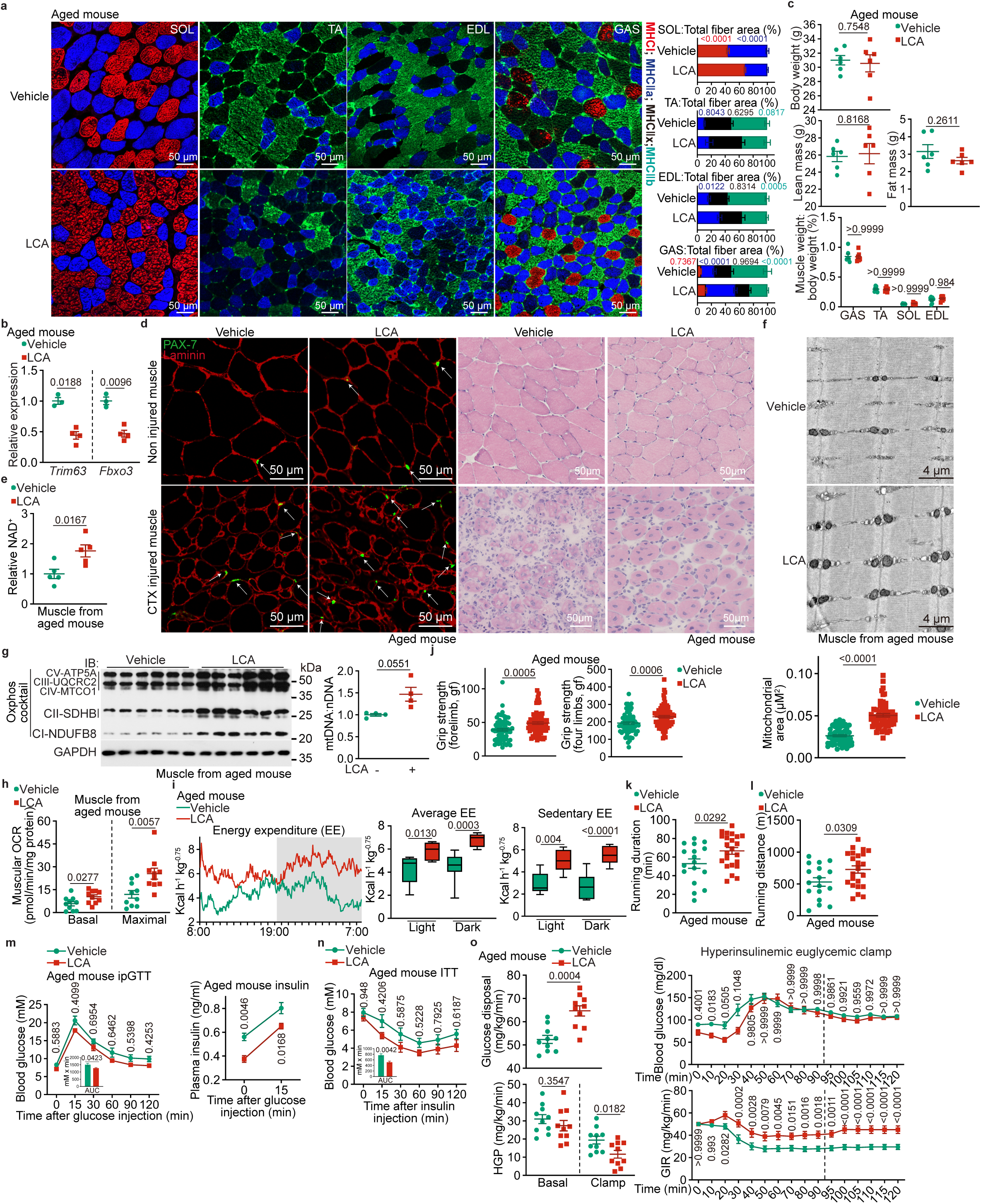
LCA exerts rejuvenating effects in mice. **a**, LCA induces oxidative fibre conversion in aged mice. Mice were fed with (2-hydroxypropyl)-β-cyclodextrin-coated LCA at 1 g/l in drinking water for 1 month, followed by determination of muscle fibre type by immunohistochemistry (see “histology” section of Methods for details). The representative images from whole-muscle cross sections for soleus (SOL), extensor digitorum longus (EDL), tibialis anterior (TA) and gastrocnemius (GAS) muscle are shown on the left panel, and the statistical analysis data right panel (mean ± s.e.m.; *n* = 6 mice for each condition, and *P* value by two-way ANOVA, followed by Tukey). **b**, **c**, LCA prevents muscle atrophy in aged mice. Mice were treated as in **a**, followed by determining the mRNA levels of atrophy markers *Trim63* and *Fbxo3* (**b**) and the body composition (the lean mass, fat mass and body weight, also the muscle mass; **c**). Results are shown as mean ± s.e.m.; *n* = 7 (muscle mass) or 6 (others) mice, and *P* value by two-way ANOVA, followed by Tukey (muscle mass) or by two-sided Student’s *t*-test (others). **d**, LCA accelerates muscle regeneration in damaged mice. Mice were treated with LCA as in **a**, and were intramuscularly injected with cardiotoxin to induce muscle damage. The morphology (right panel; by H&E staining) and the PAX7-positive muscle stem cells (left panel; by immunohistochemistry, and the white arrows indicate muscle stem cells) were used to determine muscle regeneration at 7 days after cardiotoxin injection. Representative images are shown in this figure. **e**, LCA elevates NAD^+^ levels in aged mice. Mice were treated with LCA as in **a**, followed by determination of muscular NAD^+^ levels. Results are shown as mean ± s.e.m., normalised to the vehicle group; *n* = 5 mice for each condition, and *P* value by two-sided Student’s *t*-test. **f**-**h**, LCA improves respiratory function in aged mouse muscles. Mice were treated with LCA as in **a**, followed by quantifying muscular mitochondrial contents (**f**; by TEM; representative images are shown on the upper panel, and statistical analysis data (the area of each mitochondrion in the section) on the lower panel (mean ± s.e.m.; *n* = 57 (vehicle) or 56 (LCA) mitochondria for each condition, and *P* value by two-sided Student’s *t*-test)), the protein levels of muscular OXPHOS complexes (left panel of **g**; by immunoblotting), mtDNA:nDNA ratios (right panel of **g**; by RT-PCR; results are shown on the right panel as mean ± s.e.m., normalised to the vehicle group; *n* = 4 mice for each condition, and *P* value by two-sided Student’s *t*-test), and the OCR in muscles (**h**; by the Seahorse Mito Stress Test; results are mean ± s.e.m.; *n* = 10 mice for each condition, and *P* value by two-sided Student’s *t*-test). **i**, LCA elevates energy expenditure (EE) in aged mice. Mice were treated with LCA as in **a**, followed by determining EE. Data are shown as mean (left panel; at 5-min intervals during a 24-h course after normalisation to the body weight (kg^0.75^)), or as box-and-whisker plots (middle panel, in which the lower and upper bounds of the box represent the first and the third quartile scores, the centre line represents the median, and the lower and upper limits denote minimum and maximum scores, respectively; and the same hereafter for all box-and-whisker plots; *P* value by two-way ANOVA, followed by Tukey), *n* = 6 (vehicle, light cycle of sedentary EE), 7 (LCA, dark cycle of sedentary EE), or 8 (others) mice for each condition. See also Extended Data Fig. 2c, d for the respiratory quotient (RQ) and the ambulatory activity data generated in this experiment. **j**-**l**, LCA promotes muscle strength and endurance in aged mice. Mice were treated with LCA as in **a**, followed by determining the grip strength (**j**, determined for the forelimb and four limbs separately, and results are mean ± s.e.m.; *n* = 65 (vehicle) or 75 (LCA) mice for each condition, and *P* value by two-sided Student’s *t*-test), running duration (**k**, results are mean ± s.e.m.; *n* = 17 (vehicle) or 23 (LCA) mice for each condition, and *P* value by two-sided Student’s *t*-test) and maximal running distance (**l**, results are mean ± s.e.m.; calculated according to **k**). **m**-**o**, LCA ameliorates age-associated insulin resistance in mice. Aged mice were treated as in **a**, followed by performing ipGTT (**m**, results of blood glucose and area under the curve (AUC) are shown as mean ± s.e.m.; *n* = 5 (vehicle) or 6 (LCA) mice for each condition, and *P* value by two-way repeated-measures (RM) ANOVA followed by Sidak’s test (blood glucose), or by two-sided Student’s *t*-test (AUC); and the serum insulin levels during ipGTT are shown as an inset, results are mean ± s.e.m.; *n* = 4 mice and *P* value by two-way RM ANOVA followed by Sidak’s test), ITT (**n**, results of blood glucose and AUC are shown as mean ± s.e.m.; *n* = 5 (vehicle) or 6 (LCA) mice for each condition, and *P* value by two-way RM ANOVA followed by Sidak’s test (blood glucose), or two-sided Student’s *t*-test (AUC)), and the hyperinsulinemia euglycemic clamp (**o**, the blood glucose levels and the GIR values during the clamp are shown on the right panel as mean ± s.e.m.; *n* = 10 mice; *P* value by two-way RM ANOVA followed by Sidak’s test; and the glucose disposal rates and the HGP rates, calculated according to the average values GIR during the steady-state (90-120 min, indicated by a dashed line), on the left panel as mean ± s.e.m.; *P* value by two-sided Student’s *t*-test). Experiments in this figure were performed three times.

**Fig. 4.**
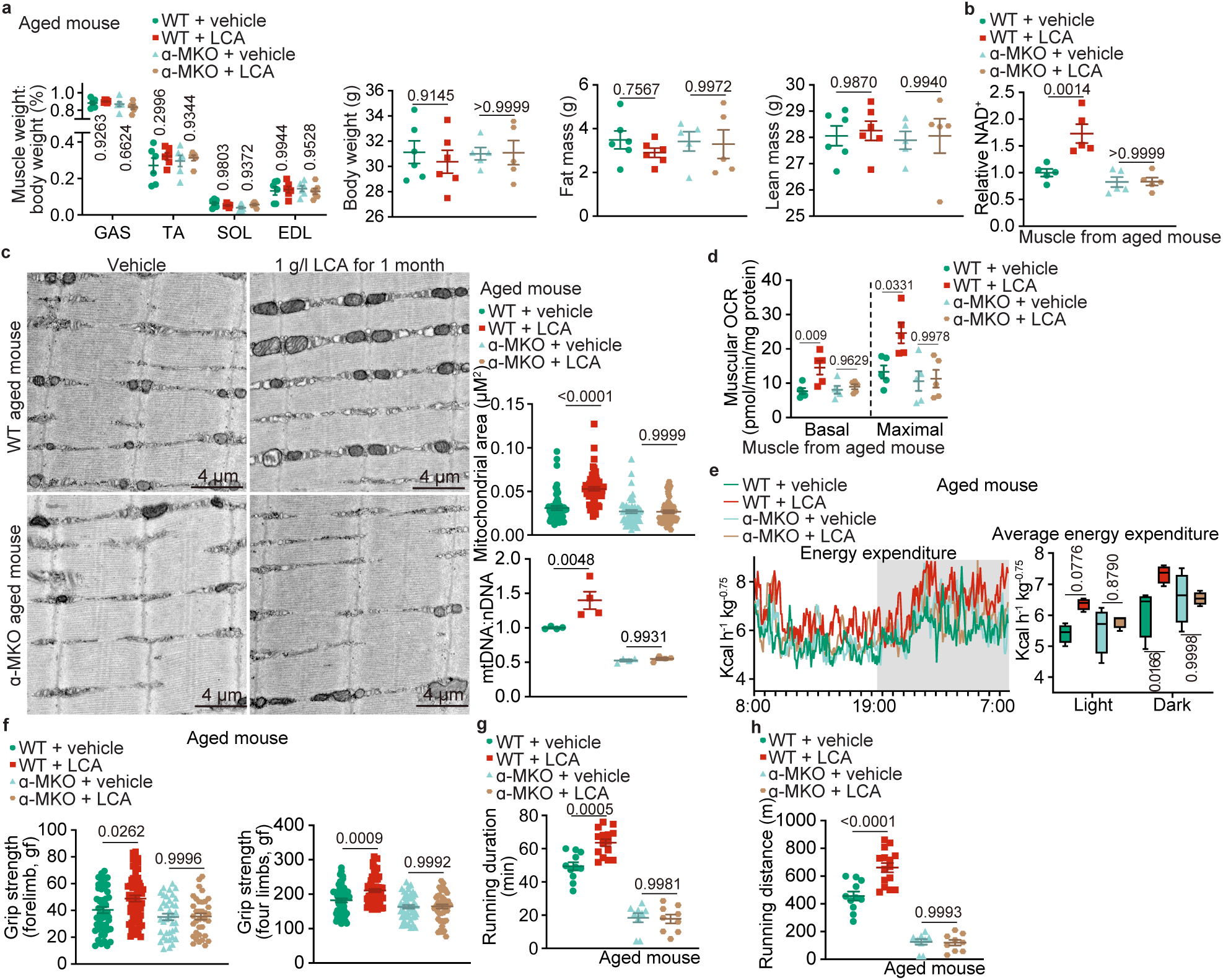
LCA-mediated rejuvenation depends on AMPK. **a**, **f**-**h**, Muscle-specific AMPK knockout abolishes the effects of LCA on muscle strength and endurance. The aged, *AMPK*α-MKO (α-MKO) mice and their wildtype (WT) littermates were fed with (2-hydroxypropyl)-β-cyclodextrin-coated LCA at 1 g/l in drinking water for 1 month, followed by determining muscle mass (also the body composition; **a**; results are mean ± s.e.m.; *n* = 5 (body composition of α-MKO) or 6 (others) mice for each genotype/treatment, and *P* value by two-way ANOVA followed by Tukey’s test), grip strength (**f**, determined for the forelimb and four limbs separately, and results are mean ± s.e.m.; *n* = 50 (WT, vehicle, forelimb), 61 (WT, LCA, forelimb), 38 (α-MKO, vehicle, forelimb), 36 (α-MKO, LCA, forelimb), 54 (WT, vehicle, four limbs), 65 (WT, LCA, four limbs) or 45 (others) mice for each condition, and *P* value by two-way ANOVA followed by Tukey’s test), running duration (**g**, results are mean ± s.e.m.; *n* = 11 (WT, vehicle), 15 (WT, LCA) or 9 (others) mice for each condition, and *P* value by two-way ANOVA followed by Tukey’s test) and maximal running distance (**h**, results are mean ± s.e.m.; calculated according to **g**). **b**, AMPK is required for the elevation of muscular NAD^+^ by LCA. Mice were treated as in **a**, followed by determination of muscular NAD^+^ levels. Results are shown as mean ± s.e.m., normalised to the WT, vehicle group; *n* = 5 mice for each genotype/treatment, and *P* value by two-way ANOVA followed by Tukey’s test. **c**, **d**, LCA improves mitochondrial respiratory function in aged mouse muscles in an AMPK-dependent manner. Mice were treated as in **a**, followed by determining the mitochondrial area in the section (**c**; representative images are shown on the left panel, and statistical analysis data on the right panel (mean ± s.e.m.; *n* = 53 (WT, vehicle), 62 (WT, LCA), 58 (α-MKO, vehicle) or 57 (α-MKO, LCA) mitochondria for each genotype/treatment, and *P* value by two-way ANOVA followed by Tukey’s test)), the mtDNA:nDNA ratios (right panel of **c**; results are shown on the right panel as mean ± s.e.m., normalised to the WT vehicle group; *n* = 4 mice for each genotype/treatment, and *P* value by two-way ANOVA followed by Tukey’s test), and the muscular OCR (**d**; results are shown on the right panel as mean ± s.e.m.; *n* = 5 mice for each genotype/treatment, and *P* value by two-way ANOVA followed by Tukey’s test). **e**, LCA elevates EE in aged mice depending on AMPK. Mice were treated as in **a**, followed by determining EE. Data are shown as mean (left panel; at 5-min intervals during a 24-h course after normalisation to the body weight (kg^0.75^); mean), or as box-and-whisker plots (right panel; *P* value by two-way ANOVA followed by Tukey’s test), *n* = 4 mice for each genotype/treatment. See also Extended Data Fig. 4b for the respiratory quotient (RQ) and the ambulatory activity data generated in this experiment. Experiments in this figure were performed three times.

**Fig. 5.**
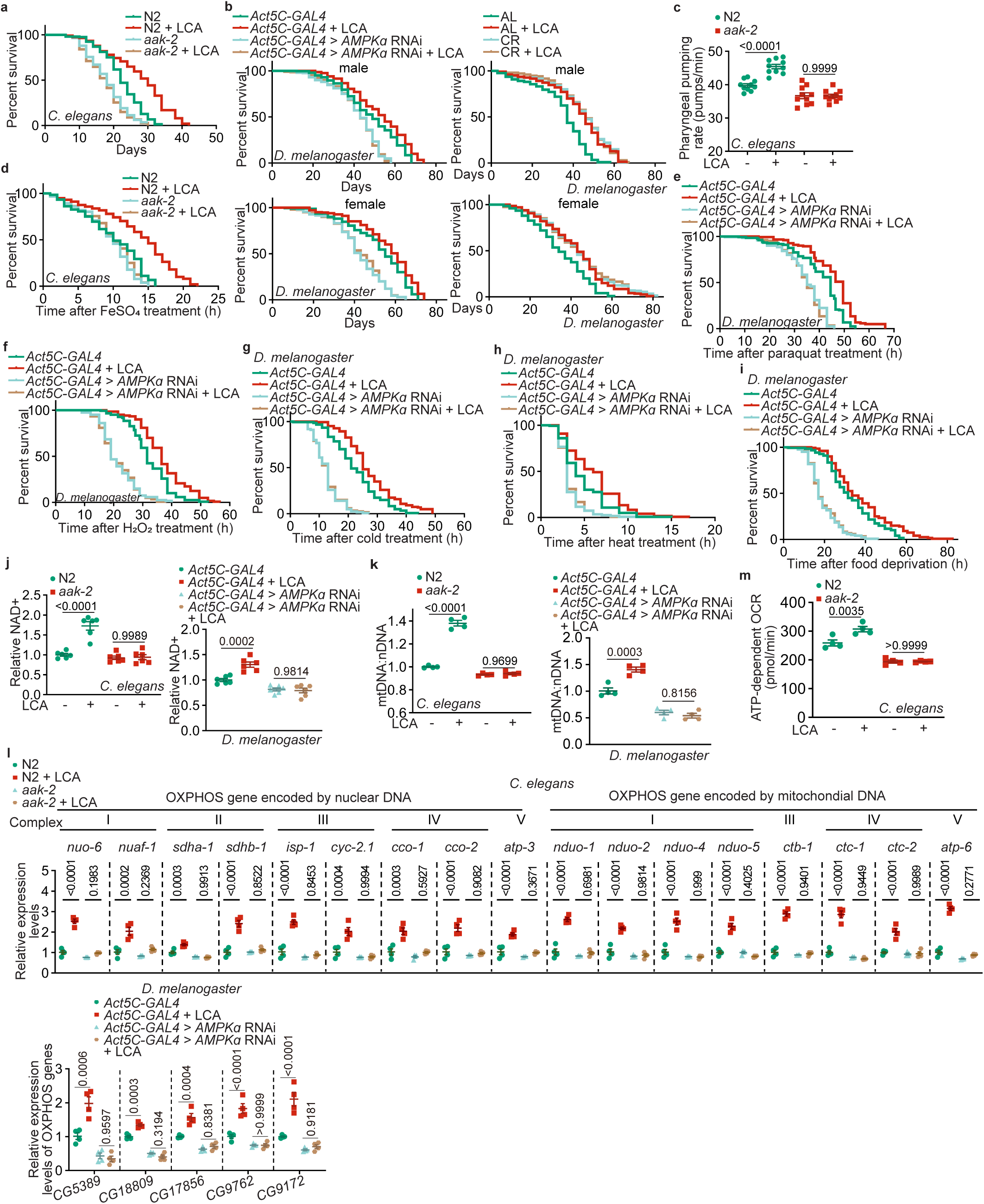
LCA extends lifespan and healthspan through AMPK. **a**, **b**, LCA extends lifespan in nematodes and flies through AMPK. Wildtype (**a**; N2) and *AMPK*α knockout (**a**; *aak*-*2*) nematodes, or the control (**b**; *Act5C*-*GAL4*) and the *AMPK*α knockdown (*Act5C*-*GAL4* > *AMPK*α RNAi) flies were cultured in medium containing LCA at 100 μM, which led to an activation of AMPK in a similar way to that in mice (**a**, and left panel of **b**; see validation data in Extended Data Fig 5a-d), or subjected to CR (right panel of **b**). Lifespan data are shown as Kaplan-Meier curves (see also statistical analyses data on Extended Data Table 2, and the same hereafter for all lifespan data). See also lifespan analysis on wildtype flies (*w^1118^*) in Extended Data Fig. 5e. **c**, LCA promotes nematode pharyngeal pumping rates in an AMPK-dependent manner. Wildtype and *AMPK*α knockout nematodes were treated with LCA for a day, followed by determining pharyngeal pumping rates. Results are mean ± s.e.m.; *n* = 10 worms for each genotype/treatment, and *P* value by two-way ANOVA followed by Tukey’s test). **d**-**f**, LCA promotes oxidative stress resistance of nematodes and flies via AMPK. Wildtype and *AMPK*α knockout nematodes (**d**), the control and the *AMPK*α knockdown flies (**e**, **f**), were treated with LCA for 2 days (**d**) or 30 days (**e**, **f**), followed by transfer to media containing 15 mM FeSO_4_ (**d**), 20 mM paraquat (**e**) or 5% H_2_O_2_ (**f**) to elicit oxidative stress. Lifespan data are shown as Kaplan-Meier curves. See also oxidative stress resistance analysis on wildtype flies in Extended Data Fig. 5f, g. **g**-**i**, LCA improved cold, heat and starvation resistance in flies through AMPK. the control and the *AMPK*α knockdown flies were treated with LCA as in **e**, followed by transferring to cold (4 °C; **g**), heat (37 °C; **h**) or food deprivation (**i**) conditions. Lifespan data are shown as Kaplan-Meier curves. See also cold, heat and starvation resistance analysis on wildtype flies in Extended Data Fig. 5h-j. **j**, LCA elevates NAD^+^ levels in nematodes and flies in an AMPK-dependent manner. Wildtype and *AMPK*α knockout nematodes (left panel), or the control and the *AMPK*α knockdown flies (right panel), were treated with LCA as in **d** (left panel) or **e** (right panel), followed by determining NAD^+^ levels. Results are mean ± s.e.m., normalised to the WT/control vehicle group; *n* = 6 (nematode) or 6 (fly) samples for each genotype/condition, and *P* value by two-way ANOVA followed by Tukey’s test. See also NAD^+^ levels of wildtype flies after LCA treatment in Extended Data Fig. 5h-k. **k**-**m**, LCA elevates mitochondrial contents and improves mitochondrial functions in nematodes and flies depending on AMPK. Wildtype and *AMPK*α knockout nematodes (left panels of **k** and **m**, and upper panel of **l**), or the control and the *AMPK*α knockdown flies (right panel of **k**, and lower panel of **l**), were treated with LCA as in **d** (left panels of **k** and **m**, and upper panel of **l**) and **e** (right panel of **k**, and lower panel of **l**), followed by determining the ratios of mtDNA:nDNA (**k**), the mRNA levels of mitochondrial OXPHOS complexes (**l**), and OCR (**m**). Results are mean ± s.e.m., normalised to the WT/control vehicle group; *n* = 4 samples for each genotype/treatment, and *P* value by two-way ANOVA followed by Tukey’s test. See also mitochondrial content and function analysis wildtype flies in Extended Data Fig. 5k-m. Experiments in this figure were performed three times.

## Discussion

In this current study, we have demonstrated that LCA is constantly present in dramatically higher concentrations in calorie-restricted mice, especially compared to the post-absorptive state of ad libitum fed mice. Importantly, the LCA was also found to be among the elevated metabolites in humans in some clinical trials^85^. As a secondary metabolite of bile acid synthesised in the liver, LCA is produced from precursors cholic acid (CA) and chenodeoxycholic acid (CDCA). These precursors are secreted from the liver into the intestine, where they are converted to LCA by gut microbiome, specifically the species of *Lactobacillus*^86^, *Clostridium*^87^ and *Eubacterium*^88^. These species express bile salt hydrolase and the 7α-dehydroxylase enzymes, which are sequentially responsible for converting CA and CDCA to LCA^89^. Given that all of the three genera were found to be increased after CR^90, 91^, it is reasonable to suggest that elevation of LCA during CR may be caused by the changes of these gut microbiome. Indeed, we have observed significantly higher concentrations of LCA in the intestine of CR mice (Extended Data Fig. 5n). Consistently, it has been shown that LCA, along with derivatives, is found at higher levels in healthy centenarians who carried high faecal levels of *Clostridioides*^92^.

We have provided multiple lines of evidence to show that LCA acts as a calorie restriction mimetic (CRM), which recapitulate the effects of CR, including the activation of AMPK, the rejuvenating effects and anti-ageing effects. First of all, LCA is the only metabolite that can activate AMPK at around 1 μM, which is similar to that detected in the serum of CR mice, and is way below the concentrations (i.e., hundreds of micromoles) that would cause cytotoxic effects^93, 94^. Although we have also found that at supra-high concentrations, other metabolites such as ferulic acid, 4-methyl-2-oxovaleric acid, 1-methyladenosine, methylmalonic acid and mandelic acid were able to activate AMPK (seen in our initial screening assays, as listed in Extended Data Table 1), the concentrations required for such AMPK activation were too high to be physiologically relevant. In addition, the series of derivatives of LCA did not activate AMPK. Second, LCA elevates the mitochondrial contents in the muscles of aged mice, which is known to trigger the glycolytic-oxidative fibre type transition^95, 96^, preserving the muscle force and endurance. Moreover, LCA enhances the ability of muscles to regenerate, a hallmark of rejuvenation^97^, likely through elevating muscular NAD^+^ levels, which is a downstream event of AMPK activation and is shown as a critical factor for mitochondrial biogenesis^33, 98^. The AMPK-dependent improvement of muscular mitochondrial function also helps ameliorate age-associated insulin resistance that may be caused by the increased inflammation and oxidative stress due to impaired mitochondrial function^99,100^. Third, we have also demonstrated that AMPK activation is required for LCA to extend lifespan, as assessed in nematodes and flies. Although one may argue that nematodes or flies do not synthesise LCA de novo, our findings are consistent with the reports that CR can induce AMPK activation and lifespan extension in nematodes^29, 30^ and flies^84^. It is hence reasonable to speculate that these two animal models possess similar downstream machineries that can transmit LCA signalling to the activation of AMPK for extension of lifespan. Indeed, as we have explored in the accompanying paper, the molecular mechanism that mediates LCA to activate AMPK is through intersecting the conserved lysosomal AMPK pathway via the v-ATPase, a critical node for AMPK activation after sensing low glucose^101–104^ or by metformin^105–107^.

An adverse effect of CR is muscle loss, possibly due to a prolonged need of supplementation for amino acids from muscle mass during calorie restriction^15, 80^. We found that administration of LCA alone to ad libitum fed mice avoids such an effect. Thus, LCA, as a natural body metabolite, will help overcome the adverse effect of muscle loss and provide a better means to improve healthspan in humans practising CR. Our observation that LCA alone can confer anti-ageing benefits testifies that CR can induce metabolites to exert a multitude of functions to ultimately prolong lifespan. Together, we have identified LCA as a CR-induced metabolite that can phenocopy the effects of CR in an AMPK-dependent manner.

## Supporting information

A complete list of the changes in serum metabolites during CR

Source data and statistical analysis data for all graphs

Uncropped gel images

## Online content

Any methods, additional references, Nature Portfolio reporting summaries, source data, extended data, supplementary information, acknowledgements, details of author contributions and competing interests; and statements of data and code availability are available at https://doi.org/10.1038/.

**Publisher’s note** Springer Nature remains neutral with regard to jurisdictional claims in published maps and institutional affiliations.

## Methods

### Data reporting

The chosen sample sizes were similar to those used in this field: *n* = 4-12 samples were used to evaluate the levels of metabolites in serum^10, 108^, cells^101, 104^, tissues^11, 57, 101, 104^, nematodes^109–111^ and flies^112–114^; *n* = 4-10 samples to determine OCR in tissues^104, 115^ and nematodes^116–118^; *n* = 3-4 samples to determine the mRNA levels of a specific gene^119^; *n* = 2-6 samples to determine the expression levels and phosphorylation levels of a specific protein^119^; *n* = 200 worms to determine lifespan^120–122^; *n* = 60 worms to determine healthspan^123–125^, except *n* = 10 worms for pharyngeal pumping rates^104^; *n* = 200 flies, male or female, to determine lifespan^126–128^; *n* = 60 flies, male or female, to determine healthspan^129–131^; *n* = 4-8 mice for EE and RQ^104^; *n* = 10 mice for hyperinsulinemic-euglycemic clamping^104, 132^; *n* = 5-6 mice for GTT and ITT^104^; *n* = 6 mice for body composition^104^; *n* = 6 mice for muscle fibre type^79, 133, 134^; *n* = 3 mice for muscle regeneration^125, 135, 136^; *n* = 53-62 mitochondria from 3 mice for muscular mitochondrial content^137, 138^; *n* = 9-23 mice for running duration^102, 104^; and *n* = 36-75 mice for grasp strength^104^. No statistical methods were used to predetermine sample size. All experimental findings were repeated as stated in figure legends, and all additional replication attempts were successful. For animal experiments, mice, nematodes and flies were housed under the same condition or place. For cell experiments, cells of each genotype were cultured in the same CO_2_ incubator and were parallel seeded. Each experiment was designed and performed along with proper controls, and samples for comparison were collected and analysed under the same conditions. Randomisation was applied wherever possible. For example, during MS analyses, samples were processed and subjected to MS in random orders. For animal experiments, sex-matched (for mice and flies), age-matched litter-mate animals in each genotype were randomly assigned to LCA or vehicle treatments. In cell experiments, cells of each genotype were seeded in parallel and randomly assigned to different treatments. Otherwise, randomisation was not performed. For example, when performing immunoblotting, samples needed to be loaded in a specific order to generate the final figures. Blinding was applied wherever possible. For example, samples, cages or agar plates/vials during sample collection and processing were labelled as code names that were later revealed by the individual who picked and treated animals or cells, but did not participate in sample collection and processing, until assessing outcome. Similarly, during microscopy data collection and statistical analyses, the fields of view were chosen on a random basis, and are often performed by different operators, preventing potentially biased selection for desired phenotypes. Otherwise, blinding was not performed, such as the measurement of OCR, as different reagents were added for particular reactions.

### Mouse strains

Wildtype C57BL/6J mice (#000664) were obtained from The Jackson Laboratory. *AXIN*^F/F^ and *LAMTOR1*^F/F^ mice were generated and validated as described previously^119^. *AMPK*α*1*^F/F^ (#014141) and *AMPK*α*2*^F/F^ mice (#014142) were obtained from Jackson Laboratory, provided by Dr. Sean Morrison. *AMPK*α-MKO mice were generated by crossing *AMPK*α*1*/*2*^F/F^ mice with *Mck*-*Cre* mice, as described and validated previously^104^. *Cyp2c* cluster-KO mice (Cat. NM-KO-18019) were purchased from Shanghai Model Organisms Center, Inc.

For analysing AMPK activation, wild-type and *AMPK*α-MKO mice of 4 weeks old were used; for determining rejuvenating effects of LCA: wild-type, and *AMPK*α-MKO mice, aged 18 months (treated with LCA for 1 month starting from 17 months old); for analysing the pharmacokinetics of LCA: wild-type mice aged 18 months (aged mice; treated with LCA or for 1 month starting from 17 months old); for determining the changes of LCA concentrations in serum and tissue after CR: wild-type mice aged 8 months old (subjected to CR for 4 months starting from 4 months old); for determination of serum metabolome in CR mice: wild-type mice aged 8 months old (subjected to CR for 4 months starting from 4 months old); and for isolating primary hepatocytes and myocytes: wild-type mice aged 1 month.

### CR, fasting, and cardiotoxin treatments of mice

Protocols for all rodent experiments were approved by the Institutional Animal Care and the Animal Committee of Xiamen University (XMULAC20180028 and XMULAC20220050). Unless stated otherwise, mice were housed with free access to water and standard diet (65% carbohydrate, 11% fat, 24% protein) under specific pathogen-free conditions. The light was on from 8:00 to 20:00, with the temperature kept at 21-24 °C and humidity at 40-70%. Only male mice were used in the study, and male littermate controls were used throughout the study.

Mice were individually caged for 1 week before each treatment. For fasting, the diet was withdrawn from the cage at 5 p.m., and mice were sacrificed at desired time points by cervical dislocation. For CR, each mouse was fed with 2.5 g of standard diet (approximately 70% of ad libitum food intake for a mouse at 4 months old and older) at 5 p.m. of each day. Cardiotoxin treatment was performed as described previously^125^. Briefly, mice were anesthetised with 3% isoflurane in air via a vaporiser (R540, RWD Life Science), followed by removing the hairs, some 50 µl of 20 µM cardiotoxin was injected intramuscularly into the tibialis anterior muscle. Muscles were analysed at day 7 post-cardiotoxin injection.

### Formulation and treatment of LCA

For cell-based experiments, LCA powder was dissolved in DMSO to a stock concentration of 500 mM, and was aliquoted and stored at −20 °C. The solution was placed at room temperature for 10 min (until no precipitate was visible) before adding to the culture medium. Note that any freeze-thaw cycle was not allowed to avoid the re-crystallisation of LCA (otherwise forms in sheet-like, insoluble crystals) in the stock solution.

For mouse experiments, LCA was coated with (2-hydroxypropyl)-β-cyclodextrin before treating animals. To coat LCA, the LCA powder was dissolved in 100 ml of methanol to a concentration of 0.01 g/ml, followed by mixing with 308 ml of (2-hydroxypropyl)-β-cyclodextrin solution (by dissolving (2-hydroxypropyl)-β-cyclodextrin in 30% (v/v, in water) methanol to 0.04 g/ml, followed by a 30 min of sonication). The control vehicle was similarly prepared, with no LCA added to the (2-hydroxypropyl)-β-cyclodextrin solution. After evaporating at 50 □, 90 r.p.m. by the rotary evaporator (Rotavator R-300, Vacuum Pump V-300, BUCHI), the coated powder was stored at 4 °C for no more than 2 weeks and was freshly dissolved in free drinking water to 1 g/l before feeding mice.

For nematode experiments, LCA at desired concentrations was freshly dissolved in DMSO, and was added to warm (cooled to approximately 60 °C after autoclaving) nematode growth medium^139^ (NGM, containing 0.3% (wt/vol) NaCl, 0.25% (wt/vol) bacteriological peptone, 1 mM CaCl_2_, 1 mM MgSO_4_, 25 mM KH_2_PO_4_-K_2_HPO_4_, pH 6.0, 0.02% (wt/vol) streptomycin, and 5 μg/ml cholesterol. The medium was used to make the NGM plate by adding 1.7% (wt/vol) agar. The plates were stored at 20 °C for no more than 3 days.

For fly experiments, LCA was coated and dissolved in water as in the mouse experiments, and was added to the BDSC Standard Cornmeal Medium^140^ (for regular culture), the 2% cornmeal-sugar-yeast (CSY) agar diet (for CR experiments; see ref. ^126^), or the 3% CSY agar diet (the control diet for CR experiments). The BDSC Standard Cornmeal Medium was prepared as described previously, with minor modifications^140^. Briefly, 60.5 g of dry yeast, 35 g of soy flour, 255.5 g of cornmeal, 20 g of agar, and 270 ml of corn syrup were mixed with 3,500 ml of water in a stockpot. The mixture was thoroughly stirred using a long-handled soup spoon, and then boiled, during which lumps formed were pressed out using the back of the spoon. After cooling to approximately 60 °C, 16.8 ml of propionic acid was added to the medium, followed by stirring with the spoon. The 2% or 3% CSY agar diets were prepared as in the BDSC Standard Cornmeal Medium, except that 175 g of cornmeal, 367.5 g of sucrose, 70 (for 2% CSY agar diet) or 105 g (for 3% CSY agar diet) of dry yeast, 24.5 g of agar, 16.8 ml of propionic acid and 3,500 ml water were used. The medium was then dispensed into the culture vials, 6 ml each. The media vials were covered with a single-layer gauze, followed by blowing with the breeze from a fan at room temperature overnight. Some 100 μl of LCA solution (coated with (2-hydroxypropyl)-β-cyclodextrin) at desired concentration was then layered (added dropwise) onto the medium surface of each vial, followed by blowing with the breeze for another 8 h at room temperature. The media vials were kept at 4 °C (for no more than 3 days) before experiment.

### Determination of mouse running capacity and grip strength

The maximal running capacity was determined as described previously^102, 141^, with minor modifications. Briefly, mice were trained on Rodent Treadmill NG (UGO Basile, cat. 47300) for 3 days with normal light-dark cycle, and tests were performed during the dark period. Before the experiment, mice were fasted for 2 h. The treadmill was set at 5° incline, and the speed of treadmill was set to increase in a ramp-mode (commenced at a speed of 5 m/min followed by an increase to a final speed of 25 m/min within 120 min). Mice were considered to be exhausted, and removed from the treadmill, following the accumulation of 5 or more shocks (0.1 mA) per minute for two consecutive minutes. The distances travelled and the endurance were recorded as the running capacity. Note that mice subjected to the test for running capacity were sacrificed and no longer used for other experiments.

Grip strength was determined on a grip strength meter (Ugo Basile, cat. 47200) following the protocol described previously^125^. Briefly, the mouse was held by its tail and lowered (“landed”) until forelimb or all four limbs grasped the T□bar connected to a digital force gauge. The mouse was further lowered to the extent that the body was horizontal to the apparatus, and was then slowly, steady drawn away from the T□bar until forelimb or all four limbs were removed from the bar, which gave rise to the peak force in grams. Each mouse was repeated 5 times with 5 min intervals between measurements. Note that the grip strength of forelimb and four limbs were measured at different days to prevent interference from muscle tiredness caused by earlier measurement.

### Serology, GTT and ITT

GTT and ITT were performed as described previously^104^. Before GTT and ITT, mice were individually caged for a week before experiment. For GTT, mice were fasted for 16 h (17:00 p.m. to 9:00 a.m.), then administered with glucose at 2 g/kg (intraperitoneally injected). For ITT, mice were fasted for 6 h (8:00 a.m. to 14:00 p.m.), then 0.5 U/kg insulin was intraperitoneally injected. Blood glucose was then measured at indicated time points through tail vein bleeding using the OneTouch UltraVue automatic Glucometer (LifeScan). Note that GTT and ITT were analysed using different batches of mice to avoid the interference from any stress caused by earlier blood collection.

For measuring insulin levels, approximately 100 μl of blood was collected (from submandibular vein plexus), and was placed at room temperature for 20 min, followed by centrifugation at 3,000*g* for 10 min at 4 °C. Some 25 μl of the resultant serum was used to determine the levels of insulin using the Mouse Ultrasensitive Insulin ELISA kit according to the manufacturer’s instructions. The five-parameter logistic fitted standard curve for calculating the concentrations of insulin was generated from the website of Arigo Biolaboratories (https://www.arigobio.cn/ELISA-calculator/). For measuring free fatty acids, glycerol, β-hydroxybutyrate and glucagon, 1.3 μl, 10 μl, 1 μl and 5 μl of freshly prepared serum from 8-h fasted mice was used, using the LabAssay NEFA kit, Free Glycerol Assay kit, Ketone Body Assay kit, and Mouse Glucagon ELISA Kit, respectively, all following the manufacturer’s instructions.

### Hyperinsulinemia-euglycemic clamp

The hyperinsulinemic-euglycemic clamp was performed as described previously^132, 142, 143^, with minor modifications. Briefly, mice were anesthetised with 3% isoflurane in air via a vaporiser (R540, RWD Life Science), followed by removing the hairs and disinfecting the skin with 70% (v/v, in water) ethanol at the incision site. A small incision located approximately 5 mm superior to the sternum and 5 mm to the right of the vertical midline was made, and the fat and connective tissues beneath the pectoral muscle and surrounding the right jugular vein were gently cleaned by blunt dissection. The cephalad end of the exposed vein was then tightly tied (forming an anterior ligature) through passing a 7-0 silk suture beneath the vein. After loosely tied the caudal end of the vein via another thread of suture (posterior ligature), the vein was inserted by a 19-G needle between the two threads a few millimeters below the anterior ligature. After removing the needle, a catheter (C10PUS-MFV1610, Instech) was inserted into the vein through the needle hole, with the bevel of its tip facing towards the opening, followed by pushing forwards towards the caudal end for approximately 1 cm distance (until the restraining bead reaching the superior vena cava). The catheter was flushed with 100 μl of heparin (200 U/ml, dissolved in saline), and then anchored by tightening the posterior ligature thread. The catheter was then tunnelled (pulled with eye dressing forceps) beneath the skin from the right jugular incision to the interscapular incision (approximately 5 mm in length) on the back. After exteriorising through the interscapular incision, the catheter was connected to a Mouse Vascular Access Button (VABM1BSM-25, Instech), sealed with a protective aluminium cap (VABM1C, Instech), and secured using the 6-0 silk suture. After closing the two incisions using the 6-0 silk suture, the catheterised mouse was allowed to recover for 4 days. Mice that lost < 4% of their pre-cannulation weight after recovery were used for clamp experiments.

One day prior to the experiments, a magnetic VAB tether kit (KVABM1T/25, Instech; with its 25-G luer stub (LS25/6, Instech) replaced by a PinPor-to-Tubing Connectors (PNP3MC/25, Instech) which is connected to a PU tube (VAHBPU-T25, Instech), followed by a 4-way X connector (SCX25, Instech), and three separate luer stubs (LS25, Instech): one for infusing unlabelled glucose, one for [U-^13^C]glucose, and the third for insulin - each connected with a PU tube) was anchored onto a counter-balanced lever arm (SMCLA, Instech), and was flushed with each perfusate by a PHD Ultra programmable syringe pump (HA3000P, Instech) in the following orders: unlabelled glucose (20% (m/v) in saline), 100 mU/ml insulin, and then 0.45 μg/μl [U-^13^C]glucose. The mouse VAB connector on the magnetic VAB tether kit was then connected to the Mouse Vascular Access Button on the back of catheterised mouse, after removing the protective aluminium cap. The mice were then fasted for 16 h (starting from 5:00 p.m. to 9:00 a.m. of the following day), and the experiment was performed with a two-phase protocol consisting of a 90-min equilibration period (t = −90 min to 0 min) and a 120-min experimental period (t = 0 min to 120 min). Some 0.45 μg/μl [U-^13^C]glucose was given at t = −90 min, and was infused at a rate of 30 μg/kg/min during the remaining time of the experiment. The clamping was begun at t = 0 min with a prime-continuous infusion of insulin (300 mU/kg/min for 1 min), followed by 25 mU/kg/min continuous infusion during the remaining time of the experiment. The unlabelled glucose (20% (m/v) in saline) was then infused at 50 mg/kg/min for 10 min (from t = 0 min to t = 10 min), and the rate was adjusted according to the blood glucose level (maintained at 6-7 mM; during which blood glucose was measured at every 10 min from t = 0 to 90 min, and every 5 min from t = 90 to 120 min), afterwards. At t = 0, 90, 100, 110 and 120 min (all at clamped state), some 60 μl of blood (collected from the tail vein) was taken from the tail vein, and the serum ratios of [U-^13^C]glucose to unlabelled glucose were determined using the ExionLC AD UPLC system (SCIEX) interfaced with a QTRAP 5500 MS (SCIEX), as described in the “Determination of serum metabolome” section, except that 10 μl of serum was used. The resting hepatic glucose output rate (HGP) was calculated by dividing the resting-state infusion rate of [U-^13^C]glucose with the ratio of [U-^13^C]glucose to unlabelled glucose, while the clamped HGP by dividing the value of differences between the average, clamp-state infusion rate of the unlabelled glucose and the [U-^13^C]glucose with the ratio of [U-^13^C]glucose to unlabelled glucose. The glucose disposal rate during clamping was the sum of the infusion rate of [U-^13^C]glucose, the infusion rate of unlabelled glucose, and the value of clamped HGP.

### Determination of body composition

Lean and fat body mass were measured by quantitative magnetic resonance (EchoMRI-100H Analyzer; Echo Medical Systems) as described previously^104^. Briefly, the system was calibrated with oil standard before the measurement. Mice were individually weighted and inserted into a restrainer tube, and were immobilised by gently inserting a plunger. The mouse was then positioned to a gesture that curled up like a donut, with its head against the end of the tube. Body composition of each mouse was measured with 3 repeated runs, and the average values were taken for further analysis.

### Determination of energy expenditure

Mouse energy expenditure (EE) was determined by a metabolic cage system (Promethion Line, CAB-16-1-EU; Sable Systems International) as described previously^104, 144^. Briefly, the system was maintained in a condition identical to that for housing mice. Each metabolic cage in the 16-cage system consisted of a cage with standard bedding, a food hopper and water bottle, connected to load cells for continuous monitoring. To minimise the stress of new environment, mice were acclimatised (by individually housing in the gas-calibrated chamber) for 1 week before data collection. Mice treated with LCA or vehicle control were randomly assigned/housed to prevent systematic errors in measurement. Body weights and fat proportion of mice were determined before and after the acclimation, and the food intake and water intake daily. Mice found not acclimatised to the metabolic cage (e.g., resist to eat and drink) were removed from the study. Data acquisition (5-min intervals each cage) and instrument control were performed using MetaScreen software (v.2.3.15.12, Sable Systems) and raw data processed using Macro Interpreter (v.2.32, Sable Systems). Ambulatory activity and position were monitored using XYZ beam arrays with a beam spacing of 0.25 cm (beam breaks), and the mouse pedestrial locomotion (walking distance) within the cage were calculated accordingly. Respiratory gases were measured using the GA-3 gas analyser (Sable Systems) equipped with a pull-mode, negative-pressure system. Air flow was measured and controlled by FR-8 (Sable Systems), with a set flow rate of 2,000 ml/min. Oxygen consumption (VO_2_) and carbon dioxide production (VCO_2_) were reported in ml per minute. Water vapour was measured continuously and its dilution effect on O_2_ and CO_2_ was compensated mathematically in the analysis stream. Energy expenditure (EE) was calculated using: kcal/h = 60 × (0.003941 × VO_2_ + 0.001106 × VCO_2_) (Weir Equation). Differences of average EE were analysed by analysis of covariance (ANCOVA) using body weight as the covariate. Respiratory quotient (RQ) was calculated as VCO_2_/VO_2_.

### Histology

For hematoxylin & eosin (H&E) staining, muscle tissues were quickly excised, followed by freezing in isopentane (pre-chilled in liquid nitrogen) for 2 min (until they appeared chalky white). The tissues were then quickly transferred to embedding molds containing O.C.T. Compound, and were frozen in liquid nitrogen for another 10 min. The embedded tissues were then sectioned into 6-μm slices at −20 °C using a CM1950 Cryostat (Leica), followed by fixing in 4% paraformaldehyde for 10 min, and washing with running water for 2 min at room temperature. The sections were stained in Mayer’s hematoxylin solution for 5 min, followed by washing in running water for 10 min, and were then stained in eosin Y-solution for another 1 min. The stained sections were dehydrated twice in 95% ethanol, 5 min each, twice in anhydrous ethanol, 1 min each, and two changes of xylene, 1 min each. The stained sections were mounted with Canada balsam and visualised on an AxioScan 7 scanner (Zeiss). Images were processed and analysed using Zen 3.4 software (Zeiss), and were formatted by Photoshop 2023 software (Adobe).

For immunohistochemistry staining of PAX7, tibialis anterior muscle tissues were excised, embedded and sectioned as in H&E staining. The sections were fixed with 4% paraformaldehyde 10 min, followed by washed with PBS for 5 min at room temperature. After incubating with PBST (PBS supplemented with 5% Triton X-100) for 10 min, the sections were blocked with BSA Solution (PBS containing 5% BSA) for 30 min at room temperature, followed by incubating with the PAX7 antibody (6 μg/ml, diluted in BSA Solution) for 12 h at 4 °C. The sections were then washed with PBS for 3 times, 5 min each at room temperature, followed by incubating with Alexa Fluor 488-cobjugated, goat anti-mouse IgG1 secondary antibody (1:200 diluted in BSA Solution) for 1 h at room temperature in a dark humidified chamber. The sections were washed with PBS for 3 times, 5 min each at room temperature, followed by incubating with 4% paraformaldehyde for 2 min, and then washed with PBS twice, 5 min each at room temperature. The section was then incubated with the laminin antibody (1:100 diluted in BSA Solution) for 3 h at room temperature in a dark humidified chamber, followed by washing with PBS buffer for 3 times, 5 min each at room temperature. The sections were then incubated with Alexa Fluor 594-conjugated, goat anti-rabbit IgG secondary antibody (1:200 diluted in BSA Solution) for 1 h at room temperature in a dark humidified chamber, followed by washing with PBS for 3 times, 5 min each at room temperature. Tissue sections were mounted with 90% glycerol and visualised on an LSM980 microscope (Zeiss). Images were processed and analysed on Zen 3.4 software (Zeiss), and formatted on Photoshop 2023 software (Adobe).

Muscle fibre types were determined as described previously^79, 145^, with minor modifications. Briefly, muscle tissues were excised, embedded and sectioned as in H&E staining. The sections were fixed in 4% paraformaldehyde for 10 min, and were then washed with PBS for 5 min at room temperature. After incubating with PBST (PBS supplemented with 5% (v/v) Triton X-100) for 10 min, the sections were blocked with BSA Solution (PBS containing 5% (m/v) BSA) for 30 min at room temperature. Muscle fibres were stained with the antibody against MHCIIb (6 μg/ml, diluted in BSA Solution) overnight at 4 °C, followed by washing with PBS for 3 times, 5 min each at room temperature. The sections were then incubated with Alexa Fluor 488-conjugated, goat anti-mouse IgM antibody (1:200 diluted in BSA Solution) for 1 h at room temperature in a dark humidified chamber, followed by washing with PBS for 3 times, 5 min each, incubated with 4% paraformaldehyde for 2 min, and then washed with PBS twice, 5 min each, all at room temperature. The sections were then incubated with antibody against MHCI (6 μg/ml, diluted in BSA Solution) for 3 h at room temperature in a dark humidified chamber, followed by washing with PBS buffer for 3 times, 5 min each at room temperature, and then incubated with Alexa Fluor 594-conjugated, goat anti-mouse IgG2b antibody (1:200 diluted in BSA Solution) for another 1 h at room temperature in a dark humidified chamber, followed by washing with PBS buffer for 3 times, 5 min each at room temperature. After fixing with 4% paraformaldehyde for 2 min and washing with PBS twice, 5 min each at room temperature, the sections were incubated with the antibody against MHCIIa (6 μg/ml, diluted in BSA Solution) for 3 h at room temperature in a dark humidified chamber, followed by washing with PBS buffer for 3 times, 5 min each at room temperature, and then incubated in Alexa Fluor 647-conjugated goat anti-mouse IgG1 antibody (1:200 diluted in BSA Solution) for another 1 h at room temperature in a dark humidified chamber, followed by washing with PBS buffer for 3 times, 5 min each at room temperature. Tissue sections were mounted with 90% glycerol and visualised on an LSM980 microscope (Zeiss). Images were processed and analysed on Zen 3.4 software (Zeiss), and formatted on Photoshop 2023 software (Adobe).

### *Caenorhabditis elegans* strains

Nematodes (hermaphrodites) were maintained on NGM plates spread with *E. coli* OP50 as standard food. All worms were cultured at 20 °C. Wildtype (N2 Bristol) and *aak-2* (ok524) strains were obtained from *Caenorhabditis* Genetics Center. All mutant strains were outcrossed 6 times to N2 before the experiments. Unless stated otherwise, worms were maintained on NGM plates spread with *Escherichia coli* OP50 as standard food. The administration of LCA was initiated at the L4 stage.

### Evaluation of nematode lifespan and healthspan

To determine the lifespan of nematodes, the worms were first synchronised: worms were washed off from agar plates with 15 ml of M9 buffer (22.1 mM KH_2_PO_4_, 46.9 mM Na_2_HPO_4_, 85.5 mM NaCl and 1 mM MgSO_4_) supplemented with 0.05% (v/v) Triton X-100 per plate, followed by centrifugation at 1,000*g* for 2 min. The worm sediment was suspended with 6 ml of M9 buffer containing 50% synchronising bleaching solution (by mixing 25 ml of NaClO solution (5% active chlorine), 8.3 ml of 25% (w/v) NaOH, and 66.7 ml of M9 buffer, for a total of 100 ml), followed by vigorous shaking for 2 min, and centrifugation for 2 min at 1,000*g*. The sediment was washed with 12 ml of M9 buffer twice, then suspended with 6 ml of M9 buffer, followed by rotating at 20 °C, 30 r.p.m. for 12 h. Synchronised worms were cultured to L4 stage before transfer to desired agar plates for determining lifespan. Worms were transferred to new plates every 2 d. Live and dead worms were counted during the transfer. Worms that displayed no movement upon gentle touching with a platinum picker were judged as dead. Kaplan-Meier curves were graphed by Prism 9 (GraphPad Software), and the statistical analysis data by SPSS 27.0 (IBM).

Pharyngeal pumping rates, assessed as the numbers of contraction-relaxation cycles of the terminal bulb on nematode pharynx within 1 min, were determined as described previously^146^, with minor modification. Briefly, the synchronised nematodes were cultured to L4 stage, LCA was administered thereafter. The 1-day-old nematodes were then picked and placed on a new NGM plate containing *E*. *coli*. After 10 min of incubation at room temperature, the contraction-relaxation cycles of the terminal bulb of each worm were recorded on a stereomicroscope (M165 FC, Leica) through a 63x objective for a consecutive 4 min using the Capture software (v.2021.1.13, Capture Visualisation), and the average contraction-relaxation cycles per min were calculated using the Aimersoft Video Editor software (v.3.6.2.0, Aimersoft).

The resistance of nematodes to the oxidative stress was determined as described previously^123^. Briefly, synchronised worms were cultured to L4 stage after which LCA was administered. After 2 days of LCA treatment, 20 worms were transferred to an NGM plate containing 15 mM FeSO_4_. Worms were then cultured at 20 □ on such a plate, during which the live and dead worms were counted at every 1 h.

### Drosophila melanogaster strains

All flies were cultured at 25 °C and 60% humidity with a 12-hour light and dark cycle. Adult flies were cultured in Bloomington *Drosophila* Stock Center (BDSC) Standard Cornmeal Medium (for regular culture), 2% (for CR), or 3% (the control, ad libitum fed group for CR) CSY agar diet. Larvae and the crossed fly strains were reared on Semi-Defined, Rich Medium, which is prepared as described previously^147^, with minor modifications. Briefly, 10 g of agar, 80 g of dry yeast, 20 g of yeast extract, 20 g of peptone, 30 g of sucrose, 60 g of glucose, 0.5 g of MgSO_4_·6H_2_O and 0.5g of CaCl_2_·6H_2_O were dissolved in 1,000 ml of di-distilled water, and then boiled, followed by cooling to 60 °C. Some 6 ml of propionic acid was then added to the medium, and the medium was dispensed into culture vials, 6 ml each. The media vials were covered with gauze and blown with the breeze as in BDSC and CSY diets, and were kept at 4 °C (for no more than 3 days) before experiment.

The wildtype fly strain (*w^1118^*; #3605) and the *GAL4*-expressing strain (*y^1^ w\**; *P*{*Act5C-GAL4-w*}*E1/CyO*; #25374) were obtained from the BDSC. The *GAL4*-induced, *AMPK*α RNAi-carrying strain (*w^1118^*; *P*{*GD736*}*v1827*; #1827) was obtained from the Vienna *Drosophila* Resource Center (VDRC). The *w^1118^*; *Sp/CyO* strain was obtained from the Core Facility of *Drosophila* Resource and Technology, Chinese Academy of Sciences. To obtain the flies with *AMPK*α knocked down on the *w^1118^* background, a *GAL4*-expressing strain on the *w^1118^* background (*w^1118^*; *P*{*Act5C-GAL4-w*}*E1/CyO*) was first generated by crossing the *y^1^ w\**; *P*{*Act5C-GAL4-w*}*E1/CyO* males with *w^1118^*; *Sp/CyO* females, followed by crossing the F_1_ males with straight wings (*w^1118^*; *P*{*Act5C-GAL4-w*}*E1/Sp*) with *w^1118^*; *Sp/CyO* females. The *GAL4*-expressing flies (*w^1118^* background) were then crossed with the *AMPK*α RNAi-carrying flies, and the F_1_ offspring with straight wings were the *AMPK*α-KD flies (*w^1118^*; *P*{*Act5C*-*GAL4*-*w*}*E1*/*P*{*GD736*}*v1827*; +/+). The F_1_ offspring of wildtype flies crossed with the *GAL4*-expressing flies (*w^1118^* background), i.e., the *w^1118^*; *P*{*Act5C*-*GAL4*-*w*}*E1*/+; +/+ flies, were used as the control files.

In this study, the following ages of flies were used: a) for analysing AMPK activation and the pharmacokinetics of LCA, third instar larvae or newly eclosed adults were used; b) for determining lifespan, adults at day 2 after eclosion were used (for LCA or CR treatment); c) for determining healthspan, mtDNA:nDNA, NAD^+^ levels and mitochondrial genes expression, adults at day 30 after eclosion (treated with LCA for 28 days starting from 2 days after eclosion) were used.

### Evaluation of lifespan and healthspan of flies

Fly lifespan was determined as described previously^148^, with minor modifications. Before the experiment, flies were synchronised: approximately 200 pairs of flies, housed 10 pairs per tube, were cultured in Semi-Defined, Rich Medium and allowed to lay eggs for a day. After discarding the parent flies, the embryos were cultured for another 10 days, and the flies eclosed at day 12 were anaesthetised and collected with CO_2_ (those emerged before day 12 were discarded), followed by transferring to the BDSC Standard Cornmeal Medium and cultured for another 2 days. The male and female adults were then sorted by briefly anaesthetised with CO_2_ on the anaesthetic pad using a homemade feather brush (by attaching the apical region of a vane from the secondary coverts of an adult goose, to a plastic balloon stick), and some 200 adults of each group/gender were randomly assigned to the BDSC Standard Cornmeal Medium or the CSY medium, containing LCA or not, 20 flies per tube. The files were transferred to new medium tubes every two days without anaesthesia until the last survivor was dead. During each tube transfer, the sum of dead flies in the old tubes and the dead flies carried to the new tubes were recorded as the numbers of deaths, and the escaped or accidentally died flies (i.e., died within 3 days of same-sex culturing, or squeezed by the tube plugs) were censored from the experiments. Kaplan-Meier curves were graphed by Prism 9 (GraphPad Software), and the statistical analysis data by SPSS 27.0 (IBM).

The resistance of flies to the oxidative stress was determined as described previously^149^. Briefly, synchronised adults were treated with LCA for 30 days, followed by transferring to vials (20 flies each), each containing a filter paper soaked with 20 mM paraquat or 5% (m/v) H_2_O_2_ dissolved/diluted in 5% (w/v, in water) glucose solution. To determine the resistance of flies to cold and heat stress, synchronised adults were treated with LCA for 30 days, followed by transferring to cold (4 °C) or heat (37 °C) stress conditions. To determine the resistance of flies to starvation (food deprivation), flies treated with LCA for 30 days were transferred to vials with culture medium replaced by the same volume of 1.5% agarose to remove food supply. Dead files were recorded every 2 h until the last survivor was dead.

### Quantification of mRNA levels of mitochondrial genes in mice, nematodes and flies

Mice treated with LCA were sacrificed by cervical dislocation, immediately followed by dissecting gastrocnemius muscle. The muscle tissue was roughly sliced to cubes having edge lengths of approximately 2 mm, and then soaked in RNAprotect Tissue Reagent (1 ml per 100 mg of tissue) for 24 h at room temperature. The tissue was then incubated in 1 ml of TRIzol, followed by three rounds of freeze/thaw cycles, and was then homogenised. The homogenate was centrifuged at 12,000g for 15 min at 4 °C, and 900 µl of clear supernatant (not the lipid layer on the top) was transferred to an RNase-free tube. The supernatant was then added with 200 µl of chloroform, followed by vigorous vortex for 15 s. After centrifugation at 12,000*g* for 15 min at 4 °C, 450 µl of the upper aqueous layer was transferred to an RNase free tube. The RNA was then precipitated by adding 450 µl of isopropanol, followed with centrifugation at 12,000*g* for 30 min at 4 °C. The pellet was washed twice with 75% ethanol, and once with 100% ethanol, and was dissolved with 20 µl of DEPC-treated water. The concentration of RNA was determined by a NanoDrop 2000 spectrophotometer (Thermo). Some 1 µg of RNA was diluted with DEPC-treated water to a final volume of 10 µl, heated at 65 °C for 5 min, and chilled on ice immediately. The Random Primer Mix, Enzyme Mix and 5× RT buffer (all from the ReverTra Ace qPCR RT Master Mix) were then added to the RNA solution, followed by incubation at 37 °C for 15 min, and then at 98 °C for 5 min on a thermocycler. The reverse-transcribed cDNA was quantified with Maxima SYBR Green/ROX qPCR Master Mix on a LightCycler 480 II System (Roche) with following programmes: pre-denaturing at 95 °C for 10 min; denaturing at 95 °C for 10 s, then annealing and extending at 65 °C for 30 s in each cycle (determined according to the amplification curves, melting curves, and bands on agarose gel of serial pilot reactions (in which a serial annealing temperature was set according to the estimated annealing temperature of each primer pair) of each primer pair, and same hereafter), for a total of 45 cycles. Primer pairs for mouse *Nd1*, *Nd2*, *Nd3*, *Nd4*, *Nd4l*, *Nd5*, *Nd6*, *Ndufab1*, *Cytb*, *Uqcrc1*, *Uqcrc2*, *Atp5f1b*, *Cox6a1*, *Atp6*, *Atp8*, *Cox1* and *Cox3* were generated as described previously^150^, and others using the Primer-BLAST website (https://www.ncbi.nlm.nih.gov/tools/primer-blast/index.cgi). Primer sequence are as follows: mouse *Gapdh*, 5’-GACTTCAACAGCAACTCCCAC-3’ and 5’-TCCACCACCCTGTTGCTGTA-3’; mouse *Nd1*, 5’-TGCACCTACCCTATCACTCA-3’ and 5’-C GGCTCATCCTGATCATAGAATGG-3’; mouse *Nd2*, 5’-ATACTAGCAATTACTTCTATTTTCATAGGG-3’ and 5’-GAGGGATGGGTTGTAAGGAAG-3’; mouse *Nd3*, 5’-AAGCAAATCCATATGAATGCGG-3’ and 5’-GCTCATGGTAGTGGAAGTAGAAG-3’; mouse *Nd4*, 5’-CCTCAGACCCCCTATCCACA-3’ and 5’-GTTTGGTTCCCTCATCGGGT-3’; mouse *Nd4l*, 5’-CCAACTCCATAAGCTCCATACC-3’ and 5’-GATTTTGGACGTAATCTGTTCCG-3’; mouse *Nd5*, 5’-ACGAAAATGACCCAGACCTC-3’ and 5’-GAGATGACAAATCCTGCAAAGATG-3’; mouse *Nd6*, 5’-TGTTGGAGTTATGTTGGAAGGAG-3’ and 5’-CAAAGATCACCCAGCTACTACC-3’; mouse *Tfam*, 5’-GGTCGCATCCCCTCGTCTAT-3’ and 5’-TTGGGTAGCTGTTCTGTGGAA-3’; mouse *Cs*, 5’-CTCTACTCACTGCAGCAACCC-3’ and 5’-TTCATGCCTCTCATGCCACC-3’; mouse *Ndufs8*, 5’-TGGCGGCAACGTACAAGTAT-3’ and 5’-GTAGTTGATGGTGGCAGGCT-3’; mouse *Ndufab1*, 5’-GGACCGAGTTCTGTATGTCTTG-3’ and 5’-AAACCCAAATTCGTCTTCCATG-3’; mouse *Ndufb10*, 5’-TGCCAGATTCTTGGGACAAGG-3’ and 5’-GTCGTAGGCCTTCGTCAAGT-3’; mouse *Ndufv3*, 5’-GTGTGCTCAAAGAGCCCGAG-3’ and 5’-TCAGTGCCGAGGTGACTCT-3’; mouse *Ndufa8*, 5’-GCGGAGCCTTTCACAGAGTA-3’ and 5’-TCAATCACAGGGTTGGGCTC-3’; mouse *Ndufs3*, 5’-CTGACTTGACGGCAGTGGAT-3’ and 5’-CATACCAATTGGCCGCGATG-3’; mouse *Ndufa9*, 5’-TCTGTCAGTGGAGTTGTGGC-3’ and 5’-CCCATCAGACGAAGGTGCAT-3’; mouse *Ndufa10*, 5’-CAGCGCGTGGGACGAAT-3’ and 5’-ACTCTATGTCGAGGGGCCTT-3’; mouse *Sdha*, 5’-AGGGTTTAATACTGCATGCCTTA-3’ and 5’-TCATGTAATGGATGGCATCCT-3’; mouse *Sdhb*, 5’-AGTGCGGACCTATGGTGTTG-3’ and 5’-AGACTTTGCTGAGGTCCGTG-3’; mouse *Sdhc*, 5’ -TGAGACATGTCAGCCGTCAC-3’ and 5’-GGGAGACAGAGGACGGTTTG-3’; mouse *Sdhd*, 5’-TGGTACCCAGCACATTCACC-3’ and 5’-GGGTGTCCCCATGAACGTAG-3’; mouse *Cytb*, 5’ -CCCACCCCATATTAAACCCG-3’ and 5’-GAGGTATGAAGGAAAGGTATTAGGG-3’; mouse *Uqcrc1*, 5’-ATCAAGGCACTGTCCAAGG-3’ and 5’-TCATTTTCCTGCATCTCCCG-3’; mouse *Uqcrc2*, 5’-TTCCAGTGCAGATGTCCAAG-3’ and 5’-

CTGTTGAAGGACGGTAGAAGG-3’; mouse *Atp5f1b*, 5’-CCGTGAGGGCAATGATTTATAC-3’ and 5’-GTCAAACCAGTCAGAGCTACC-3’ mouse *Cox6a1*, 5’-GTTCGTTGCCTACCCTCAC-3’ and 5’-TCTCTTTACTCATCTTCATAGCCG-3’; mouse *Atp6*, 5’-TCCCAATCGTTGTAGCCATC-3’ and 5’-TGTTGGAAAGAATGGAGTCGG-3’; mouse *Atp8*, 5’-GCCACAACTAGATACATCAACATG-3’ and 5’-TGGTTGTTAGTGATTTTGGTGAAG-3’; mouse *Atp5f1a*, 5’-CATTGGTGATGGTATTGCGC-3’ and 5’-TCCCAAACACGACAACTCC-3’; mouse *Cox1*, 5’-CCCAGATATAGCATTCCCACG-3’ and 5’-ACTGTTCATCCTGTTCCTGC-3’; mouse *Cox2*, 5’-TCTACAAGACGCCACATCCC-3’ and 5’-ACGGGGTTGTTGATTTCGTCT-3’; mouse *Cox3*, 5’-CGTGAAGGAACCTACCAAGG-3’ and 5’-CGCTCAGAAGAATCCTGCAA-3’; mouse *Cox5b*, 5’-AGCTTCAGGCACCAAGGAAG-3’ and 5’-TGGGGCACCAGCTTGTAATG-3’. The mRNA level was then calculated using the comparative ΔΔct method using the LightCycler software (v.96 1.1, Roche; same hereafter for all qPCR experiments).

Nematodes at L4 stage treated with LCA for 1 day were used for analysis of mitochondrial gene expression. Some 1,000 worms were collected with 15 ml of M9 buffer containing 0.05% Triton X-100 (v/v), followed by centrifugation for 2 min at 1,000*g*. The sediment was then washed with 1 ml of M9 buffer twice, and then lysed with 1 ml of TRIzol. Worms were then frozen in liquid nitrogen, thawed at room temperature, and then repeated freeze-thaw for another 2 times. The worm lysates were then placed at room temperature for 5 min, then mixed with 0.2 ml of chloroform, followed by vigorous shaking for 15 s. After centrifugation at 12,000*g* for 15 min at 4 °C, 450 µl of the upper aqueous layer was transferred to an RNase free tube. The RNA was then precipitated by adding 450 µl of isopropanol, followed with centrifugation at 12,000*g* for 30 min at 4 °C. The pellet was washed twice with 75% ethanol, and once with 100% ethanol, and was dissolved with 20 µl of DEPC-treated water. The concentration of RNA was determined by a NanoDrop 2000 spectrophotometer (Thermo). Some 1 µg of RNA was diluted with DEPC-treated water to a final volume of 10 µl, heated at 65 °C for 5 min, and chilled on ice immediately. The Random Primer Mix, Enzyme Mix and 5× RT buffer (all from the ReverTra Ace qPCR RT Master Mix) were then added to the RNA solution, followed by incubation at 37 °C for 15 min, and then at 98 °C for 5 min on a thermocycler. The reverse-transcribed cDNA was quantified with Maxima SYBR Green/ROX qPCR Master Mix on a LightCycler 480 II System (Roche) with following programmes: pre-denaturing at 95 °C for 10 min; denaturing at 95 °C for 10 s, then annealing and extending at 65 °C for 30 s in each cycle, for a total of 45 cycles. Primer pairs used for qPCR are as previously described^151, 152^, except that *C*. *elegans ctb-1* were designed using the Primer-BLAST website. Primer sequence are as follows: *C*. *elegans ama-1*, 5’-GACATTTGGCACTGCTTTGT-3’ and 5’-ACGATTGATTCCATGTCTCG-3’; *C*. *elegans nuo-6*, 5’-CTGCCAGGACATGAATACAATCTGAG-3’ and 5’-GCTATGAGGATCGTATTCACGACG-3’; *C*. *elegans nuaf-1*, 5’-GAGACA TAACGAGGCTCGTGTTG-3’ and 5’-GAAGCCTTCTTTCCAATCACTATCG-3’; *C*. *elegans sdha-1*, 5’-TTACCAGCGTGCTTTCGGAG-3’ and 5’-AGGGTGTGGAGAAGAGAATGACC-3’; *C*. *elegans sdhb-1*, 5’-GCTGAACGTGATCGTCTTGATG-3’ and 5’-GTAGGATGGGCATGACGTGG-3’; *C*. *elegans cyc-2.1*, 5’-CGGA GTTATCGGACGTACATCAG-3’ and 5’-GTCTCGCGGGTCCAGACG-3’; *C*. *elegans isp-1*, 5’-GCAGAAAGATGAATGGTCCGTTG-3’ and 5’-ATCCGTGACAAGGGCAGTAATAAC-3’; *C*. *elegans cco-1*, 5’-GCTGGAGATGATCGTTACGAG-3’ and 5’-GCATCCAATGATTCTGAAGTCG-3’; *C*. *elegans cco-2*, 5’-GTGATACCGTCTACGCCTACATTG-3’ and 5’-GCTCTGGCACGAAGAATTCTG-3’; *C*. *elegans atp-3*, 5’-GTCCTCGACCCAACTCTCAAG-3’ and 5’-GTCCAAGGAAG TTTCCAGTCTC-3’; *C*. *elegans nduo-1*, 5’-AGCGTCATTTATTGGGAAGAAGAC-3’ and 5’-AAGCTTGTGCTAATCCCATAAATGT-3’; *C*. *elegans nduo-2*, 5’-TCTT TGTAGAGGAGGTCTATTACA-3’ and 5’-ATGTTAAAAACCACATTAGCCCA-3’; *C. elegans nduo-4*, 5’-GCACACGGTTATACATCTACACTTATG-3’ and 5’-GATGTATGATAAAATTCACCAATAAGG-3’; *C*. *elegans nduo-5*, 5’-AGATGAGATTTATTGGGTATTTCTAG-3’ and 5’-CACCTAGACGATTAGTTAATGCTG-3’; *C*. *elegans ctc-1*, 5’-GCAGCAGGGTTAAGATCTATCTTAG-3’ and 5’-CTGTTACAAATACAGTTCAAACAAAT-3’; *C*. *elegans ctc-2*, 5’-GTAGTTTATTGTTGGGAGTTTTAGTG-3’ and 5’-CACAATAATTCACCAAACTGATACTC-3’; *C*. *elegans atp-6*, 5’-TGCTGCTGTAGCGTGATTAAG-3’ and 5’-ACTGTTAAAGCAAGTGGACGAG-3’; *C*. *elegans ctb-1*, 5’-TGGTGTTACAGGGGCAACAT-3’ and 5’-TGGCCTCATTATAGGGTCAGC-3’.

*Drosophila* adults treated with LCA for 30 days were used to determine the expression of mitochondrial genes. For each sample, some 20 adults were used. The adults were anesthetised, transferred to a 1.5-ml Eppendorf tube, followed by quickly freezing in liquid nitrogen, and then homogenised using a pellet pestle (cat. Z359963-1EA, Sigma). The homogenate was then lysed in 1 ml of TRIzol for 5 min at room temperature, followed by centrifuged at 12,000*g* for 15 min at 4 °C. Some 900 μl of supernatant (without the lipid layer) was transferred to an RNase-free tube, followed by mixing with 200 μl of chloroform. After vigorous vortexing for 15 s, the mixture was centrifuged at 12,000*g* for 15 min at 4°C, and some 450 μl of the upper aqueous layer was transferred to an RNase-free tube. The RNA was then precipitated by adding 450 μl of isopropanol, followed by centrifugation at 12,000*g* for 30 min at 4 °C. The pellet was washed twice with 75% (v/v, in water) ethanol, and was dissolved with 20 μl of DEPC-treated water. The concentration of RNA was determined by a NanoDrop 2000 spectrophotometer (Thermo). Some 1 µg of RNA was diluted with DEPC-treated water to a final volume of 10 µl, heated at 65 °C for 5 min, and chilled on ice immediately. The Random Primer Mix, Enzyme Mix and 5× RT buffer (all from the ReverTra Ace qPCR RT Master Mix) were then added to the RNA solution, followed by incubation at 37 °C for 15 min, and then at 98 °C for 5 min on a thermocycler. The reverse-transcribed cDNA was quantified with Maxima SYBR Green/ROX qPCR Master Mix on a LightCycler 480 II System (Roche) with following programmes: pre-denaturing at 95 °C for 5 min; denaturing at 95 °C for 10 s, then annealing at 60 °C for 20 s, and then extending at 72 °C for 20 s in each cycle, for a total of 40 cycles. Primer pairs used for qPCR are as previously described^153^, and are listed as follows: *D*. *melanogaster CG9172*, 5’-CGTGGCTGCGATAGGATAAT-3’ and 5’-ACCACATCTGGAGCGTCTTC-3’; *D*. *melanogaster CG9762*, 5’-AGTCACCGCATTGGTTCTCT-3’ and 5’-GAGATGGGGTGCTTCTCGTA-3’; *D*. *melanogaster CG17856*, 5’-ACCTTTCCATGACCAAGACG-3’ and 5’-CTCCATTCCTCACGCTCTTC-3’; *D*. *melanogaster CG18809*, 5’-AAGTGAAGACGCCCAATGAGA-3’ and 5’-GCCAGGTACAACGACCAGAAG-3’; *D*. *melanogaster CG5389*, 5’-ATGGCTACAGCATGTGCAAG-3’ and 5’-GACAGGGAGGCATGAAGGTA-3’; *D. melanogaster Act5C*, 5’-GCAGCAACTTCTTCGTCACA-3’ and 5’-CATCAGCCAGCAGTCGTCTA-3’.

### Analysis of mitochondrial DNA copy numbers in mice, nematodes and flies

Mouse mitochondrial DNA copy numbers were determined as described previously^104^. Briefly, mouse tissue DNA was extracted with the Biospin tissue genomic DNA extraction kit (BioFlux) following the manufacturer’s instruction, with minor modifications. Briefly, mice treated with LCA were sacrificed by cervical dislocation, quickly followed by dissecting gastrocnemius muscle. The muscle tissue was then grinded on a ceramic mortar in liquid nitrogen. Some 50 mg of grinded tissue was then transferred to a 1.5-ml Eppendorf tube, followed by addition of 600 µl of FL buffer and 10 µl of PK solution containing 2 µl of 100 mg/ml RNase A. The mixture was then incubated at 56 □ for 15 min, followed by centrifuge at 12,000*g* for 3 min. Some 500 µl of supernatant was transferred to a 2-ml Eppendorf tube, followed by mixing with 700 µl of binding buffer and 300 µl of absolute ethanol. The mixture was then loaded onto a Spin column, and was centrifuged at 10,000*g* for 1 min. The flowthrough was discarded, and 500 µl of the PW buffer was added to the Spin column, followed by centrifuge at 10,000*g* for 30 sec. Some 600 µl of washing buffer was then added to the spin column, followed by centrifuge at 10,000*g* for 30 sec, and then repeated once. The Spin column was then centrifuged for 1 min at 10,000*g* to completely remove the washing buffer, and the DNA on the column was eluted by 100 µl of Elution buffer (added to Spin column, followed by incubation at room temperature for 5 min, and then centrifuged at 12,000*g* for 1 min). Total DNA was quantified with Maxima SYBR Green/ROX qPCR Master Mix on a LightCycler 480 II System (Roche) with following programmes: 70 ng of DNA was pre-denatured at 95 °C for 10 min, and then subjected to PCR for a total of 45 cycles: denaturing at 95 °C for 10 s, annealing and extending at 65 °C for 30 s in each cycle. Primer pairs used for qPCR are as previously described^154^ (mouse *Hk2*, 5’-GCCAGCCTCTCCTGATTTTAGTGT-3’ and 5’-GGGAACACAAAAGACCTCTTCTGG-3’; mouse *Nd1*, 5’-CTAGCAGAAACAAACCGGGC-3’ and 5’-CCGGCTGCGTATTCTACGTT-3’).

Nematode mitochondrial DNA copy numbers were determined from worm lysates as described previously^104^. Briefly, 30 synchronised early L4 worms were collected, and were lysed with 10 μl of worm lysis buffer (50 mM HEPES, pH7.4, 1 mM EGTA, 1 mM MgCl_2_, 100 mM KCl, 10% (v/v) glycerol, 0.05% (v/v) NP-40, 0.5 mM DTT, and protease inhibitor cocktail). The worm lysate was frozen at −80 □ overnight, followed by incubating at 65 □ for 1 h and 95 □ for 15 min. Nematode DNA was then quantified with Maxima SYBR Green/ROX qPCR Master Mix on a LightCycler 480 II System (Roche) with following programmes: pre-denaturing at 95 °C for 10 min and then for a total of 45 cycles of denaturing at 95 °C for 10 s, and annealing and extending at 65 °C for 30 s in each cycle. Primer pairs used for qPCR are designed as described previously^124^ (*C*. *elegans nd-1*, 5’-AGCGTCATTTATTGGGAAGAAGAC-3’ and 5’-AAGCTTGTGCTAATCCCATAAATGT-3’; *C*. *elegans act-3*, 5’-TGCGACATTGATATCCGTAAGG-3’ and 5’-GGTGGTTCCTCCGGAAAGAA-3’).

*Drosophila* DNA copy numbers were determined as described previously^155^, with minor modifications. Briefly, some 20 anaesthetised adults were homogenised in 100 μl of Fly Lysis Buffer (75 mM NaCl, 25 mM EDTA, 25 mM HEPES, pH7.5) containing proteinase K (100 μg/ml). The homogenate was then frozen at −80 □ for 12 h, followed by incubating at 65 □ for 1 h and 95 □ for another 15 min. The fly DNA was then quantified using the Maxima SYBR Green/ROX qPCR Master Mix on a LightCycler 480 II System (Roche) with following programmes: pre-denaturing at 95 °C for 5 min and then for a total of 40 cycles of denaturing at 95 °C for 10 s, and annealing 60 □ for 20 s and extending at 72 °C for 20 s in each cycle. Primer pairs used for qPCR are as previously described^155^ (*D. melanogaster 16S rRNA*, 5’-TCGTCCAACCATTCATTCCA-3’ and 5’-TGGCCGCAGTATTTTGACTG-3’; *D. melanogaster RpL32*, 5’-AGGCCCAAGATCGTGAAGAA-3’ and 5’-TGTGCACCAGGAACTTCTTGAA-3’).

### Determining muscle atrophy markers in mice

Muscle atrophy markers were determined as described previously^156^, with minor modifications. Briefly, mice treated with LCA were sacrificed by cervical dislocation, followed by quickly dissecting gastrocnemius muscle. The muscle tissue was sliced to cubes having edge lengths of approximately 2 mm, and then soaked in RNAprotect Tissue Reagent (1 ml per 100 mg of tissue) for 24 h at room temperature. The tissue was then incubated in 1 ml of TRIzol, followed by three rounds of freeze/thaw cycles, and was then homogenised. The homogenate was centrifuged at 12,000*g* for 15 min at 4 °C, and 900 µl of the clear supernatant (without the lipid layer) was transferred to an RNase-free tube. The supernatant was then added with 200 µl of chloroform, followed by vigorous vortex for 15 s. The RNA was then purified, and was transcribed to cDNA as described in the “Quantification of mRNA levels of mitochondrial genes in mice, nematodes and flies” section. The reverse-transcribed cDNA was quantified with Maxima SYBR Green/ROX qPCR Master Mix on a LightCycler 480 II System (Roche) with following programmes: pre-denaturing at 95 °C for 10 min; denaturing at 95 °C for 10 s, then annealing and extending at 65 °C for 30 s in each cycle for a total of 45 cycles. Primer sequences are as previously described^156^ (mouse *Fbxo32*, 5’-TAGTAAGGCTGTTGGAGCTGATAG-3’ and 5’-CTGCACCAGTGTGCATAAGG-3’; mouse *Trim63*, 5’-CATCTTCCAGGCTGCGAATC-3’; and 5’-ACTGGAGCACTCCTGCTTGT-3’; mouse *Gapdh*, 5’-TTCACCACCATGGAGAAGGC-3’ and 5’-CCCTTTTGGCTCCACCCT-3’).

### Primary hepatocytes and myocytes

Mouse primary hepatocytes were isolated with a modified two-step perfusion method using Liver Perfusion Media and Liver Digest Buffer as described previously^102^. Before isolation of hepatocytes, mice were first anaesthetised, followed by inserting a 0.72 mm × 19 mm I.V. catheter into postcava. After cutting off the portal vein, mice were perfused with 50 ml of Liver Perfusion Media at a rate of 5 ml/min, followed with 50 ml of Liver Digest Buffer at a rate of 2.5 ml/min. The digested liver was then briefly rinsed by PBS, and then dismembered by gently tearing apart the Glisson’s capsule with two sterilised, needle-pointed tweezers on a 6-cm dish containing 3 ml of PBS. The dispersed cells were mixed with 10 ml of ice-cold William’s medium E plus 10% FBS, and were filtered by passing through a 100-μm Cell Strainer (cat. 352360; Falcon). Cells were then centrifuged at 50*g* at 4 °C for 2 min, followed by washing twice with 10 ml of ice-cold William’s medium E plus 10% FBS. Cells were then immediately plated (at 60-70% confluence) in collagen-coated 6-well plates in William’s medium E plus 10% FBS, 100 IU penicillin and 100 mg/ml streptomycin, and were maintained at 37 °C in a humidified incubator containing 5% CO_2_. After 4 h of attachment, the medium was replaced with fresh William’s medium E with 1% (w/v) BSA for another 12 h before further use.

Mouse primary myocytes were isolated as described previously^157^. Briefly, mice were sacrificed by cervical dislocation, and hindlimb muscles from both legs were excised. Tissues were minced and digested in a collagenase B/dispase/CaCl_2_ solution for 1.5 h at 37 °C on a shaking bath. DMEM supplemented with 10% fetal bovine serum (FBS) was then added to the digested tissues, the mixtures were gently triturated, followed by loading onto a 70-μm Strainer filter (cat. 352350; Falcon). Cell suspensions were then centrifuged at 1,000*g* for 5 min, and the pellets were resuspended in growth medium (Ham’s F-10 medium supplemented with 20% FBS and 2.5 ng/ml bFGF). Cells were then plated on collagen-coated dishes (cat. 354456, Corning) at 60-70% confluence.

### Determination of serum metabolome

For measuring serum metabolome, blood samples from CR and libitum-fed mice were collected at 15:00 pm, followed by incubation at room temperature for 10 min, and then centrifuge at 3,000*g* at 4°C for another 10 min. The supernatants were serum samples, which were prepared on the same day of blood collection. For heat-inactivation, serum was incubated in at 56 °C for 30 min in a water bath. For dialysis, serum was loaded into a D-Tube Dialyzer Maxi (with molecular weight cutoffs from 3.5 to 14 kDa; cat. 71508), and dialysed in a beaker containing 2 l of PBS at 4 °C for 24 h on a magnetic stirrer. The PBS was refreshed at every 4 h.

Polar metabolites were determined via HPLC-MS, GC-MS and CE-MS, as described previously^101, 104, 158^, with minor modifications^159^. For HPLC-MS and CE-MS, some 100 μl of serum was instantly mixed with 1 ml of pre-cooled methanol containing IS1 (50 µM L-methionine sulfone, 50 µM D-campher-10-sulfonic acid, dissolved in water; 1:500 (v/v) added to the methanol and used to standardise the metabolite intensity and to adjust the migration time), then mixed with 1 ml of chloroform and 400 μl of water (containing 4 μg/ml [U-^13^C]-glutamine), followed with 20 s of vortexing. After centrifugation at 15,000*g* for another 15 min at 4 °C, the supernatant (aqueous phase) was then divided into 3 portions: (a) 200 μl, for HPLC-MS analysis; (b) 200 μl, for CE-MS analysis on anion mode; and (c) 200 μl, for CE-MS analysis on cation mode. Portion (a) was then lyophilised in a vacuum concentrator (CentriVap Benchtop Centrifugal Vacuum Concentrator (cat. #7310037; Labconco), equipped with a CentriVap −84 °C Cold Trap (cat. #7460037; Labconco) and an EDWARDS nXDS15i pump) at 4 °C for 12 h, and then dissolved in 50 μl of 50% (v/v, in water) acetonitrile, followed by centrifugation at 15,000*g* for another 30 min at 4 °C. Some 20 µl of supernatant was loaded into an injection vial (cat. 5182-0714, Agilent Technologies; with an insert (cat. HM-1270, Zhejiang Hamag Technology)) equipped with a snap cap (cat. HM-2076, Zhejiang Hamag Technology), and 2 μl of supernatant was injected into an HILIC column (ZICpHILIC, 5 μm, 2.1 mm × 100 mm, PN: 1.50462.0001, Millipore) on an ExionLC AD UPLC system (SCIEX) which is interfaced with a QTRAP 5500 MS (SCIEX). The mobile phase consisted of 15 mmol/l ammonium acetate containing 3 ml/l ammonium hydroxide (>28%, v/v) in the LC-MS grade water (mobile phase A), and LC-MS grade 90% (v/v) acetonitrile in LC-MS grade water (mobile phase B), and was run at a flow rate of 0.2 ml/min. The HPLC gradient elution programme was: 95% B held for 2 min, then to 45% B in 13 min, held for 3 min, and then back to 95% B for 4 min. Each sample was analysed on both the positive and the negative modes on the HPLC-MS. The MS was run on a Turbo V ion source with spray voltages of −4,500 V (negative mode) and 5,500 V (positive mode), source temperature at 550 °C, Gas No.1 at 50 psi, Gas No.2 at 55 psi, and curtain gas at 40 psi. Metabolites were measured using the multiple reactions monitoring mode (MRM), and declustering potentials and collision energies were optimised through using analytical standards. Data were collected using Analyst software (v.1.7.1, SCIEX), and the relative amounts of metabolites were analysed using MultiQuant software (v.3.0.3, SCIEX). Portions (b) and (c) of supernatant were then filtrated through a 5-kDa cut-off filter (cat. OD003C34, PALL) by centrifuging at 12,000*g* for 3 h at 4 °C. The filtered aqueous phase was then lyophilised at 4 °C, and then re-dissolved in 100 μl of water containing IS2 (50 µM 3-aminopyrrolidine dihydrochloride, 50 µM N,N-diethyl-2-phenylacetamide, 50 µM trimesic acid, 50 µM 2-naphtol-3,6-disulfonic acid disodium salt, dissolved in methanol; used to adjust the migration time; 1:200 for portion (b), or 1:400 for portion (c). Some 20 μl of re-dissolved potion (b) and portion (c) solutions were then loaded into injection vials an injection vial (cat. 9301-0978, Agilent Technologies; equipped with a snap cap (cat. 5042-6491, Agilent Technologies)). Before CE-MS analysis, the fused-silica capillary (cat. TSP050375, i.d. 50 µm × 80 cm; Polymicro Technologies) was installed in a CE/MS cassette (cat. G1603A; Agilent Technologies) on the 7100 CE system (Agilent Technologies). For anion mode, the capillary was pre-conditioned with 1 M NaOH for 0.5 h, flushed by di-distilled water for 2 h, and then washed with Anion Conditioning Buffer (25 mM ammonium acetate, 75 mM diammonium hydrogen phosphate, pH 8.5) for 1 h, followed by balanced with Anion Running Buffer (50 mM ammonium acetate, pH 8.5; freshly prepared) for another 2 h. The capillary was then washed again by the Anion Conditioning Buffer for 5 min, followed by injection of the samples at a pressure of 50 mbar for 25 s, and then separation with a constant voltage at −30 kV for another 40 min in the Anion Running Buffer. Sheath Liquid (0.1 μM hexakis(1H, 1H, 3H-tetrafluoropropoxy)phosphazine, 10 μM ammonium trifluoroacetate, dissolved in methanol/water (50% v/v); freshly prepared) was flowed at 1 ml/min through a 1:100 flow splitter (1260 Infinity II; Agilent Technologies; actual flow rate to the MS: 10 μl/min) throughout each run. The parameters of 6545 MS (Agilent Technologies) were set as: a) ion source: Dual AJS ESI; b) polarity: negative; c) nozzle voltage: 2,000 V; d) fragmentor voltage: 110 V; e) skimmer voltage: 50 V; f) OCT RFV: 500 V; g) drying gas (N_2_) flow rate: 7 l/min; h) drying gas (N_2_) temperature: 300 °C; i) nebulizer gas pressure: 8 psig; j) sheath gas temperature: 125 °C; k) sheath gas (N_2_) flow rate: 4 l/min; l) capillary voltage (applied onto the sprayer): 3,500 V; m) reference (lock) masses: m/z 1,033.988109 for hexakis(1H, 1H, 3H-tetrafluoropropoxy)phosphazine, and m/z 112.985587 for trifluoroacetic acid; n) scanning range: 50-1,100 m/z; and o) scanning rate: 1.5 spectra/s. For cation mode, the capillary was pre-conditioned with 1 M NaOH for 30 min, followed by flushed with di-distilled water for 2 h, and then Cation Running Buffer (1 mol/l formic acid, freshly prepared) for another 2 h. Sample was separated as in the anion mode, except that the Cation Running Buffer was used, the capillary voltage was set at 3,500 V, and the fragmentor voltage 80 V. Data were collected using MassHunter LC/MS acquisition 10.1.48 (Agilent Technologies), and were processed using Qualitative Analysis B.06.00 (Agilent Technologies).

To analyse polar metabolites via GC-MS^104^, some 50 μl of each serum was instantly mixed with 200 μl of methanol containing 40 μg/ml tridecanoic acid and 10 µg/ml myristic-d27 acid as internal standards, followed with 20 s of vortexing and 30 min of incubation at −20 °C. The mixture was then centrifuged at 15,000*g* for 15 min at 4 °C, and 200 μl of supernatant (aqueous phase) was lyophilised at 4 °C for 24 h. The lyophilised sample was then vortexed for 1 min after mixing with 50 μl of freshly prepared methoxyamine hydrochloride (20 mg/ml in pyridine), followed by incubation at 4 °C for 1 h. The mixture was sonicated at 0 °C by bathing in ice slurry for 10 min, and was then incubated at 37 °C for 1.5 h, followed by mixing with 50 μl of MSTFA and incubated at 37 °C for 1 h. Before subjecting to GC-MS, samples were centrifuged at 15,000*g* for 10 min, and some 60 μl of each supernatant was loaded into an injection vial (cat. 5182-0714, Agilent; with an insert (cat. HM-1270, Zhejiang Hamag Technology)) equipped with a snap cap (cat. HM-0722, Zhejiang Hamag Technology). GC was performed on a HP-5MS column (30 m × 0.25 mm i.d., 0.25 μm film thickness) using a GC/MSD instrument (7890-5977B; Agilent Technologies). The injector temperature was set at 260 °C. The column oven temperature was first held at 70 °C for 2 min, then increased to 180 °C at the rate of 7 °C/min, then to 250 °C at the rate of 5 °C/min, then to 310 °C at the rate of 25 °C/min, where it was held for 15 min. The MSD transfer temperature was 280 °C. The MS quadrupole and source temperature were maintained at 150 °C and 230 °C, respectively. Data were collected using the MassHunter GC/MS Acquisition software (v.B.07.04.2260, Agilent Technologies) and were analysed using GC-MS MassHunter Workstation Qualitative Analysis software (v.10.1.733.0, Agilent Technologies).

Quantitative lipidomics were performed using the stable isotope dilution methods^160, 161^. Briefly, lipids were extracted from 50 µl of serum using a modified version of Bligh and Dyer’s method^162^ by mixing 750 µl of chloroform:methanol:MilliQ water(same hereafter for the lipidomics) (3:6:1) (v/v/v) with serum samples. After incubating at 1,500 r.p.m. for 1 h at 4 □ on a ThermoMixer C (Eppendorf), 350 µl of water and 250 µl of chloroform were added to the mixture to induce phase separation. After transferring the organic phase to a clean Eppendorf tube, the remaining lipid in the mixture was extracted again via the addition of another 450 μl of chloroform. The organic phase obtained from the two rounds of extraction was pooled and lyophilised using a SpeedVac Vacuum Concentrator (Genevac) under the OH mode. Samples were then dissolved in 100 µl of chloroform:methanol (1:1) (v/v) containing an Internal Standard Cocktail (see ref. ^162^). Lipidomic analyses were conducted at LipidALL Technologies using a Nexera 20-AD HPLC (Shimadzu) coupled with QTRAP 6500 PLUS MS (SCIEX), as described previously^161^. For polar lipids, a normal phase (NP)-HPLC was performed using a TUP-HB silica column (i.d. 150 × 2.1 mm, 3 µm; Tuplabs), and the gradient elution programme was: 2% B (chloroform:methanol:ammonium hydroxide:water, mixed at 55:39:0.5:5.5 (v/v); with the mobile phase A: chloroform:methanol:ammonium hydroxide, 89.5:10:0.5 (v/v)) held for 2 min, followed by three incremental increases: a) to 35% B at the 3^rd^ min, b) to 55% B at the 5^th^ min, and c) to 85% B at the 6^th^ min; and then maintained at 85% B for 1 min, followed by increasing to 100% B within 0.2 min and maintained for another 3.8 min, and finally decreased to 2% B within 0.5 min. The column was equilibrated at 2% B for 4.5 min between each run. Polar lipids were qualified by three separate injections under the ESI mode, with two injections in the positive mode at two separate dilutions (for PC, LPC, SM, Cer, GluCer, LacCer, and Sph; to guarantee that all polar lipids detected fall within the linear ranges of intensities), and one injection in the negative mode (for PE, PG, PI, PA, PS, BMP, CL, GM3, SL, FFA, LPE, LPI, LPA, LPS, and PC with fatty acyl-specific transitions). MS source parameters were set as follows: CUR 20, TEM 400 °C, GS1 20, and GS2 20. MRM transitions were set up for the quantitative analysis of polar lipids. Each polar lipid species were quantified by referencing to spiked internal standards, including d9-PC32:0(16:0/16:0), d9-PC36:1p(18:0p/18:1), d7-PE33:1(15:0/18:1), d9-PE36:1p(18:0p/18:1), d31-PS(d31-16:0/18:1), d7-PA33:1(15:0/18:1), d7-PG33:1(15:0/18:1), d7-PI33:1(15:0/18:1), C17-SL, Cer d18:1/15:0-d7, C12:0 Cer-1-P, d9-SM d18:1/18:1, C8-GluCer, C8-GalCer, d3-LacCer d18:1/16:0 Gb3 d18:1/17:0, d7-LPC18:1, d7-LPE18:1, C17-LPI, C17-LPA, C17-LPS, C17-LPG, d17:1 Sph, d17:1 S1P (Avanti Polar Lipids), GM3-d18:1/18:0-d3 (Matreya), d31-16:0 (Sigma), and d8-20:4 (Cayman Chemicals). For neutral lipids (TAGs and DAGs), a Kinetex-C18 column (i.d. 4.6 × 100 mm, 2.6 µm; Phenomenex), and an isocratic mobile phase containing chloroform:methanol:0.1 M ammonium acetate 100:100:4 (v/v/v) at a flow rate of 300 µl/min for 10 min were used. MS source parameters were set as described above, and the ESI-positive mode was used. Levels of short-, medium-, and long-chain TAGs were quantified by referencing to spiked internal standards of TAG(14:0)3-d5, TAG(16:0)3-d5 and TAG(18:0)3-d5 (CDN isotopes), while DAGs d5-DAG17:0/17:0 and d5-DAG18:1/18:1 (Avanti Polar Lipids). For free Cho and total cholesteryl esters, the method with atmospheric pressure chemical ionisation in the positive mode on a 1260 Infinity II HPLC (Agilent Technologies) coupled to a QTRAP 5500 MS (SCIEX), as established previously^163^, was used, during which lipids were separated on an Eclipse XDB C18 5-μm column (i.d. 150 × 4.6 mm; Agilent Technologies) using an isocratic mobile phase comprising chloroform:methanol (1:1 v/v) at a flow rate of 700 μl/min, and the MS source were set as follows: CUR 20, TEM 500 °C, GS1 45, and GS2 35. MS data was acquired and analysed using the Analyst 1.6.3 software (SCIEX). During the analysis, quality control (QC) samples, pooled from analysis samples, were inserted into the sample queue across every 10 biological samples.

### Measurement of adenylates and NAD^+^

ATP, ADP, AMP, and NAD^+^ from cells, tissues or flies were analysed by CE-MS as described in the “Determination of serum metabolome” section, except that cells collected from a 10-cm dish (60–70% confluence), 100 mg of liver or muscle tissue dissected by freeze clamp, or 20 anesthetised adult flies were used. Before CE-MS analysis, cells were rinsed with 20 ml of 5% (m/v) mannitol solution (dissolved in water) and instantly frozen in liquid nitrogen. Cells were then lysed with 1 ml of methanol containing IS1 (1:500 dilution), and were scraped from the dish. For analysis of metabolites in liver and muscle, mice were anaesthetised after indicated treatments. The tissue was then quickly excised by freeze-clamping, and then ground in 1 ml of methanol with IS1. For analysis of metabolites in flies, anesthetised flies were grounded in 1 ml of methanol with IS1 after freezing by liquid nitrogen. The lysate was then mixed with 1 ml of chloroform and 400 μl of water by 20 s of vortexing. After centrifugation at 15,000*g* for 15 min at 4 °C, 420 μl of aqueous phase was collected, filtrated, lyophilised, dissolved and subjected to CE-MS analysis in the negative mode as described in the “Determination of serum metabolome” section. Data were collected using MassHunter LC/MS acquisition 10.1.48 (Agilent Technologies), and were processed using Qualitative Analysis B.06.00 (Agilent Technologies). Levels of AMP, ADP, ATP and NAD^+^ were measured using full scan mode with m/z values of 346.0558, 426.0221, 505.9885 and 662.1019. Note that a portion of ADP and ATP could lose one phosphate group during in-source fragmentation, thus leaving the same m/z ratios as AMP and ADP, and should be corrected according to their different retention times in the capillary. Therefore, the total amount of ADP is the sum of the latter peak of the m/z 346.0558 spectrogramme and the former peak of the m/z 426.0221 spectrogramme, and the same is applied for ATP. Note that the retention time of each metabolite may vary between each run, and can be adjusted by isotope-labelled standards (dissolved in individual cell or tissue lysates) run between each samples, so do IS1 and IS2.

Levels of ATP, ADP, AMP and NAD^+^ in nematodes were analysed using the HPLC-MS as described in the “Determination of serum metabolome” section, except that some 150 nematodes maintained on NGM plates (containing 50 mM LCA or not) for 48 h were used. Nematodes were washed with ice-cold M9 buffer containing Triton X-100. Bacteria were removed by quickly spinning down the slurry at 100*g* for 5 s. Nematodes were then instantly lysed in 1 ml of methanol, then mixed with 1 ml of chloroform and 400 μl of water (containing 4 µg/ml [U-^13^C]-glutamine), followed by 20 s of vortexing. After centrifugation at 15,000*g* for another 15 min at 4 °C, 800 μl of aqueous phase was collected, lyophilised in a vacuum concentrator at 4 °C, and then dissolved in 30 μl of 50% (v/v, in water) acetonitrile. Metabolites were determined by HPLC-MS in the negative mode as described in the “Determination of serum metabolome” section. The following transitions (Q1/Q3) were used for monitoring each compound: 505.9/158.9 and 505.9/408.0 for ATP; 425.9/133.9, 425.9/158.8 and 425.9/328.0 for ADP; 345.9/79.9, 345.9/96.9 and 345.9/133.9 for AMP; 662.0/540.1 for NAD^+^; and 149.9/114 for [U-^13^C]-glutamine. Data were collected using Analyst software (v.1.7.1, SCIEX), and the relative amounts of metabolites were analysed using MultiQuant software (v.3.0.3, SCIEX). Similar to the CE-MS analysis, a portion of ADP and ATP could lose one or two phosphate groups during in-source-fragmentation thus leaving same m/z ratios as AMP and ADP, which was corrected according to their different retention times in column.

### Determination of bile acids concentrations

To measure the bile acid concentrations, MEFs (60-70% in 10-cm dish, rinsed with PBS and trypsinised), tissues (50 mg of liver or muscle tissues collected from each mouse after anaesthetising and blood-draining), nematodes (1,000 nematodes at L4 stage, washed with M9 buffer containing Triton X-100 for 3 times and frozen in liquid nitrogen) and flies (20 anaesthetised adults washed with PBS for 3 times and frozen in liquid nitrogen) rinsed PBS for 5 times, each in a fresh Eppendorf tube, or serum (50 μl collected from each mouse), were vigorously mixed (for serum) or homogenised (for others) with 1 ml of 80% methanol (v/v, in water) containing 100 μg/l CA-d5 and 4 µg/ml [U-^13^C]-glutamine as internal standards. After centrifugation at 15,000*g* for 15 min at 4 °C, some 30 µl of supernatant was loaded into an injection vial (cat. 5182-0714, Agilent Technologies; with an insert (cat. HM-1270, Zhejiang Hamag Technology)) equipped with a snap cap (cat. HM-2076, Zhejiang Hamag Technology), and 2 μl of supernatant was injected into an ACQUITY HSS T3 column (Waters). Measurement was performed on a QTRAP 6500 Plus MS (SCIEX) connected to an ACQUITY I-class UPLC system (Waters). The mobile phase consisted of 10 mM ammonium formate containing 0.005% formic acid (v/v) in the LC–MS-grade water (mobile phase A) and LC–MS-grade acetonitrile (mobile phase B) run at a flow rate of 0.4 ml/min. The HPLC gradient was as follows: 30% B for 1 min, then to 100% B at the 10^th^ min, hold for 2 min, and then back to 30% B, hold for another 3 min. The MS was run on a Turbo V ion source and running in negative mode run in a spray voltage of −5,500 V, with source temperature at 550 °C, gas NO.1 at 50 psi, gas NO.2 at 60 psi, curtain gas at 35 psi, and collision gas at “medium”. Compounds were measured using the MRM mode, and declustering potentials and collision energies were optimised using analytical standards. The following transitions (Q1/Q3) were used for monitoring each compound: 391.2/345.2 for deoxycholate; 407.2/345.2 for CA; 448.4/74.0 for glycochenodeoxycholate; 464.2/74.0 for glycocholate; 498.3/80.0 for taurodeoxycholate; 391.2/391.2 for ursodeoxycholate; 375.2/375.2 for LCA; 391.2/373.2 for CDCA; 482.2/80.0 for taurolithocholate; 498.2/79.9 for tauroursodeoxycholate; 448.4/74.0 for glycodeoxycholate; 514.3/79.9 for taurocholate; 498.3/80.0 for taurochenodeoxycholate; 407.3/387.3 for α-muricholate; 407.3/371.1 for β-muricholate; 407.3/405.3 for ω-muricholate; 514.3/123.9 for tauro-α-muricholate; 514.3/79.8 for tauro-β-muricholate; 149.9/114 for [U-^13^C]-glutamine; and 412.3/348.3 for CA-d5. For quantification of LCA, the LCA-d4 dissolved in individual lysates were used to generate corresponding standard curves by plotting the area ratios of labelled LCA (areas of LCA divided by the two ISs) against the added/actual concentrations of labelled LCA. The concentrations LCA were estimated according to standard curves. The average volume and density are 2,000 μm^3^ (ref. ^57^) and 1.1 g/ml (ref. ^58^), respectively for MEFs; 3,500 μm^3^ and 1.06 g/ml for mouse myocytes^59, 60^; and 1 × 10^6^ μm^3^ and 1.07 g/ml for nematodes^164, 165^. Data were collected using Analyst software (v.1.6.3, SCIEX), and the relative amounts of metabolites were analysed using MultiQuant software (v.3.0.2, SCIEX).

### Reagents

Rabbit anti-phospho-AMPKα-Thr172 (cat. #2535, RRID: AB_331250; 1:1,000 for IB), anti-AMPKα (cat. #2532, RRID: AB_330331; 1:1,000 for IB), anti-phospho-ACC-Ser79 (cat. #3661, RRID: AB_330337; 1:1,000 for IB), anti-ACC (cat. #3662, RRID: AB_2219400; 1:1,000 for IB), anti-GAPDH (cat. #5174, RRID: AB_10622025; 1:1,000 for IB), and anti-AXIN1 (cat. #2074, RRID: AB_2062419; 1:1,000 for IB) antibodies were purchased from Cell Signaling Technology. Rabbit anti-tubulin (cat. #10068-1-AP; RRID: AB_2303998; 1:1,000 for IB) was purchased from Proteintech. Mouse anti-total OXPHOS (cat. ab110413, RRID: AB_2629281; 1:5,000 for IB), and rabbit anti-Laminin (cat. ab11575, RRID: AB_298179; 1:200 for IF) antibodies were purchased from Abcam. Mouse anti-eMHC (cat. BF-G6, RRID: AB_10571455; 1:100 for IHC), anti-Pax7 (cat. Pax-7, RRID: AB_2299243; 1:100 for IHC), anti-MHCIIa (cat. SC71, RRID: AB_2147165; 1:100 for IHC), anti-MHCIIb (cat. BF-F3, RRID: AB_2266724; 1:100 for IHC), and anti-MHCI (cat. C6B12, RRID: AB_528351; 1:100 for IHC) antibodies were purchased from Developmental Studies Hybridoma Bank. Mouse anti β-ACTIN (cat. A5316, RRID: AB_476743; 1:1,000 for IB) was purchased from Sigma. Goat anti-Mouse IgM (Heavy chain) Cross-Adsorbed Secondary Antibody, Alexa Fluor 488 (cat. A-21042, RRID: AB_2535711; 1:200 for IHC), Goat anti-Mouse IgG2b Cross-Adsorbed Secondary Antibody, Alexa Fluor 594 (cat. A-21145, RRID: AB_2535781; 1:200 for IHC), Goat anti-Mouse IgG1 Cross-Adsorbed Secondary Antibody, Alexa Fluor 647 (cat. A-21240, RRID: AB_2535809; 1:200 for IHC), Goat anti-Mouse IgG1 Cross-Adsorbed Secondary Antibody, Alexa Fluor 488 (cat. A-21121, RRID: AB_2535764; 1:200 for IHC), and Goat anti-Rabbit IgG (H+L) Cross-Adsorbed Secondary Antibody, Alexa Fluor 594 (cat. A-11012, RRID: AB_2534079; 1:200 for IHC) were purchased from Thermo. The horseradish peroxidase (HRP)-conjugated goat anti-mouse IgG (cat. 115-035-003, RRID: AB_10015289; 1:5,000 dilution for IB) and goat anti-rabbit IgG (cat. 111-035-003, RRID: AB_2313567; 1:5,000 dilution for IB) antibodies were purchased from Jackson ImmunoResearch.

Information of supplier and catalogue numbers of metabolites for screening were listed in Supplementary Table 1. DMSO (cat. D2650), LCA (cat. L6250), (2-hydroxypropyl)-β-cyclodextrin (cat. C0926), NaCl (cat. S7653), CaCl_2_ (cat. C5670), MgSO_4_ (cat. M2643), H_2_O_2_ (cat. H1009), KH_2_PO_4_ (cat. P5655), K_2_HPO_4_ (cat. P9666), streptomycin (cat. 85886), cholesterol (cat. C3045), agar (cat. A1296), propionic acid (cat. P5561), sucrose (cat. S7903), glucose (cat. G7021), hematoxylin solution (cat. 51275), heparin (cat. H3149), 2-methylbutane (isopentane; cat. M32631), paraformaldehyde (cat. 158127), eosin Y-solution (cat. 318906), methanol (cat. 646377), ethanol (cat. 459836), chloroform (cat. C7559), PBS (cat. P5493), Triton X-100 (cat. T9284), xylene (cat. 534056), D-mannitol (cat. M4125), Canada balsam (cat. C1795), BSA (cat. A2153), glycerol (cat. G5516), Na_2_HPO_4_ (cat. S7907), sodium hypochlorite solution (NaClO; cat. 239305), NaOH (cat. S8045), Iron(II) sulfate heptahydrate (FeSO_4_; cat. F8633), isopropanol (cat. 34863), diethylpyrocarbonate (DEPC)-treated water (cat. 693520), paraquat (cat. 36541), HEPES (cat. H4034), EDTA (cat. E6758), EGTA (cat. E3889), MgCl_2_ (cat. M8266), KCl (cat. P9333), IGEPAL CA-630 (NP-40; cat. I3021), dithiothreitol (DTT; cat. 43815), collagenase B (cat. 11088831001), dispase II (cat. 4942078001), proteinase K (cat. P6556), agarose (cat. A9539), L-methionine sulfone (cat. M0876), D-campher-10-sulfonic acid (cat. 1087520), acetonitrile (cat. 34888), ammonium acetate (cat. 73594), ammonium hydroxide solution (cat. 338818), 3-aminopyrrolidine dihydrochloride (cat. 404624), N,N-diethyl-2-phenylacetamide (cat. 384011), trimesic acid (cat. 482749), diammonium hydrogen phosphate (cat. 1012070500), ammonium trifluoroacetate (cat. 56865), formic acid (cat. 5.43804), tridecanoic acid (cat. T0502), myristic-d27 acid (cat. 68698), methoxyamine hydrochloride (cat. 89803), hexane (cat. 34859), pyridine (cat. 270970), MSTFA (cat. M-132), cholic acid-2,2,3,4,4-d5 (CA-d5; cat. 614106), ammonium formate (cat. 70221), lithocholic acid-2,2,4,4-d4 (LCA-d4; cat. 589349), Trizma base (Tris; cat. T1503), sodium pyrophosphate (cat. P8135), β-glycerophosphate (cat. 50020), SDS (cat. 436143), sodium deoxycholate (cat. S1827), glutaraldehyde solution (cat. G5882), glycine (cat. G8898), K_3_Fe(CN)_6_ (cat. 455946), thiocarbonohydrazide (cat. 223220), Pb(NO_3_)_2_ (cat. 203580), acetone (cat. 534064), sodium citrate (cat. 71497), PD98059 (cat. P215), FCCP (cat. C2920), sodium azide (NaN_3_; cat. S2002), gentamycin (cat. 345814), collagenase A (cat. 11088793001), oligomycin A (cat. 75351), cardiotoxin (cat. 217503), Tween-20 (Cat. P9416), NaH_2_PO_4_ (Cat. S8282), and Ketone Body Assay kit (MAK134) were purchased from Sigma. 3-hydroxynaphthalene-2,7-disulfonic acid disodium salt (2-naphtol-3,6-disulfonic acid disodium salt; cat. H949580) was purchased from Toronto Research Chemicals. Hexakis(1H,1H,3H-perfluoropropoxy)phosphazene (hexakis(1H, 1H, 3H-tetrafluoropropoxy)phosphazine; cat. sc-263379) was purchased from Santa Cruz Biotechnology. Protease inhibitor cocktail (cat. 70221) was purchased from Roche. Seahorse XF base medium (cat. 103334) was purchased from Agilent Technologies. RNAprotect Tissue Reagent (cat. 76106) was purchased from Qiagen. OsO_4_ (cat. 18465) and uranyl acetate (cat. 19481) were purchased from Tedpella. SPI Low Viscosity “Spurr” Kit (cat. 02680-AB) was purchased from Structure Probe, Inc. Isolithocholic acid (iso-LCA; cat. 700195) was purchased from Avanti Polar Lipids. 3-oxo-5β-cholanoic acid (3-oxo-LCA; HY-125801), allolithocholic acid (allo-LCA; cat. HY-143712), isoallolithocholic acid (isoallo-LCA; HY-B0172A) and 3-oxoallo-LCA (customised) were purchased from MCE. Free Glycerol Assay kit (cat. ab65337) and Glycogen Assay kit (cat. ab65620) were purchased from Abcam. Bacteriological peptone (cat. LP0037) and yeast extract (cat. LP0021) were purchased from Oxoid. Insulin (Novolin R) was purchased from Novo Nordisk. NEBNext Poly(A) mRNA Magnetic Isolation Module (E7490), and NEBNext Ultra RNA Library Prep Kit for Illumina (cat. E7530) were purchased from NEB. AMPure XP Beads (cat. 63881) was purchased from Beckman Coulter. TruSeq PE Cluster Kit v3-cBot-HS kit (cat. PE-401-3001) was purchased from Illumina. Biospin Tissue Genomic DNA extraction Kit (cat. BSC04M1) was purchased from BioFlux. DMEM-high glucose (cat. 12800082), Liver Perfusion Medium (cat. 17701038), Liver Digest Medium (cat. 17703034), William’s E Medium (cat. 12551032), Ham’s F-10 medium (cat. 11550043), basic fibroblast growth factor (bFGF; cat. 13256-029), Schneider’s *Drosophila* Medium (cat. 21720024), DMEM containing HEPES (cat. 21063029), MEM non-essential amino acids solution (cat. 11140050), GlutaMAX (cat. 35050061), sodium pyruvate (cat. 11360070), Maxima SYBR Green/ROX qPCR Master Mix (cat. K0223), Trypan Blue Stain (cat. T10282), FBS (cat. 10099141C), Fluo-4-AM (cat. F14217), ProLong Live Antifade reagent (cat. P36975), penicillin-streptomycin (cat. 15140163), Prestained Protein MW Marker (cat. 26612), BCA Protein Assay Kit (cat. A55865), Maxima SYBR Green/ROX qPCR master mix (cat. K0223), and TRIzol (cat. 15596018) were purchased from Thermo. ReverTra Ace qPCR RT Master Mix with gDNA Remover (cat. FSQ-301) was purchased from Toyobo. WesternBright ECL and peroxide solutions (cat. 210414-73) were purchased from Advansta. Dry yeast (cat. FLY804020F) and cornmeal (cat. FLY801020) were purchased from LabScientific. Soy flour (cat. 62116) was purchased from Genesee. Light corn syrup was purchased from Karo. O.C.T. Compound (cat. 4583) was purchased from Sakura. Mouse Ultrasensitive Insulin ELISA kit (80-INSMSU-E10) was purchased from ALPCO. LabAssay Triglyceride Kit (cat. LABTRIG-M1) and LabAssay NEFA Kit (cat. LABNEFA-M1) were purchased from Wako. Mouse Glucagon ELISA Kit (cat. 81518) was purchased from Crystal Chem. [U-^13^C]-glucose (cat. CLM-1396-PK) and [U-^13^C]-glutamine (cat. CLM-1822-H-PK) were purchased from Cambridge Isotope Laboratories.

### Cell lines

In this study, no cell line used is on the list of known misidentified cell lines maintained by the International Cell Line Authentication Committee (https://iclac.org/databases/cross-contaminations/). HEK293T cells (cat. CRL-3216) and *Drosophila* Schneider 2 (S2) cells (cat. CRL-1963) were purchased from ATCC. MEFs were established by introducing SV40 T antigen via lentivirus into cultured primary embryonic cells from mouse litters as described previously^119^. HEK293T cells and MEFs were maintained in DMEM supplemented with 10% FBS, 100 IU penicillin, 100 mg/ml streptomycin at 37 °C in a humidified incubator containing 5% CO_2_. S2 cells were cultured in Schneider’s *Drosophila* Medium supplemented with 10% heat-inactivated FBS and 100 IU penicillin, 100 mg/ml streptomycin at 37 °C in a humidified incubator containing 5% CO_2_. All cell lines were verified to be free of mycoplasma contamination. HEK293T cells were authenticated by STR sequencing.

### Immunoblotting

To analyse the levels of p-AMPKα and p-ACC in HEK293T cells, MEFs, S2 cells, mouse primary hepatocytes and primary myocytes, cells grown to 70-80% confluence in a well of a 6-well dish were lysed with 250 μl of ice-cold Triton lysis buffer (20 mM Tris-HCl, pH 7.5, 150 mM NaCl, 1 mM EDTA, 1 mM EGTA, 1% (v/v) Triton X-100, 2.5 mM sodium pyrophosphate, 1 mM β-glycerophosphate, with protease inhibitor cocktail). The lysates were then centrifuged at 20,000*g* for 10 min at 4 °C and an equal volume of 2× SDS sample buffer was added into the supernatant. Samples were then boiled for 10 min and then directly subjected to immunoblotting. To analyse the levels of p-AMPKα and p-ACC in muscle and liver tissues, mice were anesthetised after indicated treatments. Freeze-clamped tissues were immediately lysed with ice-cold Triton lysis buffer (10 μl/mg tissue weight for liver, and 5 μl/mg tissue weight for muscle), followed by homogenisation and centrifugation as described above. The lysates were then mixed with 2× SDS sample buffer, boiled, and subjected to immunoblotting. To analyse the levels of p-AMPKα, p-ACC in flies, 20 adults or third instar larvae were lysed with 200 μl ice-cold RIPA buffer (50 mM Tris-HCl, pH 7.5, 150 mM NaCl, 1% NP-40, 0.5% sodium deoxycholate, with protease inhibitor cocktail) containing 0.1% SDS, followed by homogenisation and centrifugation as described above. The lysates were then mixed with 5× SDS sample buffer, boiled, and subjected to immunoblotting. To analyse the levels of p-AMPKα and p-ACC in nematodes, some 150 nematodes cultured on the NGM plate were collected for each sample. Worms were quickly washed with ice-cold M9 buffer containing Triton X-100, and were lysed with 150 μl of ice-cold lysis buffer. The lysates were then mixed with 5× SDS sample buffer, followed by homogenisation and centrifugation as described above, and then boiled before being subjected to IB. All samples were subjected to IB on the same day of preparation, and any freeze–thaw cycles were avoided.

For IB, the SDS-polyacrylamide gels were prepared in house as described previously^107^. The thickness of gels used in this study was 1.0 mm. Samples of less than 10 μl were loaded into wells, and the electrophoresis was run at 100 V (by PowerPac HC High-Current Power Supply, Bio-Rad) in a Mini-PROTEAN Tetra Electrophoresis Cell (Bio-Rad). In this study, all samples were resolved on 8% resolving gels, except those for OXPHOS proteins, which were on 15% gels (prepared as those for 8%, except that a final concentration of 15% Acryl/Bis was added to the resolving gel solution), and β-ACTIN which was on 10% gels. The resolved proteins were then transferred to the PVDF membrane (0.45 μm, cat. IPVH00010, Merck) as described previously^107^. The PVDF membrane was then blocked by 5% (w/v) BSA (for all antibodies against phosphorylated proteins) or 5% (w/v) non-fat milk (for all antibodies against total proteins) dissolved in TBST for 2 h on an orbital shaker at 60 r.p.m. at room temperature, followed by rinsing with TBST for twice, 5 min each. The PVDF membrane was then incubated with desired primary antibody overnight at 4 °C on an orbital shaker at 60 r.p.m., followed by rinsing with TBST for three times, 5 min each at room temperature, and then the secondary antibodies for 3 h at room temperature with gentle shaking. The secondary antibody was then removed, and the PVDF membrane was further washed with TBST for 3 times, 5 min each at room temperature. PVDF membrane was incubated in ECL mixture (by mixing equal volumes of ECL solution and Peroxide solution for 5 min), then life with Medical X-Ray Film (FUJIFILM). The films were then developed with X-OMAT MX Developer (Carestream), and X-OMAT MX Fixer and Replenisher solutions (Carestream) on a Medical X-Ray Processor (Carestream) using Developer (Model 002, Carestream). The developed films were scanned using a Perfection V850 Pro scanner (Epson) with an Epson Scan software (v.3.9.3.4), and were cropped using Photoshop 2023 software (Adobe). Levels of total proteins and phosphorylated proteins were analysed on separate gels, and representative immunoblots are shown. Uncropped immunoblots are shown in Supplementary Fig. 1.

### TEM imaging

The TEM imaging was performed based on the in situ embedding and sectioning method as described previously^166^, with minor modifications. Briefly, tibialis anterior muscle was quickly excised and sliced to around 1 mm × 1 mm × 5 mm cubes, followed by quickly immersed in 1 ml of 2.5% (v/v) glutaraldehyde solution (freshly prepared by diluting 25% (v/v) glutaraldehyde in 0.1 M Phosphate Buffer (by mixing 0.2 M Na_2_HPO_4_ with 0.2 M NaH_2_PO_4_ (both dissolved in water, and adjusted pH to 7.4) at a ratio of 81:19, and then diluted with equal volume of water) at 4 °C for 12 h. Muscles were then washed with 1 ml of 0.1 M Phosphate Buffer for 3 times at 4 °C, 20 min each, followed by staining with 1% (w/v) OsO_4_ solution (in 0.1 M Phosphate Buffer, supplemented with 1.5% K_3_Fe(CN)_6_) at 4 °C for 2 h, and then washed by for 5 times, 10 min each with ice-cold di-distilled water. Muscles were stained in ice-cold 2% (w/v, in water) uranyl acetate solution for 12 h at 4 °C in the dark, and were then washed for 4 times, 10 min each, with ice-cold water. Dehydration was then performed by sequentially incubating muscles in the following solutions: 30, 50, and 70% (v/v) ethanol (in water), each for 12 min at 4 °C, followed by incubating in 90, 100 and 100% (v/v) ethanol (in water), each for 12 min at room temperature, and then in acetone twice at room temperature. Muscles were then quickly submerged in acetone/Spurr resin (3:1; the Spurr resin was prepared by mixing with 15 g of NSA, 7.3 g of DER 736, 7.5 g of ERL 4206 with 320 μl of DMAE, all supplied in the SPI Low Viscosity “Spurr” Kits, for 1.5 h at room temperature) mixture at room temperature for a 1-h incubation, and then in acetone/resin (1:1) mixture at room temperature for 2 h, followed by acetone/resin (1:3) at room temperature for 2 h, and finally 100% resin at room temperature for 12 h. Resin was then completely drained, and the tissues were baked at a hot-wind drying oven at 70 °C. After baking for 24 h, the tissues were then sectioned into 70-nm slices on an EM UC7 Ultramicrotome (Leica) after cooling down to room temperature. Sections were then stained with lead citrate solution (by dissolving 1.33 g of Pb(NO_3_)_2_ and 1.76 g of sodium citrate in 42 ml of di-distilled water, followed by addition of 8 ml of 1 M NaOH) for 5 min at room temperature before imaging using an AMT-XR81DIR camera on an electron microscope (HT-7800, Hitachi) using TEM system control software (Ver. 01.20, Hitachi).

### Determination of the intracellular Ca^2+^ levels

Intracellular Ca^2+^ levels in MEFs treated with LCA were determined as described previously^102^. Briefly, cells were loaded with 5 μM (final concentration) Fluo-4-AM for 30 min, then washed twice with PBS and incubated in fresh, desired medium for another 30 min. Before image taking, ProLong Live Antifade Reagent was added to the medium. During imaging, live cells were kept at 37°C, 5% CO_2_ in a humidified incubation chamber (Incubator PM S1; Zeiss). Images were taken on an LSM980 microscope (Zeiss). Images were processed and analysed on Zen 3.4 software (Zeiss), and formatted on Photoshop 2023 software (Adobe).

### Determination of oxygen consumption rates

The OCR of nematodes was measured as described previously^167^. Briefly, nematodes were washed with M9 buffer for 3 times. Some 15 to 25 nematodes were then suspended in 200 μl of M9 buffer, and were added to a well on a 96-well Seahorse XF Cell Culture Microplate. The measurement was performed in Seahorse XFe96 Analyzer (Agilent Technologies) at 20 °C following the manufacturer’s instruction, with a Seahorse XFe96 sensor cartridge (Agilent Technologies) pre-equilibrated in Seahorse XF Calibrant solution in a CO_2_-free incubator at 37 °C overnight. Concentrations of respiratory chain inhibitors used during the assay were: FCCP at 10 μM and sodium azide at 40 mM. At the end of the assay, the exact number of nematode in each well was determined on a Cell Imaging Multi-Mode Reader (Cytation 1, BioTek) and was used for normalising/correcting OCR results. Data were collected using Wave 2.6.1 Desktop software (Agilent Technologies) and exported to Prism 9 (GraphPad) for further analysis according to manufacturer’s instructions.

The OCR of intact muscle tissue was measured as described previously^115, 168^, with modifications. In brief, mice were starved for desired durations, and were sacrificed through cervical dislocation. The gastrocnemius muscles from two hindlegs were then excised, followed by incubating in 4 ml of dissociation media (DM; by dissolving 50 μg/ml gentamycin, 2% (v/v) FBS, 4 mg/ml collagenase A in DMEM containing HEPES) in a 35-mm culture dish in a humidified chamber at 37 °C, 5% CO_2_, for 1.5 h. The digested muscle masses were then washed with 4 ml of pre-warmed collagenase A-free DM, incubated in 0.5 ml of pre-warmed collagenase A-free DM, and dispersed by passing through a 20 G needle for 6 times. Some 20 μl of muscle homogenates was transferred to a well of a Seahorse XF24 Islet Capture Microplate (Agilent Technologies). After placing an islet capture screen by a Seahorse Capture Screen Insert Tool (Agilent Technologies) into the well, 480 μl of pre-warmed aCSF medium (120 mM NaCl, 3.5 mM KCl, 1.3 mM CaCl_2_, 0.4 mM KH_2_PO_4_, 1 mM MgCl_2_, 5 mM HEPES, 15 mM glucose, 1× MEM non-essential amino acids, 1 mM sodium pyruvate, and 1 mM GlutaMAX; adjust to pH 7.4 before use) was added, followed by equilibrating in a CO_2_-free incubator at 37 °C for 1 h. OCR was then measured at 37 °C in an XFe24 Extracellular Flux Analyzer (Agilent Technologies), with a Seahorse XFe24 sensor cartridge (Agilent Technologies) pre-equilibrated in Seahorse XF Calibrant solution (Agilent Technologies) in a CO_2_-free incubator at 37 °C overnight. Respiratory chain inhibitors used during the assay were oligomycin at 100 μM, FCCP at 100 μM, 50 μM antimycin A, 1 μM rotenone (all final concentrations). Data were collected using Wave 2.6.3 Desktop software (Agilent Technologies) and exported to Prism 9 (GraphPad) for further analysis according to the manufacturer’s instructions.

### Statistical analysis

Statistical analyses were performed using Prism 9 (GraphPad Software), except for the survival curves, which were analysed using SPSS 27.0 (IBM). Each group of data was subjected to Kolmogorov-Smirnov test, Anderson-Darling test, D’Agostino-Pearson omnibus test or Shapiro-Wilk test for normal distribution when applicable. An unpaired two-sided Student’s *t*-test was used to determine significance between two groups of normally distributed data. Welch’s correction was used for groups with unequal variances. An unpaired two-sided Mann-Whitney test was used to determine significance between data without a normal distribution. For comparison between multiple groups with two fixed factors, an ordinary two-way ANOVA or two-way repeated-measures (RM) ANOVA (for blood glucose measured during GTT, ITT and clamping) was used, followed by Tukey’s or Sidak’s multiple comparisons test as specified in the legends. The assumptions of homogeneity of error variances were tested using F-test (*P* > 0.05). Geisser-Greenhouse’s correction was used where applicable. The adjusted means and s.e.m., or s.d., were recorded when the analysis met the above standards. Differences were considered significant when *P* < 0.05, or *P* > 0.05 with large differences of observed effects (as suggested in refs. ^169, 170^).

### Data availability

The data supporting the findings of this study are available within the paper and its Supplementary Information files. Materials, reagents or other experimental data are available upon request. Full immunoblots are provided as Supplementary Information Fig. 1. Source data are provided with this paper.

### Code availability

The analysis was performed using standard protocols with previously described computational tools. No custom code was used in this study.

## Acknowledgements

We thank Dr. Sean Morrison (University of Texas Southwestern Medical Center) for providing the *AMPK*α*1*^F/F^ (The Jackson Laboratory, 014141), and *AMPK*α*2*^F/F^ (The Jackson Laboratory, 014142) mice; Mingliang Zhang (Shanghai Jiao Tong University Affiliated Sixth People’s Hospital) for the technical assistance of hyperinsulinemia euglycemic clamp; Su-Qin Wu, Ying He and Jing Song (Xiamen University) for mouse in vitro fertilisation; Yong Yu (Xiamen University) and Ying Liu (Peking University) for the nematode strains and experiments; Bo Liu and Kewei Zheng (Xiamen University) for the fly strains and experiments; Qiqi Guo from Zhenji Gan laboratory (Nanjing University) for the helps on the analysis of fibre types of muscle tissues; Tong-Jin Zhao and Hua Bian (Fudan University) for the critical discussion on the manuscript; Xuan Guo (Jinzhou Medical University) for importing the fly strains; and all the other members of the S.-C.L. laboratory for technical assistance. We also acknowledge the *Caenorhabditis* Genetics Center and National BioResource Project for supplying nematode strains, and Bloomington *Drosophila* Stock Center, Vienna *Drosophila* Resource Center, and the Core Facility of *Drosophila* Resource and Technology, Center for Excellence in Molecular Cell Science, Chinese Academy of Sciences for providing fly strains and reagents. This work was supported by grants from the National Key R&D Program of China (2020YFA0803402 and 2022YFA0806501), the National Natural Science Foundation of China (#92057204, #82088102, #32070753, #31900542, #91854208 and #31922034), the Fundamental Research Funds for the Central Universities (#20720200069 and #20720190101), the Project “111” sponsored by the State Bureau of Foreign Experts and Ministry of Education of China (#BP2018017), the Joint Funds for the Innovation of Science and Technology, Fujian province (2021Y9232 and 2021Y9227), the Fujian provincial health technology project (2022ZD01005 and 2022ZQNZD009), the XMU Open Innovation Fund and Training Programme of Innovation and Entrepreneurship for Undergraduates (KFJJ-202103 and S202210384682), and the Agilent Applications and Core Technology - University Research Grant (#4769).

## Author contributions

Q.Q., C.-S.Z. and S.-C.L. conceived the study and designed the experiments. Q.Q., W.W. and H.-Y.Y. performed mice CR and LCA administrations. Q.Q. screened the metabolites responsible for AMPK activation, with the assistance from S.L., H.-Y.Y., and X.T. Y.C. determined the benefits of LCA on mice and flies, with the assistance from Q.Q., M.L., W.W. and H.-Y.Y. Y.W. performed nematode experiments. X.W. determined the OCR of mouse muscles. Y.-H.L., S.X. and Z.-Z.Z. determined the mRNA levels of mitochondrial OXPHOS complex. C.Z. conducted an analysis of polar metabolites in serum, tissues, and cells utilising HPLC-MS, H.-L.P. CE-MS, and M.Z. GC-MS. S.M.L. and G.S. performed lipidomic analyses. B.Z. and X.D. designed the formulation of LCA for mouse administration. C.-S.Z. and S.-C.L. wrote the manuscript.

## Additional information

**Supplementary information** The online version contains supplementary material available at https://doi.org/10.1038/.

**Correspondence and requests for materials** should be addressed to Sheng-Cai Lin.

**Peer review information** Nature thanks anonymous reviewer(s) for their contribution to the peer review of this work. Peer reviewer reports are available.

**Reprints and permissions information** is available at http://www.nature.com/reprints.

