## Supplementary material for "Lithocholic acid phenocopies rejuvenating and life-extending effects of calorie restriction": Uncropped gel images

**Fig. 1a**

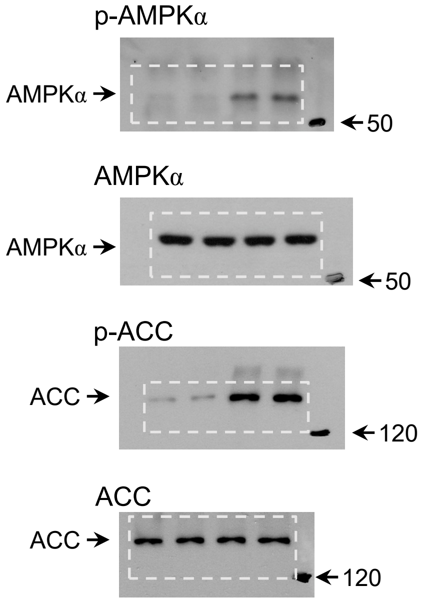


**Fig. 1b**

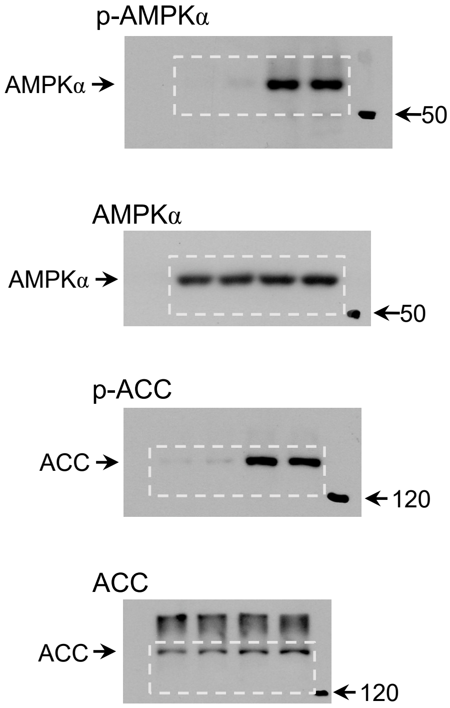


**Fig. 1c**

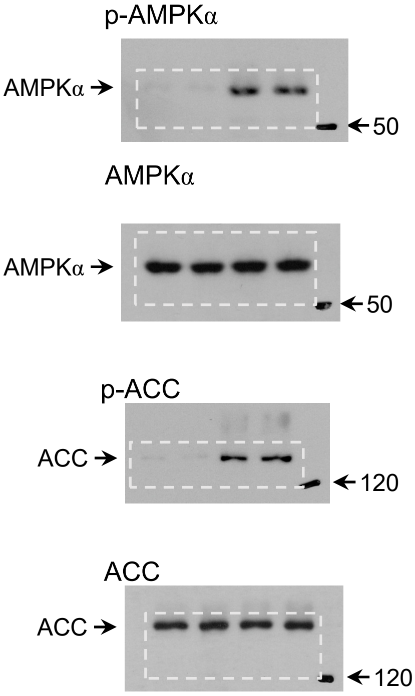


**Fig. 1d**

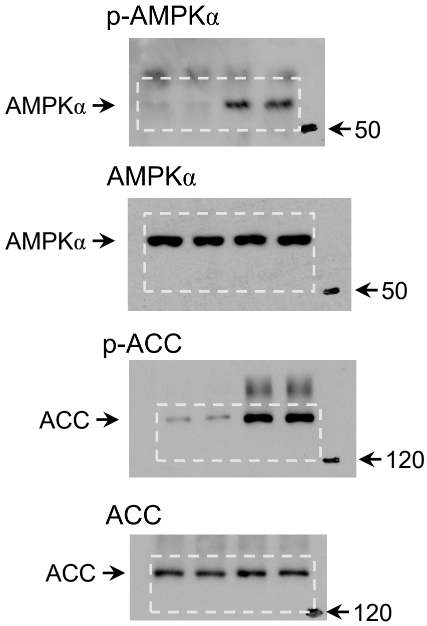

### Fig. 1e

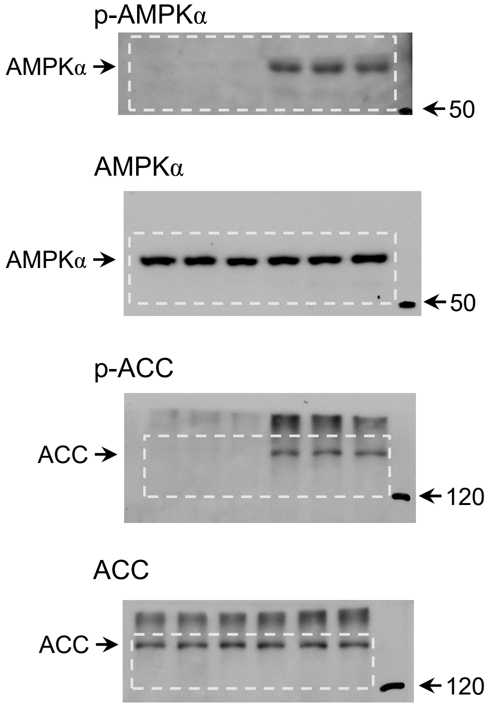


### Fig. 1f

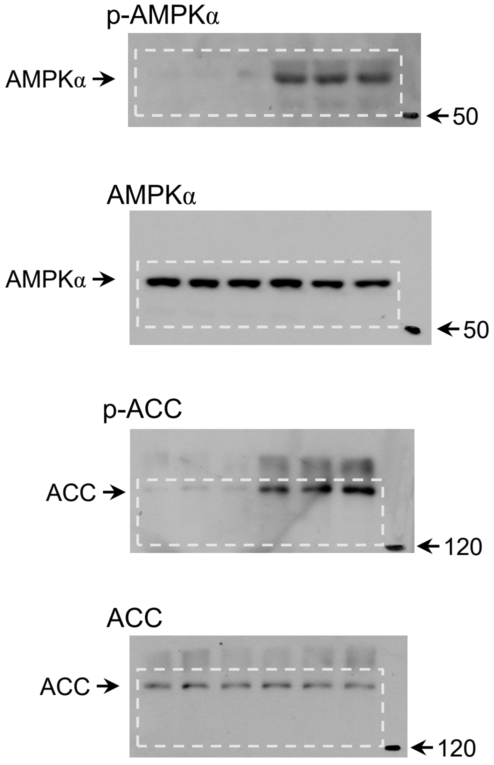


### Fig. 1h

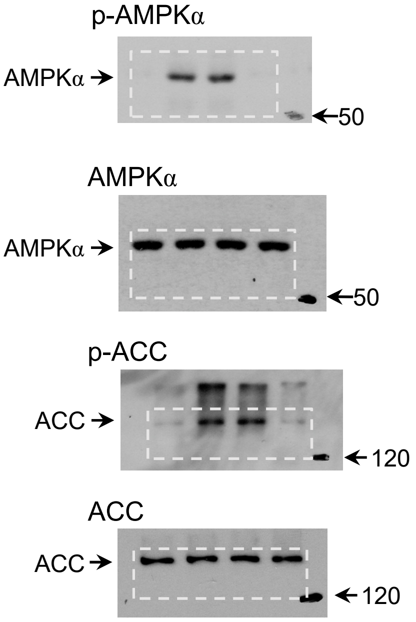


### Fig. 1i

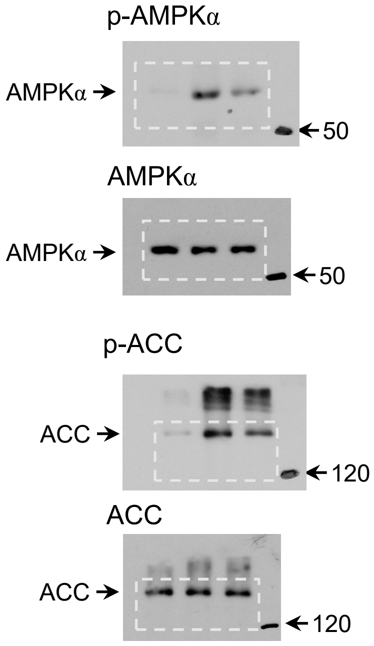

### Fig. 2b

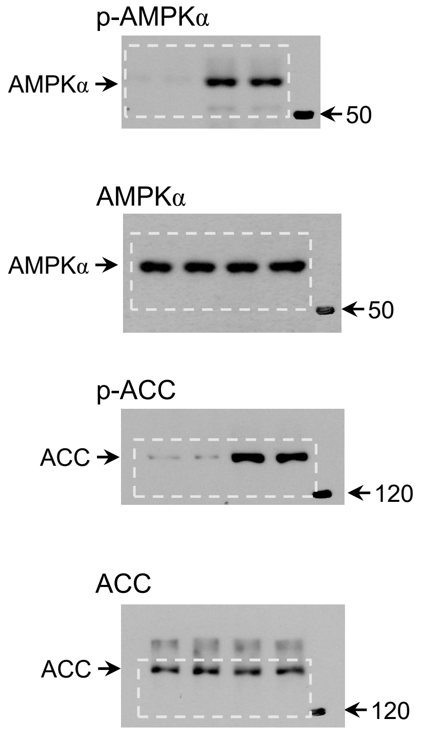


### Fig. 2c

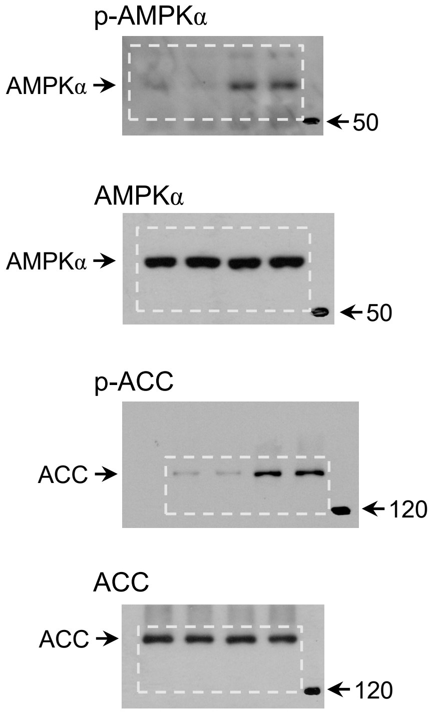


### Fig. 2i

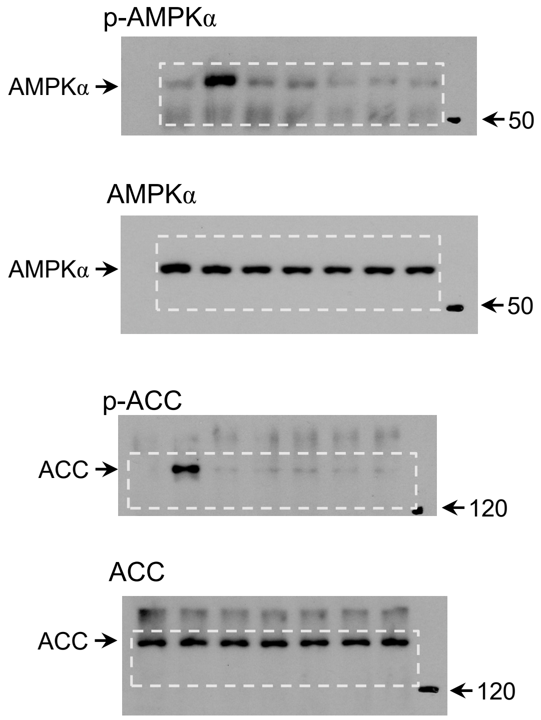

### Fig. 2l (upper)

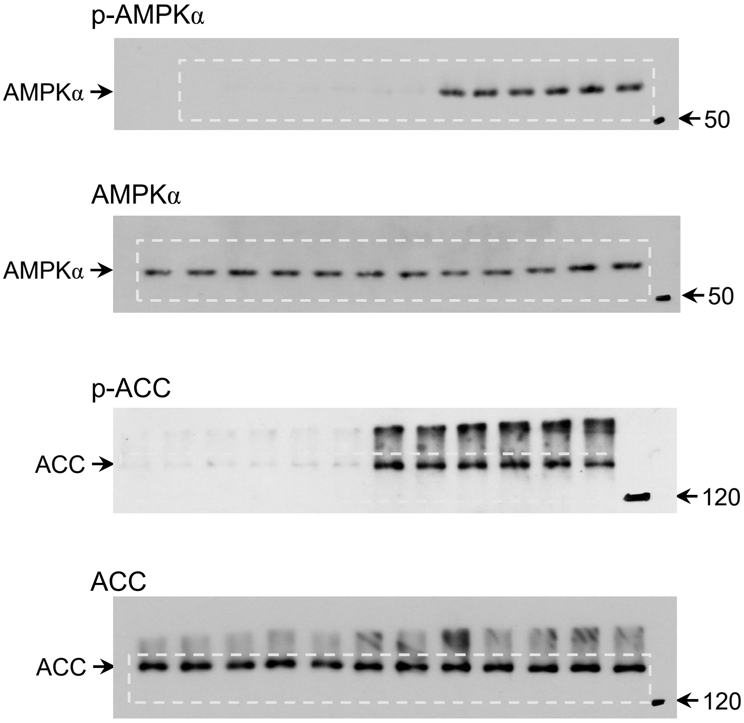


### Fig. 2m

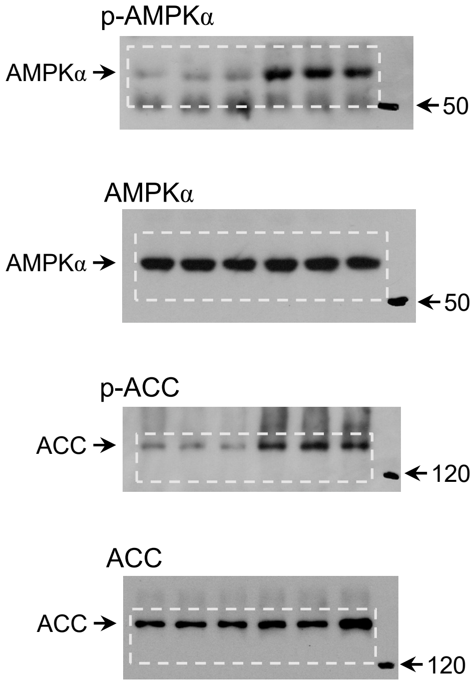


### Fig. 2l (lower)

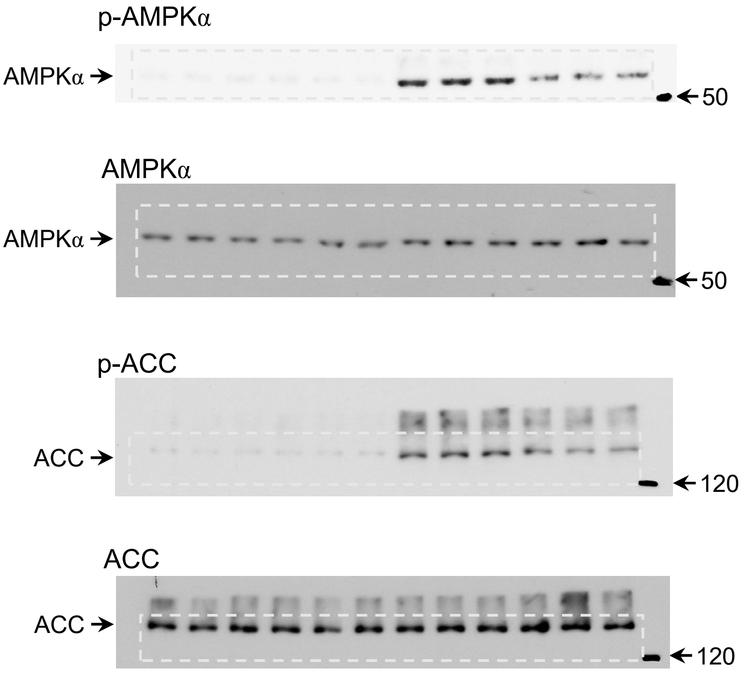

**Fig. 3g**

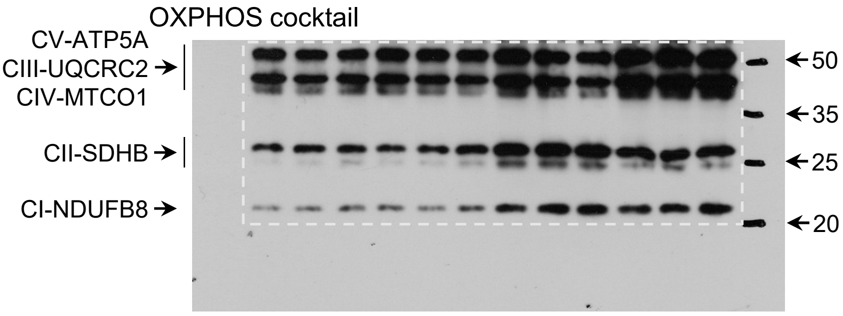
OXPHOS cocktail
CV-ATP5A
CIII-UQCRC2
CIV-MTCO1
CII-SDHB
CI-NDUFB8
50
35
25
20

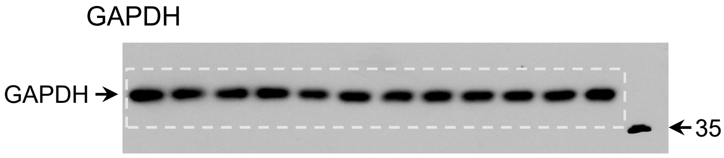
GAPDH
GAPDH
35

### Extended Data Fig. 1a

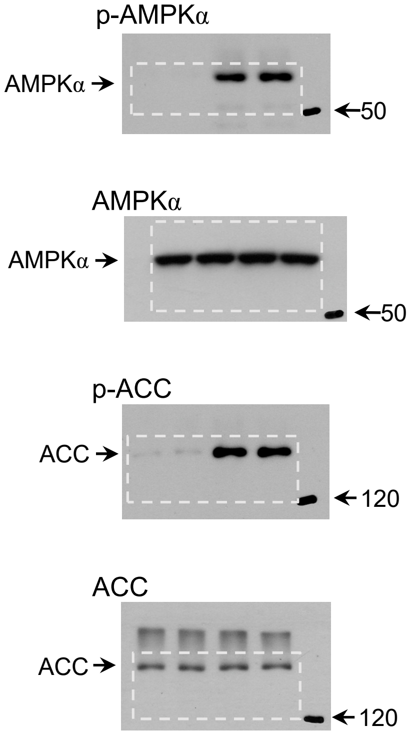


### Extended Data Fig. 1b

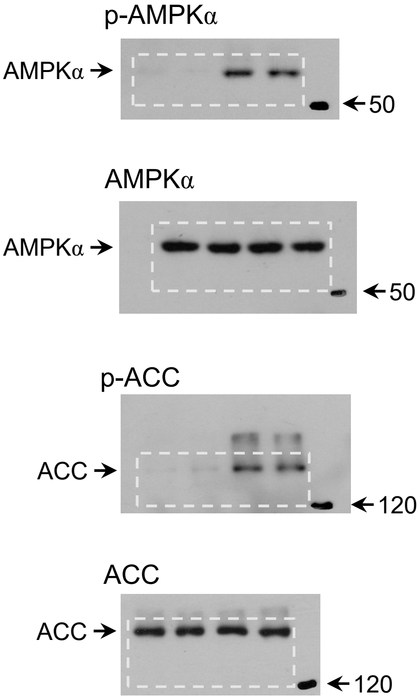


### Extended Data Fig. 1d

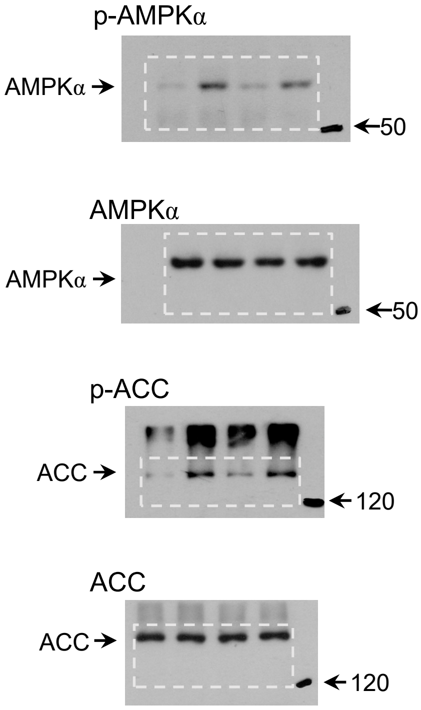

### Extended Data Fig. 5a

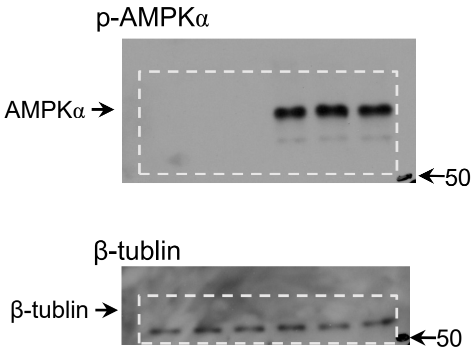


### Extended Data Fig. 5b

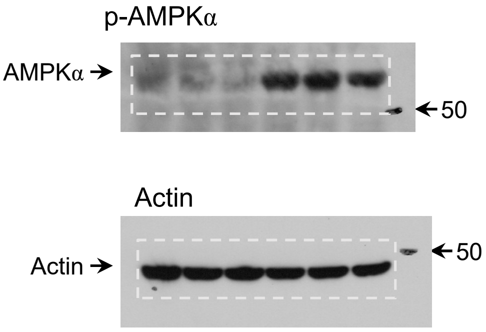


### Extended Data Fig. 5c

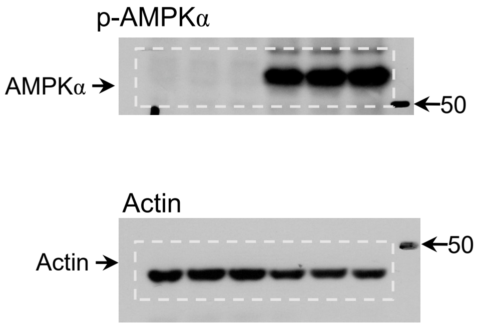


### Extended Data Fig. 5d

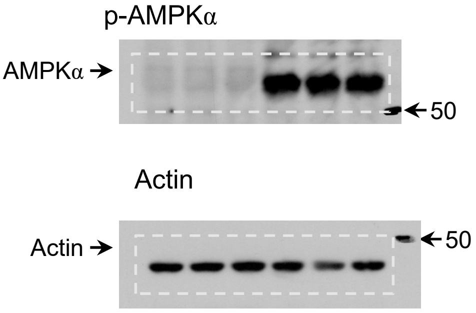
