## Supplementary material for "Lithocholic acid phenocopies rejuvenating and life-extending effects of calorie restriction": ED figures 1-5 and ED tables 1-2

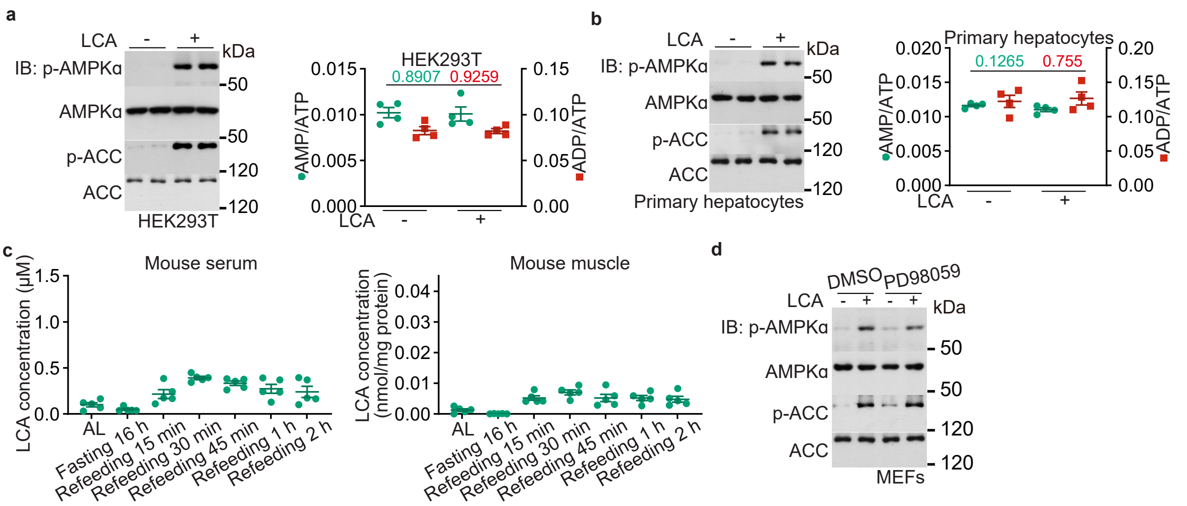


**Extended Data Fig. 1 | LCA activates AMPK at a level seen in CR serum**.

**a**, **b**, LCA at concentrations similar to that in the serum of CR mice activates AMPK in HEK293T cells and primary hepatocytes. HEK293T cells (**a**) and primary hepatocytes (**b**) were treated with 1 μM LCA for 4 h, followed by determining the activation of AMPK, and the AMP:ADP and ADP:ATP ratios (results are mean ± s.e.m.; *n* = 4 samples for each treatment, and *P* value by two-sided Student's *t*-test).

**c**, Ad libitum-fed mouse serum and muscle contain little LCA, with levels increased after refeeding. The ad libitum-fed mice were fasted for 8 h and re-fed. LCA concentrations in serum and muscle at different time points following refeeding were determined. Results are shown as mean ± s.e.m.; *n* = 5 mice for each time point.

**d**, LCA does not activate AMPK through the cAMP-Epac-MEK pathway. MEFs were treated with 1 μM LCA with or without 100 μM PD98059 for 4 h, followed by determining the activation of AMPK by immunoblotting.

Experiments in this figure were performed three times.

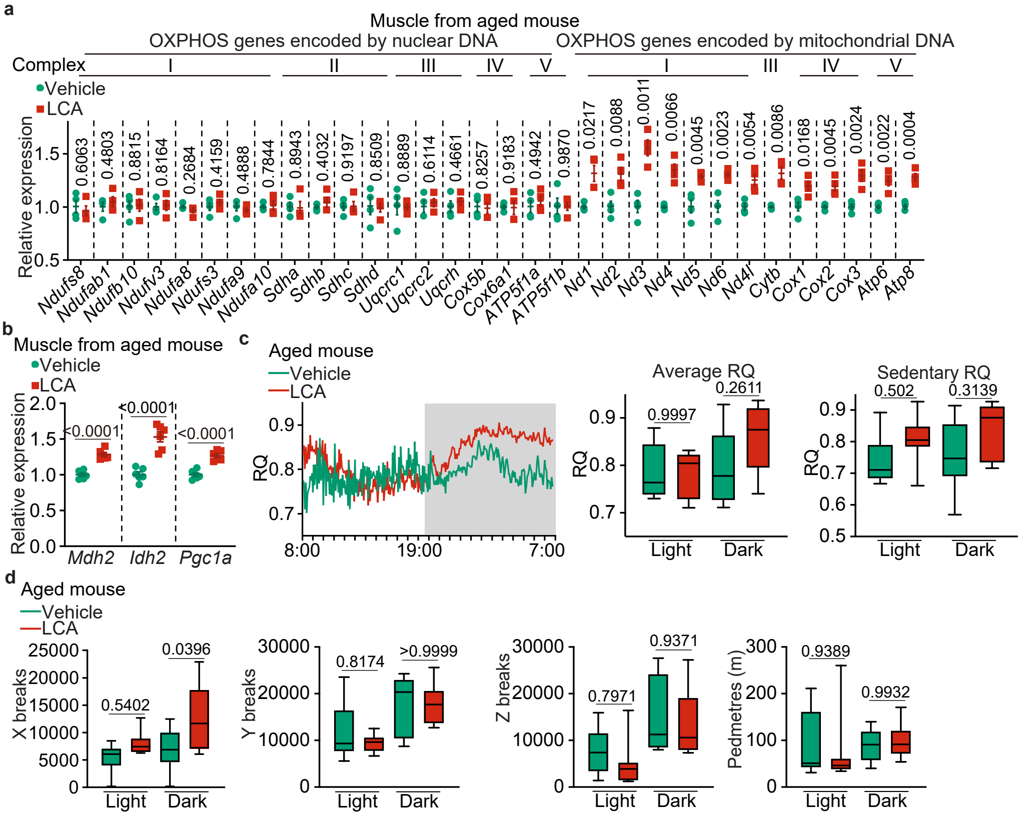


**Extended Data Fig. 2 | LCA retard ageing in mice.**

**a**, **b**, LCA elevates mitochondrial contents in the muscles of aged mice. Mice were treated as in Fig. 3a, followed by determining the mRNA levels of OXPHOS (**a**) and TCA cycle (**b**) genes. Results are mean ± s.e.m., normalised to the vehicle group; $n = 4$ mice for each treatment, and $P$ value by two-sided Student's $t$-test.

**c**, **d**, LCA elevates RQ in aged mice. Mice were treated as in Fig. 3a, followed by determining RQ (**c**) and ambulatory activity (**d**). Data are shown as mean (left panel of **c**; at 5-min intervals during a 24-h course), or as box-and-whisker plots (middle and right panels of **c**; and **d**); $n = 6$ (vehicle, light cycle of sedentary RQ), 7 (LCA, dark cycle of sedentary RQ), or 8 (others) mice for each treatment, and $P$ value by two-way ANOVA followed by Tukey's test.

Experiments in this figure were performed three times.

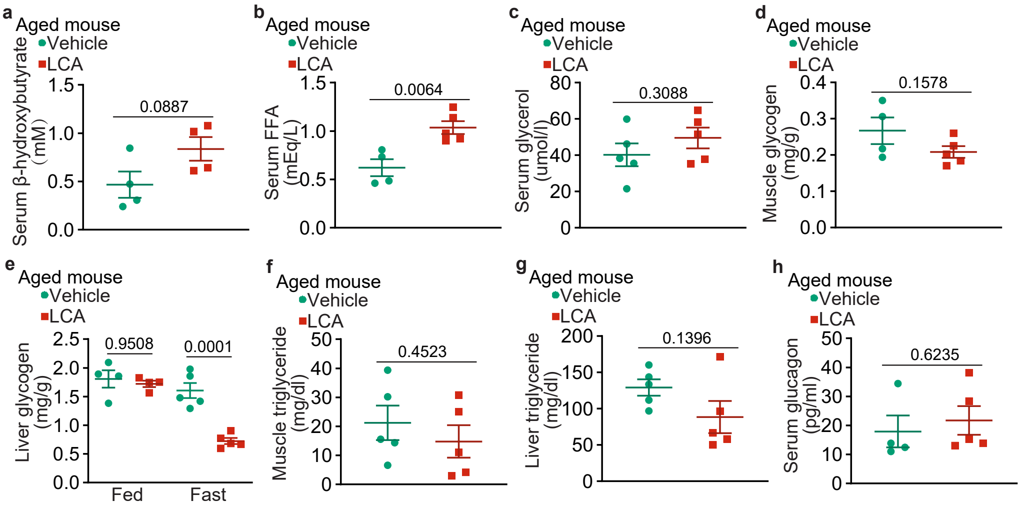


**Extended Data Fig. 3 | LCA ameliorates age-related insulin resistance without decreasing glucose production.**
**a**-**h**, Mice were treated as in Fig. 3a, followed by an 8 h-fasting period (except for liver glycogen, in which both the feeding and fasting mice were used). The carbon sources responsible for glucose production, including serum β-hydroxybutyrate (**a**), serum free fatty acids (**b**), serum glycerol (**c**), muscle glycogen and liver glycogen (**d**, **e**), and muscle and liver triglyceride (**f**, **g**), along with serum glucagon (**h**), were determined. Results are mean ± s.e.m.; $n$ = 4 (**a**, vehicle group of **b**, feeding group of **e**, and vehicle group of **h**) or 5 (others) mice for each treatment, and $P$ value by two-way ANOVA followed by Tukey's test (**e**), or by two-sided Student's $t$-test (others). Experiments in this figure were performed three times.

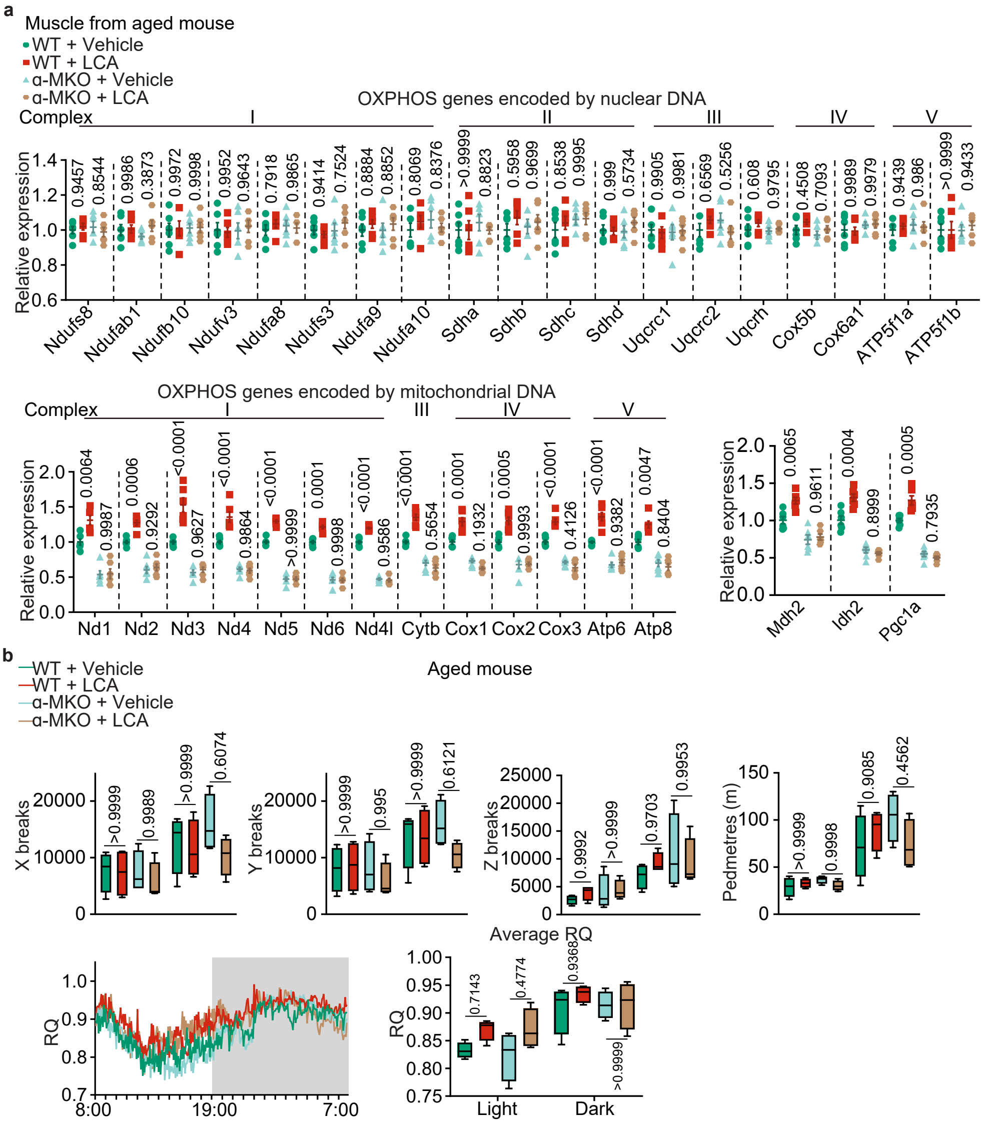


**Extended Data Fig. 4 | LCA exerts the rejuvenating activity through activating AMPK.**
**a**, LCA elevates mitochondrial gene expression in an AMPK-dependent manner in aged mice. Mice were treated as in Fig. 4a, followed by determining the mRNA levels of OXPHOS and TCA cycle genes in the muscle. Results are mean ± s.e.m.; $n = 6$, and $P$ value by two-way ANOVA followed by Tukey's test.
**b**, LCA elevates RQ in an AMPK-dependent manner in aged mice. Mice were treated as in Fig. 4e, followed by determining RQ and ambulatory activity. Data are shown as mean (leftmost panel; at 5-min intervals during a 24-h course), or as box-and-whisker plots (other panles); $n = 4$ mice for each treatment, and $P$ value by two-way ANOVA followed by Tukey's test.
Experiments in this figure were performed three times.

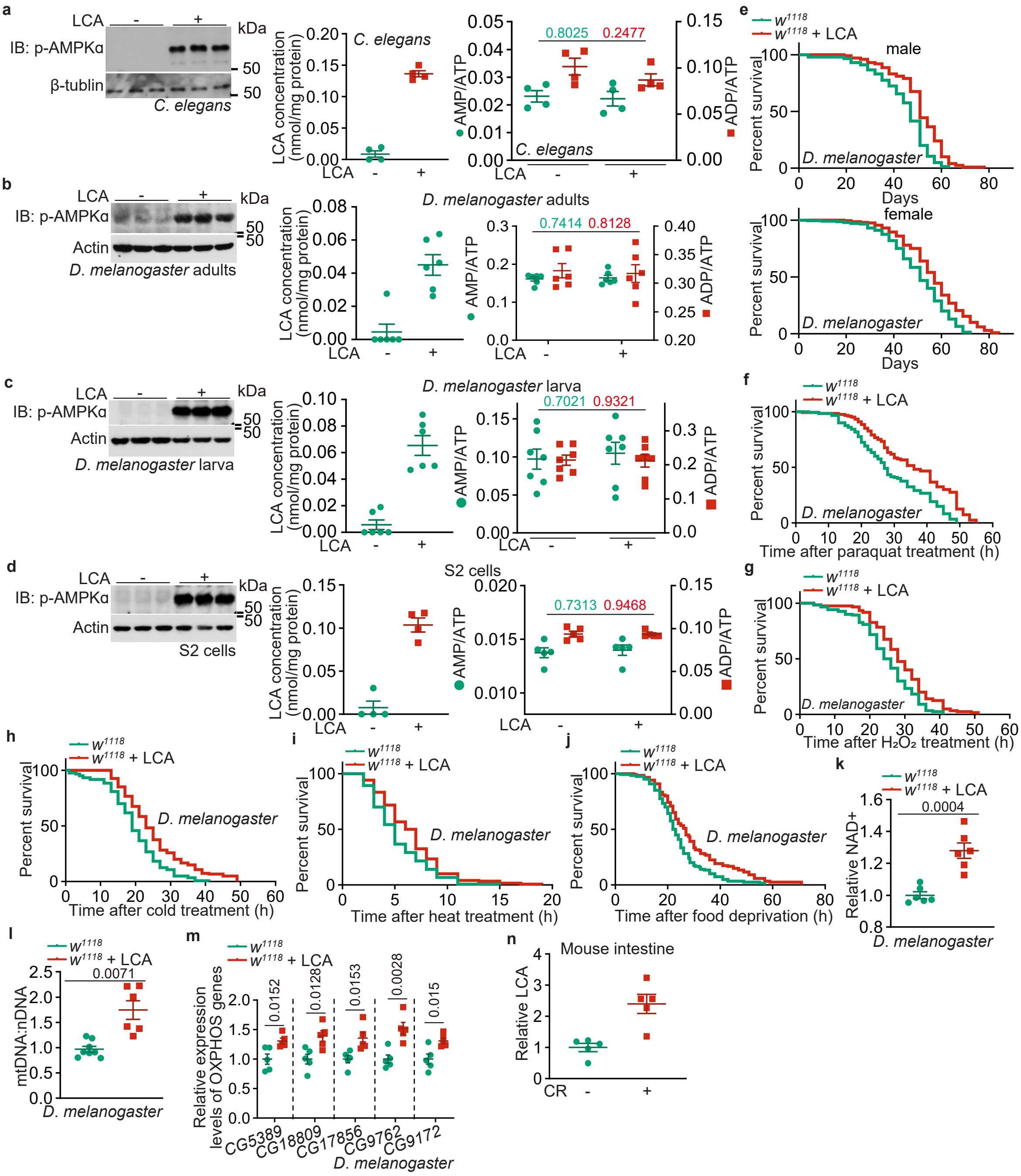


**Extended Data Fig. 5 | LCA activates AMPK in nematodes and flies in a similar way to that in mice.**

**a-d**, LCA, when absorbed into nematodes and flies to the same as seen in mouse muscles, activates AMPK without elevating AMP levels. Nematodes at L4 stage (**a**), adult flies (**b**, mixed gender), third instar larvae of flies (**c**) and the S2 cells (**d**) were cultured in agar medium containing 100 μM LCA (**a-c**) for 1 day (**a**, **c**) or 7 days (**b**), or in Schneider's *Drosophila* Medium containing 100 μM LCA for 2 h (**d**), followed by determining AMPK activation by immunoblotting (left panels of **a-d**), concentrations of LCA by HPLC-MS (middle panels of **a-d**) and the AMP:ATP and ADP:ATP ratios (right panels of **a-d**). Results are mean ± s.e.m.; *n* = 4 (**a**, and middle panel of **d**), 5 (right panel of **d**), 7 (right panel of **c**), or 6 (others) samples for each treatment, and *P* value by two-sided Student's *t*-test.

**e-m**, LCA improves the lifespan and healthspan of flies. Flies were treated with LCA as in Fig. 5b, followed by determination of lifespan (**e**), oxidative resistance (**f**, **g**), cold resistance (**h**), heat resistance (**i**), food deprivation resistance (**j**), $NAD^+$ levels (**k**), mtDNA:nDNA levels (**l**), and OXPHOS gene expression levels (**m**) as in Fig. 5b, e, f, g, h, i, j and k, respectively, except that the wildtype ($w^{1118}$) strain were used. Results of **k**, **l** and **m** are shown as mean ± s.e.m.; *n* = 5 (**m**), 8 (vehicle group of **l**) or 6 (others) samples for each treatment, and *P* value by two-sided Student's *t*-test.

**n**, CR elevates intestinal concentrations of LCA. Mice were subjected to CR for 4 months, followed by determining the concentrations of LCA in the intestine. Results are mean ± s.e.m., normalised to the AL group; *n* = 5 mice for each treatment.

Experiments in this figure were performed three times.

| Name | CAS number | Fold changes (CR vs AL) | *P* value | Dissolution | Stock concentration* | Concentration reported | Reference (PMID) | Concentration used in screening assay | Concentration ranges able to activate AMPK |
|---|---|---|---|---|---|---|---|---|---|
| (S)-3,4-Dihydroxybutyric acid | 51267-44-8 | 1.497 | < 0.0001 | $H_2O$ | 500 mM | | | 100 µM - 10 mM | |
| 1,3-Diaminopropane | 109-76-2 | 2.695 | 0.0394 | | 11.98 M | | | 100 µM - 10 mM | |
| 1,5-Anhydro-D-sorbitol | 154-58-5 | 0.8497 | 0.002 | $H_2O$ | 200 mM | | | 100 µM - 10 mM | |
| 10-Hydroxydecanoic acid | 1679-53-4 | 3.283 | < 0.0001 | DMSO | 200 mM | 5 mM | 23995057 | 100 µM - 20 mM | |
| 18α-Glycyrrhetinic acid | 1449-05-4 | 26.2 | < 0.0001 | DMSO, ultrasonic | 20 mM | 16 µM | 23362936 | 1 µM - 100 µM | |
| 18β-Glycyrrhetinic acid | 471-53-4 | 26.2 | < 0.0001 | DMSO | 200 mM | 320 µM | 24695790 | 100 µM - 10 mM | |
| 1-Aminocyclopentanecarboxylic acid | 52-52-8 | 0.6778 | 0.0068 | $H_2O$ | 100 mM | 10 mM | 37323630 | 100 µM - 10 mM | |
| 1-Methyladenosine | 15763-06-1 | 1.491 | 0.0349 | DMSO, ultrasonic | 500 mM | | | 100 µM - 10 mM | 1 mM |
| 1-Methylnicotinamide | 1005-24-9 | 0.5474 | 0.0011 | $H_2O$ | 100 mM | 1 mM | 33907844 | 100 µM - 10 mM | |
| 1-Monopalmitin | 542-44-9 | 0.6402 | < 0.0001 | DMSO | 100 mM | 300 µM | 15630255 | 100 µM - 10 mM | |
| 2',3'-cCMP | 15718-51-1 | 1.566 | 0.0181 | $H_2O$ | 100 mM | | | 100 µM - 10 mM | |
| 2,3-Dihydroxybenzoic acid | 303-38-8 | 0.295 | < 0.0001 | DMSO | 500 mM | | | 100 µM - 10 mM | |
| 2,3-Dihydroxybutanoic acid | 759-06-8 | 2.848 | < 0.0001 | DMSO | 10 mM | | | 10 µM - 1 mM | |
| 2,4-Diaminobutyric acid | 1758-80-1 | 1.94 | 0.0248 | $H_2O$, ultrasonic | 5 mM | 5 mM | 7430241 | 5 µM - 500 µM | |
| 2,4-Dihydroxybutanoic acid | 1518-62-3 | 0.8602 | 0.0033 | DMSO | 10 mM | | | 10 µM - 1 mM | |
| 2,4-Dihydroxypyrimidine-5-carboxylic acid | 23945-44-0 | 2.014 | 0.041 | DMSO | 100 mM | | | 100 µM - 10 mM | |
| 2,6-Diaminopimelic acid | 583-93-7 | 1.901 | 0.0071 | $H_2O$ | 20 mM | | | 10 µM - 1 mM | |
| 24-Nor UDCA | 99697-24-2 | 18.46 | 0.0019 | DMSO | 10 mM | | | 10 µM - 1 mM | |
| 2-Aminoadipic acid | 542-32-5 | 0.6892 | 0.0002 | $H_2O$ | 10 mM | | 37311782 | 10 µM - 1 mM | |
| 2'-CMP | 85-94-9 | 0.2884 | 0.0022 | | NA | | | | |
| 2'-Deoxycytidine | 951-77-9 | 0.7203 | 0.0045 | $H_2O$ | 100 mM | 30 µM | 36227698 | 10 µM - 1 mM | |
| 2-Deoxyribose 1-phosphate | 17210-42-3 | 5.159 | 0.0337 | | NA | | | | |
| 2-Deoxystreptamine | 84107-26-6 | 0.6636 | 0.0002 | | 4.4 M | | | 100 µM - 10 mM | |
| 2-Furoic acid | 88-14-2 | 1.81 | 0.0238 | DMSO | 500 mM | | | 100 µM - 10 mM | |
| 2-HG | 103404-90-6 | 1.891 | 0.0006 | $H_2O$ | 200 mM | | 21251613 | 100 µM - 10 mM | |
| 2-Hydroxy-2-methylbutanedioic acid | 597-44-4 | 1.683 | 0.0033 | | NA | | | | |
| 2-Hydroxy-3-methylbutyric acid | 4026-18-0 | 2.486 | < 0.0001 | DMSO, ultrasonic | 500 mM | | | 100 µM - 10 mM | |
| 2-Hydroxy-4-(methylthio)butanoate | 4857-44-7 | 0.2511 | < 0.0001 | $H_2O$ | 20 mM | | | 10 µM - 1 mM | |
| 2-Hydroxy-4-methylvaleric acid | 498-36-2 | 4.192 | < 0.0001 | | 8.7 M | 75.7 mM | 10.1016/j.joen.2019.01.012 | 870 µM - 8.7 mM | |
| 2-Hydroxybutyrate | 5094-24-6 | 2.45 | 0.0016 | $H_2O$ | 200 mM | | | 100 µM - 10 mM | |
| 2-Hydroxybutyric acid | 600-15-7 | 4.063 | < 0.0001 | DMSO | 500 mM | 32 mM | 16922922 | 100 µM - 40 mM | |
| 2-Keto-3-methylbutanoic acid | | 1.608 | < 0.0001 | | NA | | | | |
| 2-Keto-3-methylvaleric acid | 488-15-3 | 2.156 | < 0.0001 | DMSO | 10 mM | | | 10 µM - 1 mM | |
| 2-Methoxybenzoic acid | 579-75-9 | 1.78 | 0.0002 | Ethanol | 500 mM | 2 mM | 2172923 | 100 µM - 10 mM | |
| 2-Methylglutarate | 617-62-9 | 1.356 | 0.0065 | DMSO, ultrasonic | 500 mM | | | 100 µM - 10 mM | |
| 2-Methylmaleate | 498-23-7 | 1.378 | 0.0007 | $H_2O$ | 500 mM | | | 100 µM - 10 mM | |
| 2-Methylserine | 16820-18-1 | 1.276 | 0.0191 | DMSO | 15.28 mM | | | 15.28 µM - 1.528 mM | |
| 2-oxo-4-Methylthiobutanoate | 51828-97-8 | 0.2614 | < 0.0001 | $H_2O$ | 500 mM | 10 mM | 23385593 | 100 µM - 50 mM | |
| 2-Oxobutyric acid | 600-18-0 | 2.885 | 0.0097 | DMSO | 200 mM | | | 100 µM - 10 mM | |
| 2-Oxoisovaleric acid | 759-05-7 | 1.761 | 0.0007 | DMSO | 500 mM | | | 100 µM - 10 mM | |
| 2-quinolinecarboxylate | 93-10-7 | 3.107 | 0.0033 | DMSO | 500 mM | | | 100 µM - 10 mM | |
| 2-Thiopheneacetic acid | 1918-77-0 | 4.08 | 0.0097 | | 9.6 M | | | 100 µM - 10 mM | |
| 3-Aminophenol | 591-27-5 | 1.542 | 0.0483 | $H_2O$ | 300 mM | | | 100 µM - 10 mM | |
| 3'-AMP | 84-21-9 | 0.00568 | 0.0014 | $H_2O$ | 100 mM | 30 µM | 3083111 | 10 µM - 1 mM | |
| 3-AP | 143621-35-6 | 0.7758 | 0.0036 | DMSO | 200 mM | 1 µM | 37097390 | 100 nM - 10 µM | |
| 3-Guanidinopropionic acid | 353-09-3 | 0.3029 | < 0.0001 | $H_2O$ | 200 mM | 10 mM | 34613776 | 100 µM - 20 mM | |
| 3-Hydroxyanthranilate | 548-93-6 | 17.78 | 0.0149 | DMSO | 300 mM | | | 100 µM - 20 mM | |
| 3-Hydroxybutyric acid | 300-85-6 | 12.07 | < 0.0001 | $H_2O$ | 200 mM | 10 mM | 25686106 | 100 µM - 20 mM | |
| 3-Hydroxyisovaleric acid | 625-08-1 | 3.686 | < 0.0001 | DMSO | 200 mM | | | 100 µM - 10 mM | |
| 3-Hydroxymethylglutarate | 503-49-1 | 1.656 | 0.0126 | DMSO | 1 M | | | 100 µM - 10 mM | |
| 3-Indolelactic acid | 1821-52-9 | 1.321 | 0.0004 | DMSO | 10 mM | 5 µM | 33850185 | 1 µM - 100 µM | |
| 3-Keto-7α,12α-dihydroxy-5α-CA | 14772-92-0 | 68.92 | 0.0114 | | NA | | | | |
| 3-Methyl-2-ketobutyric acid | | 1.882 | < 0.0001 | | NA | | | | |
| 3-Methyl-2-oxovaleric acid | 1460-34-0 | 2.267 | 0.0002 | $H_2O$ | 500 mM | 20 µM | 33772024 | 10 µM - 1 mM | |
| 3-Methyladenine | 5142-23-4 | 2.519 | 0.0097 | $H_2O$ | 10 mM | 10 mM | 22545128 | 10 µM - 1 mM | |
| 3-Methylhistidine | 368-16-1 | 0.6847 | < 0.0001 | $H_2O$ | 500 mM | | | 100 µM - 10 mM | |
| 3-S-Methylthiopropionate | 13532-18-8 | 0.3758 | 0.0068 | | 8 M | | | 100 µM - 10 mM | |
| 3β-UDCA | 78919-26-3 | 56.41 | 0.0069 | DMSO | 10 mM | | | 10 µM - 1 mM | |
| 4-Amino-3-hydroxybutyric acid | 924-49-2 | 0.3422 | 0.0041 | $H_2O$ | 100 mM | | | 100 µM - 10 mM | |
| 4-Aminosalicylic acid | 65-49-6 | 0.1185 | 0.0337 | DMSO | 500 mM | | | 100 µM - 10 mM | |
| 4-guanidinobutanoate | 463-00-3 | 0.5057 | < 0.0001 | $H_2O$ | 100 mM | | | 100 µM - 10 mM | |
| 4-Methyl-2-oxovaleric acid | 816-66-0 | 2.267 | 0.0002 | $H_2O$ | 500 mM | 200 µM | 35614056 | 100 µM - 10 mM | 1 mM - 10 mM |
| 4-Pyridoxic acid | 82-82-6 | 1.701 | 0.0118 | DMSO | 20 mM | 10 µM | 33840062 | 1 µM - 100 µM | |
| 5,8,11-Eicosatrienoic acid | 20590-32-3 | 0.5564 | 0.0002 | Ethanol | 32.63 mM | | | 32.63 µM - 3.263 mM | |
| 5-Amino-4-oxovaleric acid | 106-60-5 | 0.7426 | 0.0001 | DMSO, ultrasonic | 500 mM | 20 µM | 10.1038/s41419-021-04277-4 | 10 µM - 1 mM | |
| 5-Aminovaleric acid | 660-88-8 | 0.5865 | 0.0346 | $H_2O$ | 500 mM | | | 100 µM - 10 mM | |
| 5-Hydroxylysine | 13204-98-3 | 0.2964 | 0.0005 | $H_2O$ | 500 mM | | | 100 µM - 10 mM | |
| 5-Methyl-2'-deoxycytidine | 838-07-3 | 0.7017 | 0.0125 | $H_2O$ | 400 mM | | | 100 µM - 10 mM | |
| 5-Methylcytosine | 554-01-8 | 1.544 | 0.0386 | $H_2O$, ultrasonic | 50 mM | | | 10 µM - 1 mM | |
| 5-Oxohexanoic acid | 3128-06-1 | 0.4241 | 0.0007 | DMSO | 500 mM | | | 100 µM - 10 mM | |
| 6,8-Thioctic acid | 1077-28-7 | 3.036 | 0.0018 | DMSO | 200 mM | 1 mM | 25123628 | 100 µM - 10 mM | |
| 6-Aminohexanoic acid | 60-32-2 | 2.577 | 0.0283 | $H_2O$ | 200 mM | | | 100 µM - 10 mM | |
| 6-Hydroxynicotinic acid | 5006-66-6 | 5.183 | 0.0337 | DMSO | 200 mM | | | 100 µM - 10 mM | |

**Extended Data Table 1**

| Name | CAS number | Fold changes (CR vs AL) | *P* value | Dissolution | Stock concent ration* | Concent ration reported | Reference (PMID) | Concentration used in screening assay | Concentration ranges able to activate AMPK |
|---|---|---|---|---|---|---|---|---|---|
| 7,8-Dihydrobiopterin | 6779-87-9 | 0.6307 | 0.0016 | $H_2O$ | 10 mM | | | 10 µM - 1 mM | |
| Acetoacetic acid | 3483-11-2 | 3.668 | 0.0059 | $H_2O$ | 100 mM | 10 mM | 31242642 | 100 µM - 10 mM | |
| Acetylcholine | 60-31-1 | 0.229 | 0.0003 | $H_2O$ | 500 mM | 50 µM | 18386025 | 100 µM - 50 mM | |
| Acetylphosphate | 94249-01-1 | 1.292 | 0.0037 | $H_2O$ | 200 mM | | | 100 µM - 10 mM | |
| Aconitate | 585-84-2 | 1.348 | 0.0082 | $H_2O$ | 500 mM | | | 100 µM - 10 mM | |
| Adenine | 73-24-5 | 0.08213 | 0.002 | DMSO | 50 mM | | | 10 µM - 1 mM | |
| Adenosine | 58-61-7 | 0.08209 | < 0.0001 | $H_2O$ | 20 mM | 10 µM | 32205841 | 1 µM - 100 µM | |
| ADP-ribose | 68414-18-6 | 1.818 | 0.0337 | $H_2O$ | 10 mM | | | 10 µM - 1 mM | |
| Agmatine | 2482-00-0 | 2.201 | 0.001 | $H_2O$ | 200 mM | | 7906055 | 100 µM - 10 mM | |
| Alanylalanine | 1948-31-8 | 1.929 | 0.0335 | DMSO | 1 M | | | 100 µM - 10 mM | |
| Allantoic acid | 99-16-1 | 1.988 | < 0.0001 | DMSO | 10 mM | | | 10 µM - 1 mM | |
| Allantoin | 97-59-6 | 1.376 | 0.0256 | DMSO | 200 mM | 100 µM | 26290782 | 10 µM - 1 mM | |
| Allocholic acid | 2464-18-8 | 180.2 | 0.0011 | DMSO | 50 mM | 50 µM | 19464381 | 10 µM - 1 mM | |
| Aminomalonic acid | 1068-84-4 | 1.329 | 0.0241 | $H_2O$, ultrasonic | 50 mM | | | 10 µM - 1 mM | |
| AMP | 61-19-8 | 0.008405 | 0.0003 | DMSO | 20 mM | 10 µM | | 1 µM - 100 µM | |
| Anserine | 584-85-0 | 0.4378 | < 0.0001 | $H_2O$, ultrasonic | 200 mM | | | 100 µM - 10 mM | |
| Anthranilate | 134-20-3 | 0.4903 | 0.0008 | | 7.7 M | | | 100 µM - 10 mM | |
| Anthranilic acid | 118-92-3 | 0.3389 | < 0.0001 | | 32.8 mM | | | 32.8 µM - 3.28 mM | |
| Arachidonic acid | 506-32-1 | 0.7109 | 0.0003 | DMSO | 10 mM | 40 µM | 37611494 | 10 µM - 1 mM | |
| Arginine | 74-79-3 | 0.697 | 0.0009 | $H_2O$ | 200 mM | 40 mM | 34206713 | 100 µM - 20 mM | |
| Arginine ethyl ester | 36589-29-4 | 0.787 | 0.0337 | $H_2O$ | 10 mM | | | 10 µM - 1 mM | |
| Asparagine | 70-47-3 | 0.7069 | < 0.0001 | $H_2O$ | 50 mM | 1 mM | 34704268 | 100 µM - 5 mM | |
| ATP | 34369-07-8 | 0.5874 | 0.038 | $H_2O$ | 100 mM | | 31394382 | 100 µM - 10 mM | |
| Atropine | 51-55-8 | 2.417 | 0.035 | DMSO | 200 mM | 100 µM | 29860464 | 10 µM - 1 mM | |
| Benzoate | 532-32-1 | 0.6589 | 0.0431 | DMSO | 20 mM | | | 10 µM - 1 mM | |
| Benzoic acid | 65-85-0 | 1.761 | < 0.0001 | $H_2O$ | 50 mM | 25 mM | 8140053 | 100 µM - 5 mM | |
| Betaine | 107-43-7 | 1.425 | 0.0013 | $H_2O$ | 500 mM | 10 mM | 23246691 | 100 µM - 50 mM | |
| Biotin | 58-85-5 | 1.831 | 0.0016 | DMSO | 200 mM | 1 µM | 31039616 | 100 nM - 10 µM | |
| Bis(p-nitrophenyl)phosphate | 645-15-8 | 1.517 | 0.0421 | | 4.7 M | | | 100 µM - 10 mM | |
| Butyric acid | 107-92-6 | 4.946 | < 0.0001 | | 10.94 M | 10 mM | 28356979 | 109.4 µM - 10.94 mM | |
| CA | 81-25-4 | 40.34 | 0.0041 | DMSO | 100 mM | 2.45 mM | 10.2147/IJN.S125047 | 100 µM - 10 mM | |
| Cadaverine | 462-94-2 | 3.038 | 0.0015 | | 8.5 M | | | 100 µM - 10 mM | |
| Campesterol | 474-62-4 | 0.8264 | 0.0152 | $H_2O$ | 1 mM | 49.9 µM | 17604370 | 1 µM - 100 µM | |
| Canavanine | 2219-31-0 | 4.008 | 0.0083 | $H_2O$ | 200 mM | 1 mM | 29658590 | 100 µM - 10 mM | |
| Carnitine | 541-15-1 | 0.6658 | < 0.0001 | $H_2O$ | 200 mM | 0.5 mM | 31155494 | 100 µM - 10 mM | |
| Carnosine | 305-84-0 | 0.8751 | 0.0337 | $H_2O$ | 100 mM | | | 100 µM - 10 mM | |
| cCMP | | 1.709 | 0.0181 | | NA | | | | |
| CDCA | 474-25-9 | 42.15 | 0.0074 | DMSO | 100 mM | 50 µM | 33338411 | 10 µM - 1 mM | |
| cGMP | 40732-48-7 | 1.661 | 0.0083 | $H_2O$ | 10 mM | 10 µM | 12933699 | 1 µM - 100 µM | |
| Cholesterol | 57-88-5 | 0.7934 | < 0.0001 | Ethanol, ultrasonic | 20 mM | 30 µM | 34354264 | 10 µM - 1 mM | |
| Choline | 62-49-7 | 1.221 | 0.0004 | $H_2O$ | 100 mM | 130 µM | 9263575 | 10 µM - 1 mM | |
| cis-11-Eicosenoic acid | 5561-99-9 | 0.7225 | 0.0268 | DMSO | 200 mM | 100 µM | 26918025 | 10 µM - 1 mM | |
| Citramalic acid | 6236-10-8 | 3.377 | < 0.0001 | $H_2O$ | 100 mM | | | 10 µM - 1 mM | |
| Citric acid | 77-92-9 | 1.351 | < 0.0001 | $H_2O$ | 500 mM | 12.5 mM | 24123010 | 100 µM - 50 mM | |
| Citrulline | 372-75-8 | 1.497 | 0.0098 | $H_2O$ | 200 mM | 1 mM | 24400007 | 100 µM - 10 mM | |
| CMP | 6757-06-8 | 0.4511 | < 0.0001 | $H_2O$ | 250 mM | | | 100 µM - 20 mM | |
| CMP-N-acetylneuraminate | 3063-71-6 | 0.6356 | 0.0281 | $H_2O$ | 50 mM | | | 10 µM - 1 mM | |
| Creatine | 57-00-1 | 0.6627 | < 0.0001 | $H_2O$ | 10 mM | 1 mM | 35276125 | 10 µM - 1 mM | |
| Creatinine | 60-27-5 | 1.773 | 0.0002 | $H_2O$, ultrasonic | 100 mM | | | 100 µM - 10 mM | |
| Crotonic acid | 107-93-7 | 1.817 | 0.0089 | $H_2O$ | 200 mM | | | 100 µM - 10 mM | |
| Cystathionine | 56-88-2 | 0.3881 | < 0.0001 | $H_2O$ | 100 mM | 1 mM | 31885769 | 100 µM - 10 mM | |
| Cysteine | 52-90-4 | 6.537 | < 0.0001 | $H_2O$, ultrasonic | 200 mM | 200 µM | 37781038 | 10 µM - 1 mM | |
| Cystine | 56-89-3 | 0.6813 | < 0.0001 | 0.1 M HCl, ultrasonic | 10 mM | | | 10 µM - 1 mM | |
| Cytidine | 65-46-3 | 1.344 | 0.0058 | $H_2O$ | 200 mM | 50 µM | 35022212 | 10 µM - 1 mM | |
| Cytosine | 71-30-7 | 2.299 | 0.0141 | DMSO | 100 mM | | | 100 µM - 10 mM | |
| D-(+)-Talose | 2595-98-4 | 0.4597 | < 0.0001 | $H_2O$ | 500 mM | | | 100 µM - 10 mM | |
| D-2-Aminobutyric acid | 2623-91-8 | 1.564 | 0.0005 | $H_2O$ | 500 mM | 1 mM | 37445626 | 100 µM - 10 mM | |
| dAMP | 653-63-4 | 5.548 | < 0.0001 | DMSO | 100 mM | 100 µM | 31634358 | 10 µM - 1 mM | |
| DCA | 83-44-3 | 29.2 | 0.0026 | DMSO | 200 mM | 50 µM | 33338411 | 10 µM - 1 mM | |
| dCDP | 151151-32-5 | 10.98 | 0.0023 | $H_2O$ | 200 mM | | | 100 µM - 10 mM | |
| dCMP | 1032-65-1 | 2.621 | 0.0222 | $H_2O$ | 20 mM | | | 10 µM - 1 mM | |
| Decanoic acid | 334-48-5 | 1.405 | < 0.0001 | DMSO | 500 mM | 1 mM | 30324610 | 100 µM - 10 mM | |
| D-Glucose | 50-99-7 | 0.4664 | < 0.0001 | $H_2O$ | 200 mM | 10 mM | 34486387 | 100 µM - 20 mM | |
| dGMP | 33430-61-4 | 0.008708 | 0.0004 | $H_2O$ | 200 mM | 1.2 mM | 19221117 | 100 µM - 10 mM | |
| DHA | 6217-54-5 | 0.7943 | 0.0169 | DMSO | 100 mM | 10 µM | 35177641 | 1 µM - 100 µM | |
| Dihydroorotate | 5988-19-2 | 2.293 | < 0.0001 | $H_2O$ | 20 mM | | | 10 µM - 1 mM | |
| DL-Leucine | 328-39-2 | 1.984 | 0.0097 | $H_2O$, ultrasonic | 50 mM | | | 10 µM - 1 mM | |
| DL-Ornithine | 70-26-8 | 0.7869 | 0.0288 | $H_2O$ | 200 mM | 10 mM | 31770578 | 100 µM - 20 mM | |
| D-Mannitol | 69-65-8 | 0.5612 | 0.0004 | $H_2O$ | 100 mM | 40 mM | 37258572 | 100 µM - 10 mM | |
| Dodecanoic acid | 143-07-7 | 1.488 | < 0.0001 | DMSO, ultrasonic | 200 mM | 1 mM | 24356281 | 100 µM - 10 mM | |
| D-Tryptophan | 153-94-6 | 1.256 | 0.0034 | DMSO, ultrasonic | 10 mM | | | 10 µM - 1 mM | |
| Eicosanoic acid | 506-30-9 | 0.6611 | < 0.0001 | DMSO, ultrasonic | 10 mM | 100 µM | 36997100 | 10 µM - 1 mM | |
| Erythronic acid | 13752-84-6 | 1.239 | < 0.0001 | $H_2O$ | 500 mM | | | 100 µM - 10 mM | |
| Ethyl 2, 5-dihydroxybenzoate | 3943-91-7 | 0.2895 | < 0.0001 | DMSO | 100 mM | | | 10 µM - 1 mM | |
| Ethylmalonate | 601-75-2 | 1.856 | 0.0008 | DMSO | 1 M | | | 100 µM - 10 mM | |

**Extended Data Table 1 (cont.)**

| Name | CAS number | Fold changes (CR vs AL) | *P* value | Dissolution | Stock concent ration* | Concent ration reported | Reference (PMID) | Concentration used in screening assay | Concentration ranges able to activate AMPK |
|---|---|---|---|---|---|---|---|---|---|
| FAD | 84366-81-4 | 2.159 | 0.0259 | $H_2O$ | 10 mM | | | 10 μM - 1 mM | |
| Ferulic acid | 537-98-4 | 2.209 | 0.0005 | DMSO | 200 mM | 0.6 mM | 27733866 | 10 μM - 1 mM | 200 μM |
| Fructose | 7660-25-5 | 0.8553 | 0.0023 | $H_2O$ | 500 mM | 2.5 mM | 26104185 | 100 μM - 10 mM | |
| Fumaric acid | 110-17-8 | 2.159 | < 0.0001 | DMSO | 200 mM | 400 μM | 27580029 | 100 μM - 10 mM | |
| Galactosamine | 1772-03-8 | 0.2442 | 0.0005 | $H_2O$ | 100 mM | | | 100 μM - 10 mM | |
| Galacturonic acid | 91510-62-2 | 2.269 | 0.0045 | $H_2O$ | 500 mM | | | 100 μM - 10 mM | |
| GCA | 475-31-0 | 1.773 | 0.0293 | DMSO | 100 mM | 250 μM | 18606222 | 100 μM - 10 mM | |
| GDCA | 360-65-6 | 3.952 | 0.012 | DMSO | 200 mM | | | 100 μM - 10 mM | |
| Gluconic acid | 526-95-4 | 0.6811 | 0.0009 | $H_2O$ | 5.098 M | | | 509.8 μM - 50.9 mM | |
| Glucono-D-lactone | 90-80-2 | 0.1418 | < 0.0001 | $H_2O$ | 500 mM | | | 100 μM - 10 mM | |
| Glucosamine | 3416-24-8 | 0.272 | 0.0023 | $H_2O$, ultrasonic | 100 mM | 40 mM | 24959958 | 100 μM - 10 mM | |
| Glucosamine 6-phosphate | 3616-42-0 | 1.77 | 0.0287 | $H_2O$ | 100 mM | | | 100 μM - 10 mM | |
| Glucuronic acid | 6556-12-3 | 2.289 | 0.0028 | $H_2O$ | 500 mM | | | 100 μM - 10 mM | |
| Glutamate | 6106-04-3 | 1.095 | 0.0333 | $H_2O$ | 500 mM | | | 100 μM - 10 mM | |
| Glutaric acid | 110-94-1 | 8.302 | < 0.0001 | $H_2O$ | 500 mM | 5 mM | 31104295 | 100 μM - 50 mM | |
| Glutathione (GSSG) | 27025-41-8 | 0.815 | 0.0131 | $H_2O$ | 200 mM | | | 100 μM - 10 mM | |
| Glyceric acid | 473-81-4 | 1.313 | 0.0018 | DMSO | 500 mM | 7.8 mM | 25229854 | 100 μM - 20 mM | |
| Glycerol-3-phosphate | 17989-41-2 | 0.7651 | 0.0031 | $H_2O$ | 500 mM | 10 μM | 32065590 | 1 μM - 100 μM | |
| Glycolic acid | 79-14-1 | 1.265 | 0.0002 | $H_2O$ | 1 M | 6.57 mM | 14756523 | 100 μM - 20 mM | |
| Glyoxylic acid | 298-12-4 | 1.311 | 0.0205 | DMSO | 1 M | 0.8 mM | 37049603 | 100 μM - 10 mM | |
| GMP | | 0.02821 | < 0.0001 | | NA | | | | |
| Guanidinoacetate | 352-97-6 | 6.971 | < 0.0001 | HCL | 200 mM | | | 100 μM - 10 mM | |
| Guanidinosuccinate | 6133-30-8 | 1.741 | 0.0003 | DMSO, ultrasonic | 400 mM | | | 100 μM - 10 mM | |
| HDCA | 83-49-8 | 14.89 | 0.0026 | DMSO | 100 mM | 50 μM | 33338411 | 10 μM - 1 mM | |
| Heptadecanoic acid | 506-12-7 | 1.22 | 0.0079 | DMSO | 100 mM | 250 μM | 31002344 | 100 μM - 10 mM | |
| Hexanoic acid | 142-62-1 | 1.289 | 0.0241 | DMSO, ultrasonic | 500 mM | 7 mM | 32708494 | 100 μM - 20 mM | |
| Hippurate | 532-94-5 | 0.3672 | < 0.0001 | $H_2O$ | 10 mM | | | 10 μM - 1 mM | |
| Histamine | 51-45-6 | 2.195 | 0.0003 | $H_2O$ | 200 mM | 50 μM | 36113478 | 10 μM - 1 mM | |
| Histidine | 71-00-1 | 0.8569 | 0.0463 | $H_2O$ | 100 mM | | | 100 μM - 10 mM | |
| Homocarnosine TFA | 3650-73-5 | 0.5318 | < 0.0001 | $H_2O$ | 20 mM | | | 10 μM - 1 mM | |
| Homoserine | 672-15-1 | 0.6646 | < 0.0001 | $H_2O$ | 1 M | | | 100 μM - 10 mM | |
| HyoCA | 547-75-1 | 177.7 | 0.0075 | DMSO | 100 mM | 50 μM | 33338411 | 10 μM - 1 mM | |
| Hypotaurine | 300-84-5 | 0.8595 | 0.0245 | $H_2O$ | 200 mM | 10 mM | 33937232 | 100 μM - 20 mM | |
| Hypoxanthine | 68-94-0 | 14.32 | 0.0111 | DMSO | 50 mM | 100 μM | 34703880 | 10 μM - 1 mM | |
| Ibuprofen | 15687-27-1 | 1.514 | 0.025 | DMSO | 200 mM | 500 μM | 25542229 | 100 μM - 10 mM | |
| Ibuprofen | 288-13-1 | 2.849 | 0.0337 | $H_2O$ | 500 mM | | | 100 μM - 10 mM | |
| Imidazole-4-acetic acid | 645-65-8 | 0.7248 | 0.0281 | DMSO | 50 mM | | | 10 μM - 1 mM | |
| IMP(Inosinic acid) | 131-99-7 | 0.1539 | 0.0004 | $H_2O$, ultrasonic | 200 mM | 15 mM | 36224248 | 100 μM - 20 mM | |
| Indole-3-acetic acid | 87-51-4 | 6.745 | < 0.0001 | DMSO | 100 mM | 0.5 mM | 37155876 | 100 μM - 10 mM | |
| Inosine | 58-63-9 | 12.72 | 0.0074 | $H_2O$, ultrasonic | 10 mM | 300 μM | 26903141 | 10 μM - 1 mM | |
| Isoamylamine | 107-85-7 | 2.22 | 0.0024 | | 8.6 M | | | 100 μM - 10 mM | |
| Isobutyric acid | 79-31-2 | 6.5 | < 0.0001 | | 10.78 M | | | 10.78 μM - 1.078 mM | |
| Isocitric acid | 1637-73-6 | 7.866 | 0.0023 | $H_2O$ | 10 mM | 1 mM | 37582794 | 10 μM - 1 mM | |
| Isonicotinamide | 1453-82-3 | 0.4582 | < 0.0001 | $H_2O$ | 500 mM | | | 100 μM - 10 mM | |
| Isopropanolamine | 78-96-6 | 2.893 | < 0.0001 | DMSO | 1 M | | | 100 μM - 10 mM | |
| Kynurenine | 343-65-7 | 1.879 | < 0.0001 | 1 M HCl, ultrasonic | 50 mM | 100 μM | 27020609 | 10 μM - 1 mM | |
| L-5-Oxoproline | 98-79-3 | 1.189 | 0.0004 | $H_2O$ | 500 mM | | | 100 μM - 10 mM | |
| L-Alanine | 56-41-7 | 0.7131 | 0.0154 | $H_2O$ | 1.2 M | 1200 mM | 30127448 | 1.2 mM - 120 mM | |
| L-Aspartic acid | 56-84-8 | 0.4984 | < 0.0001 | $H_2O$ | 10 mM | 200 μM | 26325019 | 10 μM - 1 mM | |
| LCA | 434-13-9 | 22.65 | 0.0095 | DMSO | 100 mM | 50 μM | 33338411 | 1 μM - 500 μM | 1 μM - 100 μM |
| L-Glutamic acid | 56-86-0 | 1.235 | 0.0047 | $H_2O$ | 100 mM | 100 μM | 35363928 | 10 μM - 1 mM | |
| L-Hydroxyproline | 51-35-4 | 0.4206 | < 0.0001 | $H_2O$ | 200 mM | 10 mM | 18287100 | 100 μM - 20 mM | |
| Linoelaidic acid | 506-21-8 | 1.23 | 0.0241 | DMSO | 100 mM | 60 μM | 27255637 | 10 μM - 1 mM | |
| Lipoamide | 940-69-2 | 0.646 | 0.0023 | DMSO | 100 mM | 200 μM | 36347997 | 10 μM - 1 mM | |
| L-Isoleucine | 73-32-5 | 1.272 | 0.0331 | $H_2O$ | 100 mM | 1.5 mM | 36111572 | 100 μM - 10 mM | |
| L-Methionine | 63-68-3 | 0.4357 | < 0.0001 | $H_2O$ | 100 mM | 120 μM | 36429035 | 100 μM - 10 mM | |
| L-Talose | 23567-25-1 | 0.4597 | < 0.0001 | $H_2O$ | 500 mM | | | 100 μM - 10 mM | |
| L-Threonic acid | 70753-61-6 | 0.8582 | 0.0097 | $H_2O$ | 10 mM | | | 10 μM - 1 mM | |
| L-Threonine | 72-19-5 | 0.8097 | 0.0048 | $H_2O$ | 200 mM | | | 100 μM - 10 mM | |
| L-Tyrosine | 60-18-4 | 1.141 | 0.0481 | DMSO | 5 mM | 776 μM | 9057892 | 5 μM - 500 μM | |
| Lysine | 56-87-1 | 0.4772 | < 0.0001 | $H_2O$ | 500 mM | 1 mM | 35923149 | 100 μM - 10 mM | |
| Maleic acid | 110-16-7 | 5.74 | 0.0337 | $H_2O$ | 500 mM | 100 μM | 28392987 | 10 μM - 1 mM | |
| Malic acid | 6915-15-7 | 2.202 | < 0.0001 | $H_2O$ | 500 mM | 100 μM | 37632443 | 10 μM - 1 mM | |
| Malonate | 141-82-2 | 1.491 | 0.0209 | $H_2O$ | 100 mM | 10 μM | 34234892 | 1 μM - 100 μM | |
| Mandelate | 90-64-2 | 0.3589 | 0.0008 | DMSO | 500 mM | | | 100 μM - 10 mM | |
| Mandelic acid | 611-71-2 | 32.33 | 0.0011 | DMSO | 500 mM | | | 100 μM - 10 mM | 200 μM - 2 mM |
| Methyl 2, 5-dihydroxybenzoate | 2150-46-1 | 0.2895 | < 0.0001 | DMSO | 100 mM | | | 10 μM - 1 mM | |
| Methylmalonic acid | 516-05-2 | 3.113 | < 0.0001 | $H_2O$ | 500 mM | 5 mM | 32814897 | 10 μM - 5 mM | 100 μM - 500 μM |
| Mevalonate | 2618458-93-6 | 1.846 | 0.0083 | $H_2O$ | 500 mM | 110 μM | 34256063 | 10 μM - 1 mM | |
| Monostearin | 123-94-4 | 0.6863 | 0.0003 | DMSO | 100 mM | | | 100 μM - 10 mM | |
| Myo-Inositol | 87-89-8 | 0.7972 | < 0.0001 | $H_2O$ | 200 mM | 500 μM | 30604967 | 100 μM - 10 mM | |
| Myristic acid | 544-63-8 | 1.881 | < 0.0001 | DMSO | 1 M | 200 μM | 36750575 | 10 μM - 1 mM | |
| N,N-Dimethylglycine | 1118-68-9 | 1.388 | 0.0018 | $H_2O$ | 500 mM | | | 100 μM - 10 mM | |
| N,N-Dimethylhistidine | 24940-57-6 | 0.5077 | 0.0012 | DMSO | 100 mM | | | 10 μM - 1 mM | |
| N5-Ethylglutamine | 3081-61-6 | 3.152 | < 0.0001 | $H_2O$ | 500 mM | 2 mM | 26164708 | 100 μM - 10 mM | |

**Extended Data Table 1 (cont.)**

| Name | CAS number | Fold changes (CR vs AL) | *P* value | Dissolution | Stock concentration* | Concentration reported | Reference (PMID) | Concentration used in screening assay | Concentration ranges able to activate AMPK |
|---|---|---|---|---|---|---|---|---|---|
| N6,N6,N6-Trimethyllysine | 19253-88-4 | 0.5916 | < 0.0001 | | NA | | | | |
| N6-Acetyllysine | 692-04-6 | 0.6147 | < 0.0001 | $H_2O$ | 200 mM | | | 100 μM - 10 mM | |
| N-acetylasparagine | 4033-40-3 | 0.7137 | 0.0369 | DMSO | 500 mM | | | 100 μM - 10 mM | |
| N-Acetylaspartate | 997-55-7 | 1.398 | 0.0153 | $H_2O$ | 200 mM | 1.5 mM | 17943458 | 100 μM - 10 mM | |
| N-Acetylcysteine | 616-91-1 | 1.082 | 0.0035 | $H_2O$, ultrasonic | 500 mM | | 9592085 | 100 μM - 10 mM | |
| N-Acetylglucosamine | 7512-17-6 | 0.3845 | 0.0008 | $H_2O$ | 400 mM | 20 μM | 37349834 | 10 μM - 1 mM | |
| N-Acetylglucosamine 1-phosphate | 31281-59-1 | 0.6444 | 0.0219 | $H_2O$ | 10 mM | | | 10 μM - 1 mM | |
| N-acetylglycine | 543-24-8 | 6.751 | < 0.0001 | $H_2O$ | 400 mM | | | 100 μM - 10 mM | |
| N-Acetylmethionine | 65-82-7 | 1.478 | 0.0357 | Methanol | 200 mM | 1 mM | 37871522 | 100 μM - 10 mM | |
| N-Acetylneuraminic acid | 131-48-6 | 0.7665 | 0.0174 | $H_2O$ | 100 mM | | 37270929 | 100 μM - 10 mM | |
| N-Acetylphenylalanine | 2018-61-3 | 2.894 | < 0.0001 | $H_2O$ | 20 mM | | | 10 μM - 1 mM | |
| N-acetylproline | 68-95-1 | 0.2256 | 0.0174 | $H_2O$ | 100 mM | | | 100 μM - 10 mM | |
| $NADP^+$ | 24292-60-2 | 2.414 | 0.0004 | $H_2O$ | 10 mM | 1 mM | 36913068 | 10 μM - 1 mM | |
| Nalidixic acid | 389-08-2 | 1.049 | < 0.0001 | $H_2O$ | 20 mM | 10 μM | | 1 μM - 100 μM | |
| N-Carbamoylaspartic acid | 923-37-5 | 2.532 | 0.0068 | DMSO | 200 mM | | | 100 μM - 10 mM | |
| N-Carbamoyl-L-aspartate | 13184-27-5 | 0.6021 | 0.0218 | | NA | | | | |
| N-Formylmethionine | 4289-98-9 | 2.394 | < 0.0001 | DMSO | 500 mM | | | 100 μM - 10 mM | |
| Nicotinamide | 98-92-0 | 0.5809 | 0.0051 | $H_2O$ | 400 mM | 108 mM | 26660162 | 100 μM - 40 mM | |
| Nicotine | 54-11-5 | 2.127 | 0.0068 | | NA | | | | |
| N-Methylalanine | 600-21-5 | 13.11 | < 0.0001 | $H_2O$ | 500 mM | | | 100 μM - 10 mM | |
| N-Methylaniline | 100-61-8 | 1.954 | 0.0105 | | 9.2 M | | | 100 μM - 10 mM | |
| N-Methylaspartate | 6384-92-5 | 1.094 | 0.0023 | $H_2O$ | 200 mM | 25 μM | 10.1186/s12987-022-00364-6 | 10 μM - 1 mM | |
| N-Methylglutamic acid | 35989-16-3 | 2.142 | 0.0066 | $H_2O$ | 300 mM | | | 100 μM - 20 mM | |
| Nor CA | 60696-62-0 | 237.1 | 0.0284 | DMSO | 200 mM | | | 100 μM - 10 mM | |
| Noradrenaline | 108341-18-0 | 31.17 | 0.0097 | $H_2O$ | 100 mM | 50 μM | 35832088 | 10 μM - 1 mM | |
| Norspermidine | 56-18-8 | 2.592 | 0.0097 | $H_2O$ | 100 mM | | | 100 μM - 10 mM | |
| O-Acetylcarnitine | 5080-50-2 | 1.701 | 0.0029 | $H_2O$ | 200 mM | 10 mM | 35185289 | 100 μM - 20 mM | |
| Octopine | 34522-32-2 | 0.4644 | < 0.0001 | | NA | | | | |
| o-Hydroxybenzoic acid | 69-72-7 | 0.1617 | < 0.0001 | DMSO | 200 mM | 100 μM | 22357964 | 10 μM - 1 mM | |
| Oleic Acid | 112-80-1 | 2.532 | < 0.0001 | DMSO, ultrasonic | 10 mM | 80 μM | 33271455 | 10 μM - 1 mM | |
| Orotate | 65-86-1 | 1.864 | < 0.0001 | DMSO | 300 mM | | | 100 μM - 10 mM | |
| Oxalic acid | 144-62-7 | 0.8033 | 0.0239 | $H_2O$ | 20 mM | 1 mM | 34834048 | 10 μM - 1 mM | |
| Oxaloacetate | 328-42-7 | 0.4386 | 0.0144 | $H_2O$ | 500 mM | | | 100 μM - 10 mM | |
| Oxoadipate | 3184-35-8 | 0.8148 | 0.0288 | $H_2O$ | 1 M | | | 100 μM - 10 mM | |
| P1,P4-di(adenosine-5') tetraphosphate | 102783-36-8 | 0.6822 | 0.0452 | $H_2O$ | 10 mM | | | 10 μM - 1 mM | |
| Palmitelaidic acid | 10030-73-6 | 2.089 | 0.0002 | Ethano, ultrasonic | 200 mM | | | 100 μM - 10 mM | |
| Palmitic Acid | 57-10-3 | 1.37 | < 0.0001 | Ethanol, ultrasonic | 50 mM | 300 μM | 28463985 | 100 μM - 5 mM | |
| Palmitoylcarnitine | 6865-14-1 | 2.204 | 0.0058 | $H_2O$ | 100 mM | 100 μM | 27403764 | 10 μM - 1 mM | |
| p-Aminobenzoate | 150-13-0 | 18.61 | 0.0149 | DMSO | 500 mM | | | 100 μM - 10 mM | |
| p-Anisic acid | 100-09-4 | 33.55 | 0.0011 | DMSO, ultrasonic | 500 mM | | | 100 μM - 10 mM | |
| Pantothenic acid | 79-83-4 | 2.219 | < 0.0001 | DMSO, ultrasonic | 200 mM | | | 100 μM - 10 mM | |
| Pelargonic acid | 112-05-0 | 1.194 | 0.0162 | DMSO | 500 mM | 3 mM | 36959559 | 100 μM - 10 mM | |
| Phenoxyacetic acid | 122-59-8 | 35 | 0.0011 | DMSO | 100 mM | | 333166 | 100 μM - 10 mM | |
| Phenyl phosphate | 66778-08-3 | 1.558 | 0.001 | $H_2O$ | 100 mM | | | 100 μM - 10 mM | |
| Phenylalanine | 63-91-2 | 1.144 | 0.0327 | $H_2O$ | 10 mM | 10 mM | 34953208 | 10 μM - 1 mM | |
| Phenyllactic acid | 828-01-3 | 2.192 | 0.0003 | DMSO | 500 mM | | | 100 μM - 10 mM | |
| Phenylpyruvate | 114-76-1 | 0.6609 | 0.0013 | $H_2O$ | 100 mM | | | 100 μM - 10 mM | |
| Phosphoenolpyruvic acid | 4265-07-0 | 11.34 | < 0.0001 | $H_2O$ | 100 mM | | | 100 μM - 10 mM | |
| Phosphorylcholine | 3616-04-4 | 0.419 | 0.0097 | $H_2O$ | 500 mM | | | 100 μM - 10 mM | |
| Phosphorylethanolamine | 1071-23-4 | 0.6001 | < 0.0001 | $H_2O$, ultrasonic | 1 M | 496 nM | 36717552 | 100 nM - 10 μM | |
| p-Hydroxybenzoate | 114-63-6 | 1.533 | 0.0001 | $H_2O$ | 100 mM | | | 100 μM - 10 mM | |
| p-Hydroxyphenylacetic acid | 156-38-7 | 44.59 | 0.0012 | DMSO | 500 mM | 657 μM | 10.3390/ijms150712861 | 100 μM - 10 mM | |
| Pimelic acid | 111-16-0 | 2.057 | 0.0056 | $H_2O$ | 500 mM | | | 100 μM - 10 mM | |
| Pipecolic acid | 535-75-1 | 0.7227 | 0.0121 | $H_2O$ | 200 mM | | | 100 μM - 10 mM | |
| Proline | 147-85-3 | 0.5769 | 0.0002 | $H_2O$ | 100 mM | 300 μM | 35786294 | 100 μM - 10 mM | |
| Propionic acid | 79-09-4 | 4.501 | 0.0003 | | 13.4 M | 20 mM | 34443546 | 1 mM - 100 mM | |
| Prostaglandin E2 | 363-24-6 | 2.262 | 0.0012 | DMSO | 10 mM | 1 μM | 33945793 | 100 nM - 10 μM | |
| Prostaglandin F2α | 4510-16-1 | 1.515 | < 0.0001 | | 3.25 M | | 34973337 | 100 μM - 10 mM | |
| p-Toluic acid | 99-94-5 | 0.8034 | < 0.0001 | DMSO | 500 mM | | | 100 μM - 10 mM | |
| Pyrazinamide | 98-96-4 | 1.999 | 0.0492 | DMSO | 400 mM | | | 100 μM - 10 mM | |
| Pyridine | 110-86-1 | 2.107 | 0.0337 | | 12.36 M | | | 100 μM - 10 mM | |
| Pyridoxamine | 85-87-0 | 0.4777 | 0.0005 | $H_2O$, ultrasonic | 20 mM | | | 10 μM - 1 mM | |
| Pyridoxine | 65-23-6 | 0.7111 | 0.0002 | DMSO | 500 mM | 5 μM | 32071304 | 1 μM - 100 μM | |
| Pyrophosphate | 7758-16-9 | 0.228 | < 0.0001 | $H_2O$ | 200 mM | 100 μM | 34142719 | 10 μM - 1 mM | |
| Pyruvate | 127-17-3 | 0.8154 | 0.0047 | $H_2O$ | 1 M | 3 mM | 37024456 | 100 μM - 20 mM | |
| Riboflavin | 83-88-5 | 5.531 | 0.0011 | DMSO | 10 mM | | | 10 μM - 1 mM | |
| Ribose 1-phosphate | 14075-00-4 | 0.5525 | < 0.0001 | $H_2O$ | 100 mM | | | 100 μM - 10 mM | |
| Ribose 5-phosphate | 4300-28-1 | 0.4588 | 0.0014 | $H_2O$ | 300 mM | 1 mM | 37597521 | 100 μM - 10 mM | |
| Ribulose 5-phosphate | 4151-19-3 | 0.3867 | < 0.0001 | PBS | 50 mM | | | 10 μM - 1 mM | |
| Sarcosine | 107-97-1 | 0.4633 | < 0.0001 | $H_2O$, ultrasonic | 1 M | | | 100 μM - 10 mM | |
| Sedoheptulose 7-phosphate | 2646-35-7 | 2.046 | 0.0001 | $H_2O$ | 100 mM | | | 100 μM - 10 mM | |
| Serine | 56-45-1 | 0.8445 | 0.0074 | $H_2O$ | 200 mM | 400 μM | 25903138 | 100 μM - 10 mM | |
| Serotonin | 50-67-9 | 0.4986 | < 0.0001 | $H_2O$, ultrasonic | 400 mM | 10 μM | 34281987 | 1 μM - 100 μM | |

**Extended Data Table 1 (cont.)**

| Name | CAS number | Fold changes (CR vs AL) | *P* value | Dissolution | Stock concent ration* | Concent ration reported | Reference (PMID) | Concentration used in screening assay | Concentration ranges able to activate AMPK |
|---|---|---|---|---|---|---|---|---|---|
| Sinapic acid | 530-59-6 | 1.958 | 0.015 | DMSO | 100 mM | 3.0 mM | 24053181 | 100 µM - 10 mM | |
| Sorbitol 6-phosphate | 108392-12-7 | 5.967 | 0.0337 | $H_2O$ | 50 mM | | | 10 µM - 1 mM | |
| Stearic acid | 57-11-4 | 1.107 | 0.0362 | DMSO | 50 mM | 100 µM | 25222131 | 10 µM - 1 mM | |
| Suberate | 505-48-6 | 1.64 | 0.0232 | DMSO | 500 mM | | | 100 µM - 10 mM | |
| Succinate | 150-90-3 | 1.374 | 0.006 | $H_2O$ | 100 mM | 2 mM | 34343908 | 100 µM - 10 mM | |
| Succinic acid | 110-15-6 | 1.876 | < 0.0001 | $H_2O$ | 200 mM | 500 µM | 37130518 | 100 µM - 10 mM | |
| Taurine | 107-35-7 | 1.075 | 0.0431 | $H_2O$ | 100 mM | 20 mM | 36402441 | 100 µM - 10 mM | |
| Tauro-α-MCA | 2260905-08-4 | 5.89 | 0.0011 | DMSO | 10 mM | | | 1 µM - 100 µM | |
| Tauro-ω-MCA | 2456348-84-6 | 1.494 | 0.0189 | DMSO | 10 mM | | | 1 µM - 100 µM | |
| TCA | 81-24-3 | 2.075 | 0.0049 | DMSO, ultrasonic | 100 mM | | | 100 µM - 10 mM | |
| TCDCA | 516-35-8 | 2.693 | 0.0074 | $H_2O$, ultrasonic | 200 mM | 100 µM | 27268718 | 10 µM - 1 mM | |
| TDCA | 110026-03-4 | 1.934 | 0.0226 | DMSO | 200 mM | 1 mM | 14739857 | 100 µM - 10 mM | |
| Thiamine | 67-03-8 | 2.315 | 0.0004 | $H_2O$ | 200 mM | 12 µM | 17463047 | 1 µM - 100 µM | |
| threo-β-Methylaspartic acid | 6667-60-3 | 63.19 | 0.0002 | $H_2O$ | 50 mM | | | 10 µM - 1 mM | |
| TLCA | 6042-32-6 | 5.397 | 0.0001 | DMSO | 100 mM | 10 µM | 10.1002/hep.510290227 | 1 µM - 100 µM | |
| trans-11-Eicosenoic acid | 62322-84-3 | 0.7225 | 0.0268 | DMSO | 200 mM | 100 µM | 26918025 | 10 µM - 1 mM | |
| trans-Oleic Acid | 112-79-8 | 1.743 | < 0.0001 | DMSO, ultrasonic | 10 mM | 800 µM | 29152653 | 10 µM - 1 mM | |
| Tridecanoic acid | 638-53-9 | 1.061 | 0.0031 | DMSO | 100 mM | | | 100 µM - 10 mM | |
| Trigonelline | 535-83-1 | 0.2812 | < 0.0001 | DMSO | 50 mM | 10 mM | 34560552 | 100 µM - 5 mM | |
| Trimesic acid | 554-95-0 | 1.269 | 0.0015 | DMSO | 200 mM | 5 µM | 22748770 | 1 µM - 100 µM | |
| Trimethylamine N-oxide | 1184-78-7 | 2.653 | < 0.0001 | $H_2O$, ultrasonic | 1 M | 300 µM | 35349832 | 10 µM - 1 mM | |
| Tryptamine | 61-54-1 | 1.746 | 0.0284 | DMSO | 500 mM | 1 mM | 16892423 | 100 µM - 10 mM | |
| Tryptophanamide | 20696-57-5 | 2.082 | < 0.0001 | | NA | | | | |
| Tyrosine methyl ester | 1080-06-4 | 1.956 | 0.0023 | DMSO | 500 mM | | | 100 µM - 10 mM | |
| UCA | 2955-27-3 | 174.5 | 0.0074 | DMSO | 10 mM | | | 10 µM - 1 mM | |
| UDCA | 128-13-2 | 47.65 | 0.0078 | DMSO | 200 mM | 100 µM | 22531947 | 10 µM - 1 mM | |
| UDP-N-acetylglucosamine | 91183-98-1 | 0.6926 | 0.0258 | $H_2O$ | 10 mM | | | 10 µM - 1 mM | |
| UMP | 58-97-9 | 0.4874 | 0.0005 | $H_2O$ | 200 mM | 50 mM | 31231750 | 100 µM - 20 mM | |
| Urea | 57-13-6 | 0.8618 | 0.0044 | $H_2O$ | 1 M | 20 mM | 30314315 | 1 mM - 100 mM | |
| Uric acid | 69-93-2 | 1.403 | 0.0035 | $H_2O$ | 1 mM | 594.8 µM | 24269076 | 1 µM - 100 µM | |
| Uridine | 58-96-8 | 0.4706 | 0.0011 | $H_2O$ | 200 mM | 200 µM | 31146110 | 10 µM - 1 mM | |
| UTP | 63-39-8 | 0.6327 | 0.0232 | $H_2O$ | 10 mM | 250 µM | 24905332 | 10 µM - 1 mM | |
| Valeric acid | 109-52-4 | 2.729 | < 0.0001 | $H_2O$ | 20 mM | 2 mM | 28683366 | 10 µM - 1 mM | |
| Valine | 72-18-4 | 0.7617 | 0.0237 | $H_2O$ | 100 mM | 1.7 mM | 36111572 | 100 µM - 10 mM | |
| Xanthine | 69-89-6 | 144.3 | < 0.0001 | DMSO | 20 mM | | | 10 µM - 1 mM | |
| Xanthosine | 146-80-5 | 10.72 | < 0.0001 | DMSO | 100 mM | | | 100 µM - 10 mM | |
| Xylose | 58-86-6 | 11.38 | < 0.0001 | $H_2O$ | 100 mM | | | 100 µM - 10 mM | |
| Xylulose 5-phosphate | 4212-65-1 | 0.4254 | 0.0006 | | NA | | | | |
| Z-Glycine | 1138-80-3 | 2.572 | 0.0127 | | 6.2 M | | | 100 µM - 10 mM | |
| α-KG | 328-50-7 | 1.488 | < 0.0001 | $H_2O$ | 200 mM | 5 mM | 33585273 | 100 µM - 20 mM | |
| α-MCA | 2393-58-0 | 149.1 | 0.0183 | DMSO | 100 mM | | | 10 µM - 1 mM | |
| α-Tocopherol | 10191-41-0 | 1.35 | < 0.0001 | $H_2O$, ultrasonic | 100 mM | 50 µM | 33098823 | 10 µM - 1 mM | |
| β-Alanine | 107-95-9 | 0.7052 | < 0.0001 | $H_2O$, ultrasonic | 100 mM | 100 mM | 24460609 | 100 µM - 10 mM | |
| β-MCA | 2393-59-1 | 40.38 | 0.0061 | DMSO | 200 mM | 100 µM | 33303970 | 10 µM - 1 mM | |
| ω-MCA | 6830-03-1 | 49.96 | 0.008 | DMSO | 40 mM | | | 1 µM - 100 µM | |

* NA represents an unavailable or untested metabolite.

**Extended Data Table 1 | A list of metabolites used in the screening assays and their effects in activating AMPK**.
The table presents the names, CAS numbers, fold changes during CR, *P* values, reported concentrations, references (PMID numbers), concentrations used in screening assays, and abilities to activate the AMPK of each compound.

**Extended Data Table 2 | Summary of lifespan and analysis in worms[a,b]**

| Genotypes/ treatments | Mean life span (days) Estimated life span ± s.e.m. | 95% confidence interval Lower bound | Upper bound | Median life span (days) Estimated life span ± s.e.m. | 95% confidence interval Lower bound | Upper bound | N[c] | N[d] | N[e] | P-value Vs saline control within each genotype (Mantel-CoX) |
|---|---|---|---|---|---|---|---|---|---|---|
| | | | | Fig. 5a | | | | | | |
| N2 | 22.373 ± 0.458 | 21.475 | 23.272 | 22.000 ±0.573 | 20.877 | 23.123 | 155 | 44 | 199 | N/A |
| N2 + LCA | 27.520 ± 0.684 | 26.180 | 28.860 | 30.000 ±1.029 | 27.983 | 32.017 | 145 | 55 | 200 | <0.001 |
| *aak-2* | 18.000 ± 0.432 | 17.152 | 18.848 | 18.000 ±0.630 | 16.766 | 19.234 | 148 | 52 | 200 | N/A |
| *aak-2* + LCA | 17.352 ± 0.417 | 16.534 | 18.169 | 18.000 ±0.632 | 16.761 | 19.239 | 169 | 31 | 200 | 0.259 |

**Extended Data Table 2 | Summary of healthspan and analysis in worms[a,b]**

| Genotypes/ treatments | Mean health span (hours) Estimated life span ± s.e.m. | 95% confidence interval Lower bound | Upper bound | Median health span (hours) Estimated life span ± s.e.m. | 95% confidence interval Lower bound | Upper bound | N[c] | N[d] | N[e] | P-value Vs saline control within each genotype (Mantel-CoX) |
|---|---|---|---|---|---|---|---|---|---|---|
| | | | Fig. 5d | | | | | | | |
| N2 | 9.845 ± 0.599 | 8.670 | 11.020 | 10.000 ± 1.095 | 7.854 | 12.146 | 49 | 11 | 60 | N/A |
| N2 + LCA | 13.583 ± 0.745 | 12.123 | 15.044 | 15.000 ± 0.866 | 13.303 | 16.697 | 51 | 9 | 60 | <0.001 |
| *aak-2* | 9.225 ± 0.492 | 8.260 | 10.189 | 10.000 ± 0.696 | 8.636 | 11.364 | 52 | 8 | 60 | N/A |
| *aak-2* + LCA | 9.492 ± 0.521 | 8.470 | 10.513 | 10.000 ± 0.883 | 8.269 | 11.731 | 52 | 8 | 60 | 0.470 |

[a]Independent repeats of each lifespan and healthspan experiment were performed. Data from representative experiments are shown.
[b]Health span data sets within each panel of this table were done in parallel and statistical analyses were done within the data set.
[c]Number of worms scored (death events).
[d]Number of worms censored.
[e]Total number of worms

**Extended Data Table 2 | Summary of lifespan and analysis in flies[a,b]**

| Genotypes/ treatments | Mean life span (days) | | | Median life span (days) | | | | | | P-value Vs saline control within each genotype (Mantel-CoX) |
|---|---|---|---|---|---|---|---|---|---|---|
| | | 95% confidence interval | | | 95% confidence interval | | | | | |
| | Estimated life span ± s.e.m. | Lower bound | Upper bound | Estimated life span ± s.e.m. | Lower bound | Upper bound | N[c] | N[d] | N[e] | |
| **Fig. 5b male** | | | | | | | | | | |
| *Act5C-GAL4* | 47.895 ± 1.010 | 45.915 | 49.875 | 49.000 ±1.816 | 45.441 | 52.559 | 200 | 0 | 200 | N/A |
| *Act5C-GAL4* + LCA | 52.655 ± 0.989 | 50.717 | 54.593 | 55.000 ±1.563 | 51.936 | 58.064 | 200 | 0 | 200 | <0.001 |
| *Act5C-GAL4* > *AMPKα* RNAi | 41.415 ± 0.802 | 39.844 | 42.986 | 43.000 ±1.010 | 41.020 | 44.980 | 200 | 0 | 200 | N/A |
| *Act5C-GAL4* > *AMPKα* RNAi+LCA | 41.450 ± 0.772 | 39.938 | 42.962 | 43.000 ±0.986 | 41.067 | 44.933 | 200 | 0 | 200 | 0.594 |
| **Fig. 5b female** | | | | | | | | | | |
| *Act5C-GAL4* | 52.120 ± 1.043 | 50.076 | 54.164 | 55.000 ±1.246 | 52.559 | 57.441 | 200 | 0 | 200 | N/A |
| *Act5C-GAL4* + LCA | 56.220 ± 1.016 | 54.229 | 58.211 | 61.000 ±1.098 | 58.847 | 63.153 | 200 | 0 | 200 | <0.001 |
| *Act5C-GAL4* > *AMPKα* RNAi | 42.095 ± 0.862 | 40.406 | 43.784 | 43.000 ±0.875 | 41.285 | 44.715 | 200 | 0 | 200 | N/A |
| *Act5C-GAL4* > *AMPKα* RNAi+LCA | 42.690 ± 0.890 | 40.946 | 44.434 | 43.000 ±1.247 | 40.555 | 45.445 | 200 | 0 | 200 | 0.580 |
| **Fig. 5b male** | | | | | | | | | | |
| AL | 34.920 ± 0.875 | 33.205 | 36.635 | 37.000 ±0.554 | 35.915 | 38.085 | 200 | 0 | 200 | N/A |
| AL+LCA | 43.220 ± 1.015 | 41.231 | 45.209 | 46.000 ±0.975 | 44.088 | 47.912 | 200 | 0 | 200 | <0.001 |
| CR | 44.206 ± 1.038 | 42.225 | 46.295 | 46.000 ±1.115 | 43.814 | 48.186 | 200 | 0 | 200 | N/A |
| CR+LCA | 45.015 ± 0.909 | 43.234 | 46.796 | 46.000 ±1.087 | 43.869 | 48.131 | 200 | 0 | 200 | 0.875 |
| **Fig. 5b female** | | | | | | | | | | |
| AL | 34.780 ± 0.930 | 32.958 | 36.602 | 34.000 ±1.354 | 31.347 | 36.653 | 200 | 0 | 200 | N/A |
| AL+LCA | 42.500 ± 1.090 | 40.363 | 44.637 | 43.000 ±1.248 | 40.555 | 45.445 | 200 | 0 | 200 | <0.001 |
| CR | 43.245 ± 1.184 | 40.925 | 45.565 | 43.000 ±1.243 | 40.564 | 45.436 | 200 | 0 | 200 | N/A |
| CR+LCA | 43.605 ± 1.231 | 41.192 | 46.018 | 43.000 ±1.719 | 39.630 | 46.370 | 200 | 0 | 200 | 0.784 |
| **Extended Data Fig. 5e male** | | | | | | | | | | |
| *w*$^{1118}$ | 44.090 ± 0.814 | 42.494 | 45.686 | 47.000 ±0.653 | 45.720 | 48.280 | 200 | 0 | 200 | N/A |
| *w*$^{1118}$ + LCA | 50.660 ± 0.761 | 49.169 | 52.151 | 51.000 ±0.611 | 49.802 | 52.198 | 200 | 0 | 200 | <0.001 |
| **Extended Data Fig. 5e female** | | | | | | | | | | |
| *w*$^{1118}$ | 50.465 ± 0.886 | 48.728 | 52.202 | 51.000 ±1.335 | 48.383 | 53.617 | 200 | 0 | 200 | N/A |
| *w*$^{1118}$ + LCA | 56.435 ± 0.921 | 54.630 | 58.240 | 57.000 ±1.081 | 54.881 | 59.119 | 200 | 0 | 200 | <0.001 |

[a]Independent repeats of each lifespan experiment were performed. Data from representative experiments are shown.
[b]Health span data sets within each panel of this table were done in parallel and statistical analyses was done within the data set.
[c]Number of flies scored (death events).
[d]Number of flies censored.
[e]Total number of flies.

**Extended Data Table 2 | Summary of healthspan and analysis in flies[a,b]**

| Genotypes/ treatments | Mean health span (hours) | | | Median health span (hours) | | | | | | |
|---|---|---|---|---|---|---|---|---|---|---|
| | Estimated life span ± s.e.m. | 95% confidence interval | | Estimated life span ± s.e.m. | 95% confidence interval | | | | | P-value Vs saline control within each genotype (Mantel-CoX) |
| | | Lower bound | Upper bound | | Lower bound | Upper bound | N[c] | N[d] | N[e] | |
| **Fig. 5e** | | | | | | | | | | |
| *Act5C-GAL4* | 40.242 ± 0.926 | 38.426 | 42.057 | 44.500 ± 0.657 | 43.213 | 45.787 | 120 | 0 | 120 | N/A |
| *Act5C-GAL4* + LCA | 45.746 ± 0.893 | 43.995 | 47.496 | 49.500 ± 0.597 | 48.331 | 50.669 | 120 | 0 | 120 | <0.001 |
| *Act5C-GAL4* > *AMPKα* RNAi | 34.108 ± 0.778 | 32.583 | 35.633 | 35.000 ± 0.829 | 33.375 | 36.625 | 120 | 0 | 120 | N/A |
| *Act5C-GAL4* > *AMPKα* RNAi+LCA | 34.142 ± 0.655 | 32.858 | 35.426 | 35.000 ± 0.777 | 33.477 | 36.523 | 120 | 0 | 120 | 0.387 |
| **Fig. 5f** | | | | | | | | | | |
| *Act5C-GAL4* | 32.254 ± 0.638 | 31.004 | 33.504 | 31.500 ± 0.542 | 30.439 | 32.561 | 120 | 0 | 120 | N/A |
| *Act5C-GAL4* + LCA | 37.008 ± 0.734 | 35.571 | 38.445 | 36.500 ± 0.772 | 34.988 | 38.012 | 120 | 0 | 120 | <0.001 |
| *Act5C-GAL4* > *AMPKα* RNAi | 21.483 ± 0.569 | 20.367 | 22.599 | 19.000 ± 0.625 | 17.776 | 20.224 | 120 | 0 | 120 | N/A |
| *Act5C-GAL4* > *AMPKα* RNAi+LCA | 21.267 ± 0.560 | 20.168 | 22.365 | 19.000 ± 0.841 | 17.352 | 20.648 | 120 | 0 | 120 | 0.679 |
| **Fig. 5g** | | | | | | | | | | |
| *Act5C-GAL4* | 22.775 ± 0.748 | 21.309 | 24.241 | 21.000 ± 1.217 | 18.615 | 23.385 | 120 | 0 | 120 | N/A |
| *Act5C-GAL4* + LCA | 27.675 ± 0.813 | 26.082 | 29.268 | 25.000 ± 0.730 | 23.569 | 26.431 | 120 | 0 | 120 | <0.001 |
| *Act5C-GAL4* > *AMPKα* RNAi | 12.446 ± 0.403 | 11.656 | 13.235 | 13.000 ± 0.342 | 12.329 | 13.671 | 120 | 0 | 120 | N/A |
| *Act5C-GAL4* > *AMPKα* RNAi+LCA | 12.583 ± 0.424 | 11.753 | 13.414 | 13.000 ± 0.393 | 12.230 | 13.770 | 120 | 0 | 120 | 0.774 |
| **Fig. 5h** | | | | | | | | | | |
| *Act5C-GAL4* | 4.817 ± 0.233 | 4.360 | 5.274 | 4.000 ± 0.321 | 3.372 | 4.628 | 120 | 0 | 120 | N/A |
| *Act5C-GAL4* + LCA | 6.150 ± 0.285 | 5.591 | 6.709 | 6.000 ± 0.365 | 5.284 | 6.716 | 120 | 0 | 120 | 0.001 |
| *Act5C-GAL4* > *AMPKα* RNAi | 3.325 ± 0.130 | 3.071 | 3.579 | 3.000 ± 0.083 | 2.838 | 3.162 | 120 | 0 | 120 | N/A |
| *Act5C-GAL4* > *AMPKα* RNAi+LCA | 3.408 ± 0.140 | 3.133 | 3.683 | 3.000 ± 0.094 | 2.816 | 3.184 | 120 | 0 | 120 | 0.657 |
| **Fig. 5i** | | | | | | | | | | |
| *Act5C-GAL4* | 33.242 ± 1.104 | 31.078 | 35.405 | 31.500 ± 1.557 | 28.448 | 34.552 | 120 | 0 | 120 | N/A |
| *Act5C-GAL4* + LCA | 37.800 ± 1.304 | 35.243 | 40.357 | 33.500 ± 1.823 | 29.926 | 37.074 | 120 | 0 | 120 | 0.003 |
| *Act5C-GAL4* > *AMPKα* RNAi | 19.425 ± 0.673 | 18.106 | 20.744 | 17.000 ± 0.453 | 16.111 | 17.889 | 120 | 0 | 120 | N/A |
| *Act5C-GAL4* > *AMPKα* RNAi+LCA | 19.717 ± 0.696 | 18.352 | 21.082 | 17.000 ± 0.753 | 15.524 | 18.476 | 120 | 0 | 120 | 0.753 |

[a]Independent repeats of each healthspan experiment were performed. Data from representative experiments are shown.
[b]Health span data sets within each panel of this table were done in parallel and statistical analyses was done within the data set.
[c]Number of flies scored (death events).
[d]Number of flies censored.
[e]Total number of flies.

**Extended Data Table 2 | Summary of healthspan and analysis in flies (cont.)[a,b]**

| Genotypes/ treatments | Mean health span (hours) | | | Median health span (hours) | | | | | | |
|---|---|---|---|---|---|---|---|---|---|---|
| | Estimated life span ± s.e.m. | 95% confidence interval | | Estimated life span ± s.e.m. | 95% confidence interval | | | | | P-value Vs saline control within each genotype (Mantel-CoX) |
| | | Lower bound | Upper bound | | Lower bound | Upper bound | N[c] | N[d] | N[e] | |
| **Extended Data Fig. 5f** | | | | | | | | | | |
| *W[1118]* | 29.083 ± 1.046 | 27.033 | 31.134 | 27.000 ± 0.966 | 25.106 | 28.894 | 120 | 0 | 120 | N/A |
| *W[1118]*+LCA | 35.725 ± 1.141 | 33.488 | 37.962 | 36.000 ± 2.053 | 31.976 | 40.024 | 120 | 0 | 120 | <0.001 |
| **Extended Data Fig. 5g** | | | | | | | | | | |
| *W[1118]* | 24.808 ± 0.736 | 23.367 | 26.250 | 24.000 ± 1.043 | 21.955 | 26.045 | 120 | 0 | 120 | N/A |
| *W[1118]*+LCA | 28.917 ± 0.777 | 27.393 | 30.440 | 28.000 ± 0.995 | 26.051 | 29.949 | 120 | 0 | 120 | <0.001 |
| **Extended Data Fig. 5h** | | | | | | | | | | |
| *W[1118]* | 19.733 ± 0.717 | 18.328 | 21.139 | 19.000 ± 0.728 | 17.574 | 20.426 | 120 | 0 | 120 | N/A |
| *W[1118]*+LCA | 25.267 ± 0.848 | 23.605 | 26.928 | 23.000 ± 0.876 | 21.283 | 24.717 | 120 | 0 | 120 | <0.001 |
| **Extended Data Fig. 5i** | | | | | | | | | | |
| *W[1118]* | 5.317 ± 0.242 | 4.842 | 5.791 | 5.000 ± 0.264 | 4.483 | 5.517 | 120 | 0 | 120 | N/A |
| *W[1118]*+LCA | 6.517 ± 0.285 | 5.959 | 7.074 | 6.000 ± 0.378 | 5.260 | 6.740 | 120 | 0 | 120 | 0.003 |
| **Extended Data Fig. 5j** | | | | | | | | | | |
| *W[1118]* | 24.183 ± 0.873 | 22.473 | 25.894 | 22.000 ± 0.730 | 20.569 | 23.431 | 120 | 0 | 120 | N/A |
| *W[1118]*+LCA | 29.604 ± 1.238 | 27.177 | 32.031 | 27.000 ± 1.362 | 24.330 | 29.670 | 120 | 0 | 120 | <0.001 |

[a]Independent repeats of each healthspan experiment were performed. Data from representative experiments are shown.
[b]Health span data sets within each panel of this table were done in parallel and statistical analyses was done within the data set.
[c]Number of worms or flies scored (death events).
[d]Number of worms or flies censored.
[e]Total number of flies.
